# Neural correlates of pure presence

**DOI:** 10.1101/2024.04.18.590081

**Authors:** Melanie Boly, Richard Smith, Giulietta Vigueras Borrego, Joan Paul Pozuelos, Tariq Alauddin, Peter Malinowski, Giulio Tononi

**Affiliations:** Department of Neurology, University of Wisconsin-Madison, USA; Department of Psychiatry, University of Wisconsin-Madison, USA; Neuromindset, University of Granada, Spain; School of Psychology, Liverpool John Moores University, UK

**Author notes:** Corresponding authors: Melanie Boly, Giulio Tononi. **Email:**. **Author Contributions:** MB, RS, JPP, PM and GT designed research, MB, RS, GVB, JPP and TA performed research, MB, RS, GVB, JPP, TA, PM and GT analyzed data, MB, PM and GT supervised research, MB, RS, GVB, JPP, TA, PM and GT wrote the paper.

**Keywords:** meditation, consciousness, high-density EEG

## Abstract

Pure presence (PP) is described in several meditative traditions as an experience of a vast, vivid space devoid of perceptual objects, thoughts, and self. Integrated information theory (IIT) predicts that such vivid experiences may occur when the cortical substrate of consciousness is virtually silent. To test this, we analyzed high-quality 256-electrode high-density electroencephalography (hdEEG) from twenty-two long-term Vajrayana and Zen meditators who reported reaching PP during a week-long retreat. Because neural activity typically increases gamma power, we predicted PP would show widespread gamma reductions. Across both traditions, PP was associated with broadband power decrease compared to within-meditation mind-wandering, most consistent in the gamma range (30– 45 Hz). Source reconstruction revealed widespread gamma decreases, strongest in posteromedial cortex. PP gamma power was lower than in all other control states, including watching or imagining a movie, active thinking, and open-monitoring. PP delta power (1–4Hz) was also markedly reduced compared to dreamless sleep. Meditative states resembling PP—with minimal perceptual contents or accompanied by bliss— showed similar signatures. Overall, PP appears to be a state of vivid consciousness during which the cortex is highly awake (low delta) yet widely quiescent (low gamma), in line with IIT’s predictions.

## Introduction

It is generally assumed that consciousness requires neural activity. Neuronal firing would be needed to support specific computations, the “processing” of information, or the performance of certain cognitive functions. For example, consciousness might depend on recurrent processing within posterior cortex (1), on interactions between prefrontal and posterior cortex (2), on global information broadcasting (3), on top-down, predictive processing (4–6), and so on (7).

In contrast, integrated information theory (IIT, presented in detail in (8) and in an online wiki (9)) argues that an experience corresponds to a “cause–effect structure”, rather than to a computation, a function, or a process. The cause–effect structure expresses the powers of the units of a neural substrate in their current state to “take and make a difference” within it. It is composed of all the intrinsic causes and effects specified over the substrate by subsets of its units (causal distinctions), as well as by the overlaps among those causes and effects (causal relations). Intrinsic causes and effects themselves capture how the current state of subsets of substrate units affects the probability of past and future states. According to the theory, the cause–effect structure specified by a substrate in its current state accounts in full for both the quality of the experience and its quantity, with no additional ingredients.

A counterintuitive prediction of the theory is that a substrate with the appropriate anatomical organization—a densely connected, hierarchical lattice as found primarily in posterior-central cortical areas—should support an experience even when its units are in a state of inactivity (or near inactivity). This is because inactive units, similar to active ones, increase the probability of specific past and future states. Therefore, a virtually silent network specifies a cause-effect structure that can be similarly rich as one specified by a network with subsets of active units. Moreover, IIT shows that a lattice-like substrate supports a cause-effect structure that can account for the feeling of extendedness that characterizes spatial experiences (10). Accordingly, IIT predicts that as long as its causal powers are intact, a lattice-like substrate in a state of near-inactivity should specify a cause–effect structure corresponding to a vivid experience of pure “extendedness” (11). On the other hand, if the substrate’s causal powers were disrupted, notably by the bistability typical of early NREM sleep (12), consciousness should be lost despite ongoing neural activity (13).

While it is currently not feasible to test this prediction in the human brain at the level of local populations of neurons, it is possible to assess its plausibility by recording gamma power with high-density electroencephalography (hdEEG), to the extent that gamma power can serve as a proxy for neuronal activation (14–16). To begin addressing these predictions experimentally, we focused on states of “pure presence” (PP), also known as “pure consciousness” (17, 18). Such states are acknowledged by several spiritual traditions (17, 18) and are described as experiences of vast, vivid luminosity, devoid of thoughts, objects, particular sensory features, the perception of a self, or any distinction between subject and object (non-duality). For example, the *Tibetan Book of the Dead* speaks of “a lucid clarity without anyone being there who is the observer…” where “only a naked manifest awareness is present” … “a vast luminous expanse, clarity inseparable from emptiness” (19). States of “pure being” with no self, no thoughts, and no perceptual contents have also been reported after severe trauma (20) and certain phases of sleep (21). It is also helpful to think of such states as an extreme on a continuum of cortical activation: different contents of experience—whether faces or objects (22, 23), simple sensations (24), thoughts (25), or self-related ideation (26, 27)—are typically accompanied by neuronal activations in specific neuronal populations, surrounded by widespread quiescence or deactivation. One would thus expect that during quiet, thoughtless, and selfless moments, in the absence of exogenous or endogenous stimuli, the entire neural substrate of consciousness should be in a largely quiescent state.

In this work, we employed high-density electroencephalography (hdEEG, 256 channels) to record brain activity from experienced meditators who could reliably reach states of PP. Given that hdEEG gamma power (25–45 Hz) tracks locally increased neuronal activity when experiencing different conscious contents (22, 25), we predicted that PP states should be associated with a widespread decrease in gamma power. By contrast, because increased delta activity (0.5–4 Hz) is a hallmark of a breakdown of causal interactions typically associated with states of unconsciousness (12, 22), we predicted that delta power should stay the same or even decrease.

This study sought to obtain a clear picture of the neural correlates of PP by minimizing potential confounding factors in several ways. First, we relied on long-term meditators (LTMs) because of their training in reaching such states. Moreover, we provided an explicit characterization of the target state for this study: no self, no thoughts, no perceptual objects or sensory features, non-duality, and instead a pure sense of spatial extendedness. Preliminary sessions were devoted to clarifying the phenomenal properties of the target PP state, disentangling them from the specific jargon or goals specific to a particular tradition. For example, some descriptions from LTMs may mention that “the visual perception of space disappears” during PP (18), but it turns out that it is the “external” space—the world—that vanishes, with the landmarks that delimit it, rather than the “internal” space—the feeling of extendedness of the experience as such. Second, we recruited LTMs from two different schools—the Vajrayana tradition, in which the “dissolving phase” corresponding to PP is a desired target, and the Zen tradition, in which PP-like states are encountered on the way to other target states. Third, we took advantage of week-long retreats to acclimate the subjects to the experimental setup and facilitate the attainment of PP. Fourth, we employed both a delayed-report paradigm and a no-report paradigm. As confirmed by individual interviews (see also (18)), PP is incompatible with immediate report. However, subjects can be trained to report its occurrence after a delay upon exiting the state (sometimes characterized as an “afterglow” (22)). Moreover, Vajrayana meditators were also recorded in a no-report paradigm while undergoing a prescribed sequence of meditation states (with instructions provided before each phase). Sixth, we employed a number of control conditions, both within meditation (mind-wandering and other meditative states with rich perceptual contents) and outside meditation (resting with eyes open or eyes closed, movie watching, imagining, performing a mental calculation). Seventh, we considered conditions of near-PP in which meditators reported minimal perceptual contents and PP accompanied by a feeling of bliss, to assess the robustness of the findings. Eight, we compared PP with dreamless sleep to assess the expected differences between PP and loss of consciousness.

As to the recordings, because PP states may be hard to maintain, we focused the analysis of hdEEG on a five-second epoch of recording occurring just before the report in the delayed-report paradigm or just after the start of the dissolving phase in the no-report paradigm. This approach maximized our chances of identifying the neural correlates of PP in both cases. Crucially, we employed a state-of-the-art procedure to remove any non-neural contaminant from the EEG signal (28). This is essential to ensure that any differences between PP and other states are due to neural changes rather than to changes in muscle activity, eye movements, sweat, or other factors (29). The use of 256 channels is especially important in this regard as it offers great sensitivity in revealing extraneous components (30, 31). Furthermore, we considered all frequency bands below 45Hz and all channels, in addition to performing source localization, to avoid missing localized activations.

## Results

### LTM demographics and retreat procedure

Thirty-eight LTMs from the Vajrayana Karma Kagyu and Zen traditions of Buddhism were recorded during week-long retreats (Fig. 1). Inclusion criteria included at least 3000 hours of lifetime meditation experience, current meditation practice of several times a week, and previous meditation experience in a retreat setting. Exclusion criteria included a history of neurological disorder or current substance abuse (see Methods and Supplementary Material for details). During Vajrayana 8^th^ Karmapa Guru Yoga or Zen Shikantaza practices, LTMs were instructed to signal with a mouse click when they had just emerged from a state of PP—defined as a “phenomenal state without thought, self, or perceptual contents.” Additionally, they were asked to signal PP-like states with minimal perceptual content (PP-mpc), and states of mind-wandering (MW) occurring during their practice. Zen LTMs were also instructed to report tradition-specific states with “minimal thoughts” or “no thoughts” with preserved perceptual content, and non-dual states where “the self–other distinction disappears while the whole world is present”. An ‘unconscious’ rating was also available in the Zen scale, but was never used by the participants throughout the retreats. Vajrayana LTMs were also asked to report tradition-specific states of “bliss-awareness”—that is, PP-like states with “a feeling of bliss arising from mind itself” (PP-bliss). A range of other controls such as eyes open, eyes closed, watching then imagining a movie, active thinking (counting backwards), and open-monitoring were recorded each day. In Vajrayana LTMs, PP states were also acquired along with other content-rich meditative states by recording hdEEG during different phases of a pre-recorded, timed 16^th^ Karmapa Guru Yoga practice. When feasible, LTMs who reported PP states during retreats were brought to the sleep laboratory for overnight hdEEG recordings.

**Fig. 1.**
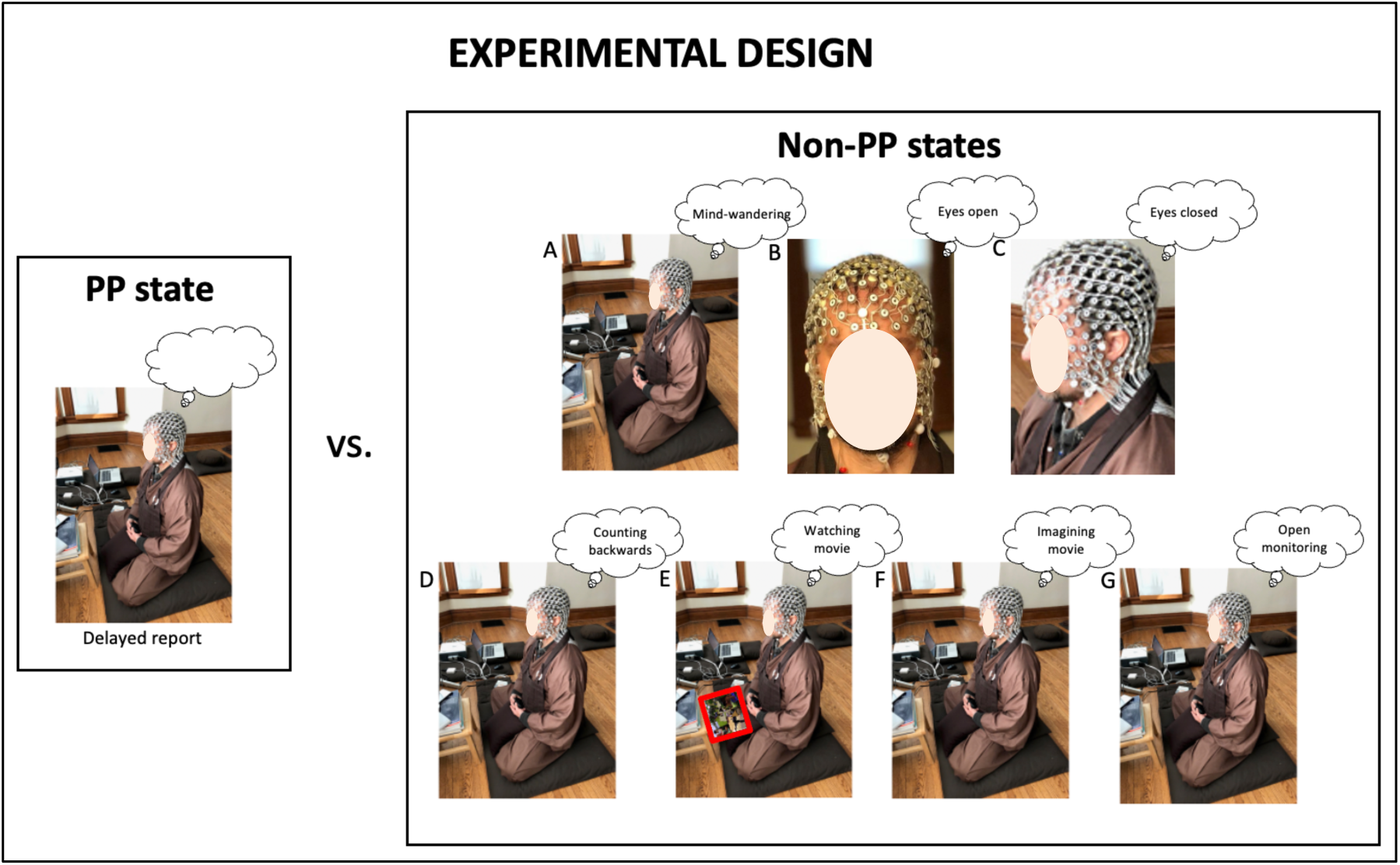
Experimental design. 256-electrode hdEEG was recorded during PP and compared to a variety of non-PP states: within-meditation mind-wandering (A), resting eyes open (B) and eyes closed (C), watching a movie (D), imagining the same movie (E), active thinking (counting backwards, F), open-monitoring (G).

Two phenomenological interviews (one from a Vajrayana practitioner, one from a Zen practitioner) in Supplementary Material describe typical phenomenology of PP states. PP is there described as an experience of vast, luminous expanse, without self, thoughts or sensory percepts.

HdEEG data from days 5–7 of the retreats was kept for analysis. Preprocessing included careful visual channel and epoch selection to reduce artifacts, followed by an iterative adaptive independent component analysis (EEGLab AMICA) procedure that has been optimized over the last 5 years (Suppl. Fig. 1). In brief, the time courses of the first 20 components of AMICA decomposition (containing most of the signal variance) were systematically inspected; artifact components were identified both through their focal aspect on 256-electrode topography (unlike brain components which are spatially filtered by the skull and are thus spatially smooth across >10 centimeters) and their time courses (prominent mix of low- and high-frequency signals rather than 1/f combined with oscillatory activities). Outlier epochs where ICA failed to separate brain from artifact components (reflected in mixed artifact and brain signals in several component time courses) were discarded; AMICA was then re-run on remaining time courses and component separation was re-evaluated. The only hdEEG datasets kept for further analysis were those in which a consistent separation of brain activity and artifacts within the first 20 components could be achieved (often requiring several AMICA iterations). Within these datasets, only brain activity components showing a spatially smooth topography and with consistently clean 1/f-oscillatory time courses (Suppl. Fig. 1) were kept for further analysis. After preprocessing, hdEEG data quality was once again verified through the assessment of individual hdEEG topographies. This allowed us to distinguish smooth brain activity from more discrete extra-cranial artifacts (see Suppl. Fig. 1, and examples of the many artifactual components identified by AMICA in a representative subject in Suppl. Fig. 2) in the delta (1–4 Hz), theta (4–8 Hz), alpha (8–12 Hz), beta (15–30 Hz), and gamma frequency bands (30–50 Hz for the Zen cohort recorded in the USA, and 30–45 Hz for the Vajrayana cohort recorded both in the USA and Europe owing to differences in the frequency of line noise). Power band topographies were also computed for source-reconstructed hdEEG time courses. This procedure yielded clean recordings of PP and related meditative states in 22 out of the 38 LTMs studied (14 Vajrayana LTMs and 8 Zen LTMs): Thirteen of the twenty-four Vajrayana LTMs reported states of PP during the last day of 8^th^ Karmapa practice (seven of which had high-quality recordings during PP events) and eight of the fourteen Zen meditators achieved states of PP during Shikantaza practice (all of which had high-quality recordings). Additionally, thirteen out of the twenty-four Vajrayana LTMs reported states of PP-bliss during the last day of the 8^th^ Karmapa practice (nine of them having high-quality recordings during that state), and eleven out of twenty-four had consistently high-quality recordings during the 16^th^ Karmapa practice that day (six of them overlapped with LTMs reaching PP during the 8^th^ Karmapa practice). Demographic data for the 14 Vajrayana LTMs included in the present analyses were as follows: avg 13 consecutive years of meditation (± sd 5): avg 6 practice days per week (± sd 1); avg practice 55 min/day (± sd 27); avg 5427 estimated total lifetime meditation hours (± sd 1900); avg 37.93 years old (± 6.16); 7 males and 7 females. Demographic data for the 8 Zen LTMs included in the present analyses were as follows: avg 14 consecutive years of meditation (± sd 6); avg 6 practice days per week (± sd 1); avg practice 48 min/day (± sd 14); avg 8004 estimated total lifetime meditation hours (± sd 4174); avg 41.3 years old (± sd 17.15); 5 males and 3 females. Clean sleep recordings with serial awakenings were also obtained in 2 Zen LTMs and a separate sample of 13 Vipassana LTMs (with avg 4310 practice hours [± sd 2481] and avg 49 min of daily practice [± sd 18]) for comparison with PP states.

### PP compared to MW: delayed report within meditation practice

In order to minimize confounds, our main analysis consisted in comparing neural signatures of PP states to MW captured during the same meditation session. Statistical analysis was performed using a paired t-test between PP and MW at the within-subject level, and thresholded at cluster-based p<0.05 corrected for multiple comparisons using statistical nonparametric mapping (SNPM, see Methods). Figure 2 shows the result of this main analysis. Compared to MW, PP states were characterized by a decrease in EEG power in theta to gamma frequency bands, most consistent and widespread in the gamma range. Similar results for signatures of PP states were obtained across the two different meditation traditions and at the single-subject level (Suppl. Figs. 3–6).

**Fig. 2.**
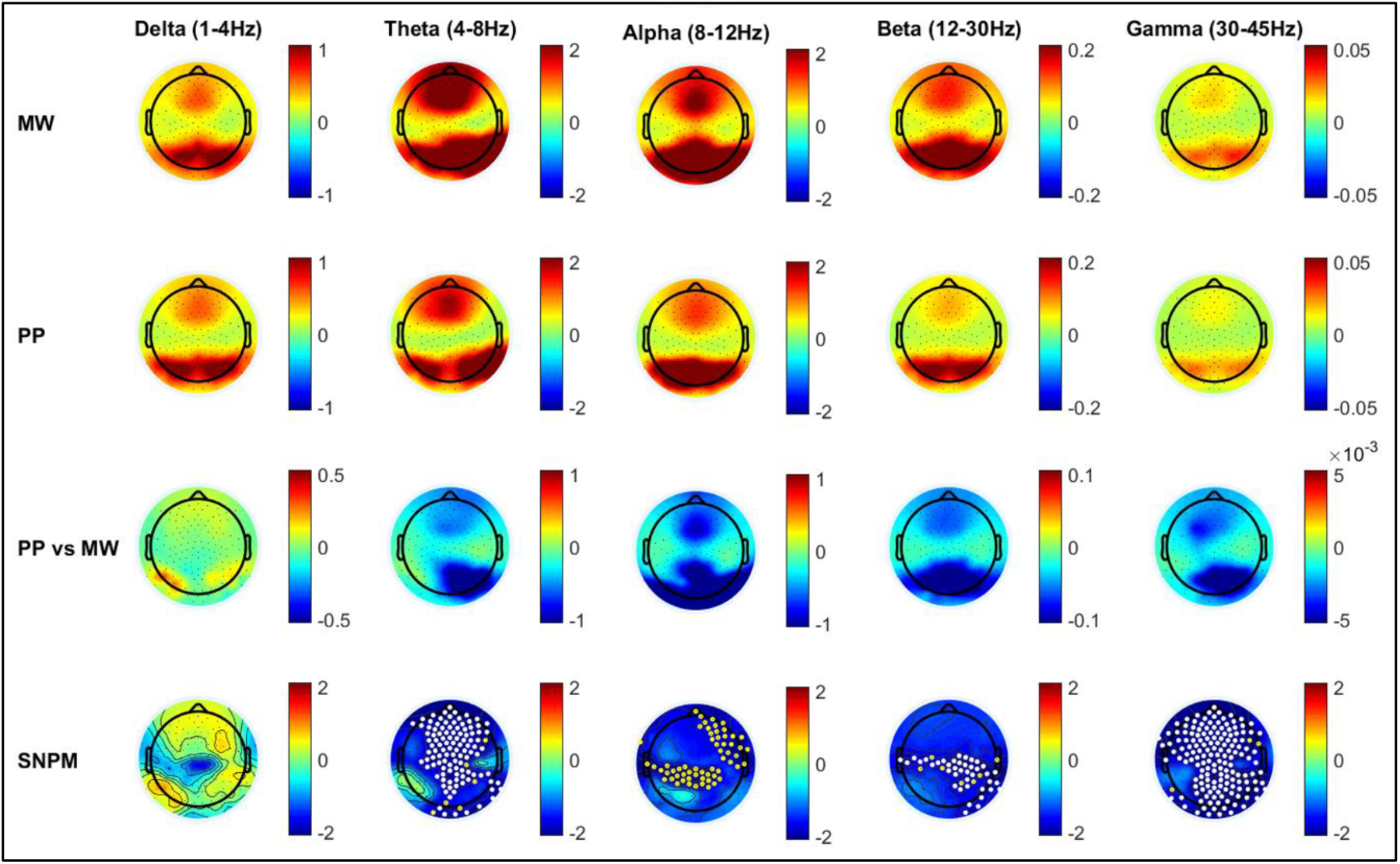
Neural correlates of pure presence (PP) compared to within-meditation mind-wandering MW in 15 long-term meditators (LTMs; 7 Vajrayana and 8 Zen) in the delta to gamma frequency ranges. Data obtained during Vajrayana 8^th^ Karmapa Guru Yoga and Zen Shikantaza practices. First row: average group topography for MW. Second row: average group topography for PP. Third row: average group difference for PP compared to MW (within-subject subtraction of MW from PP). Last row: statistical non-parametric mapping (SNPM) statistics for group difference between PP and MW. Colors represent EEG power (first to third rows) and T values (last row). White dots represent channels surviving p<0.05 corrected for multiple comparisons using SNPM; yellow dots represent channels surviving uncorrected p<0.05 values.

Next, we performed source reconstruction analysis of PP signatures (compared to MW), focusing on the gamma band (results for delta to beta bands are shown in Suppl. Fig. 7). Source reconstruction employed a template head model and minimum norm prior as implemented in Brainstorm (see Methods). Again a paired t-test was performed between PP and MW, and statistics were p<0.05 corrected for multiple comparisons across vertices using false-discovery rate (FDR). Fig. 3 displays the results of this analysis. In contrast to MW, PP was characterized by a widespread decrease in cortical gamma activity, most prominently in posteromedial areas.

**Fig 3.**
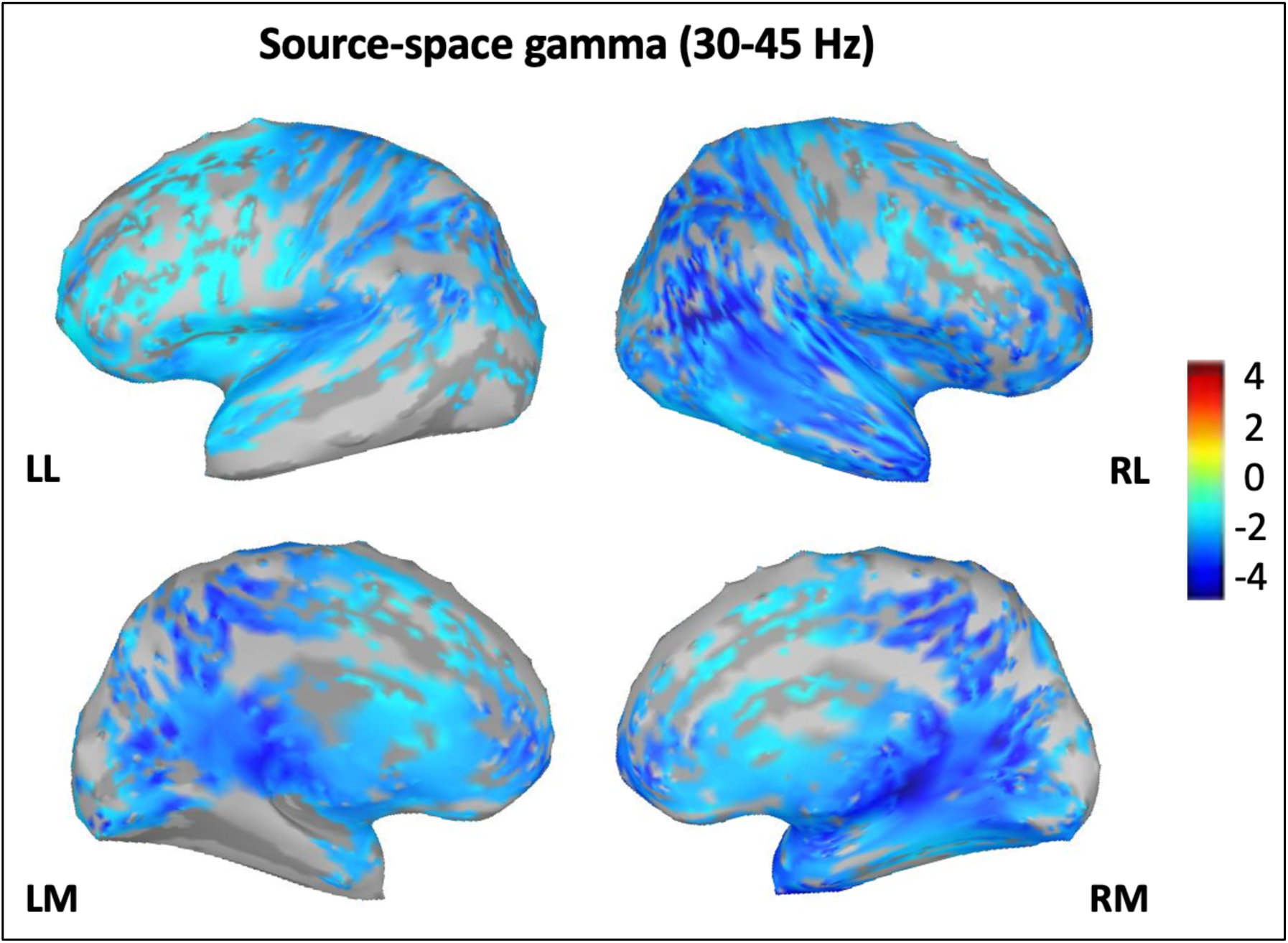
Source reconstruction results for gamma decrease in PP vs MW, corrected at FDR p<0.05. LL: left lateral; RL: right lateral; LM: left medial; RM: right medial. Colors represent T value for changes in source-level gamma power during PP compared to MW in the same 15 LTMs displayed in Figure 1. Darker blue colors, representing stronger decreases, most prominently in posteromedial areas.

In four Zen participants, MW occurred both before and after PP. Exploratory spectral analysis revealed similar decrease in gamma power during PP compared to MW occurring before vs after it (although these exploratory results did not survive correction for multiple comparisons, Suppl. Fig. 8).

Lempel-Ziv complexity values were not significantly different between the PP and the MW conditions (P = 0.11) although the PP condition did have a higher average complexity score (155.50 ± 40.15) compared to the MW condition (146.38 ± 37.79, Suppl. Fig. 9).

### PP vs. open-monitoring

Next, we compared PP to a different, well-characterized meditative state, open-monitoring (OM), which is characterized by refraining from engaging in thinking but preserved perceptual content. An OM meditation session was recorded each day before the PP meditation sessions. As shown in Fig. 4 (N = 15 LTMs), OM states were associated with an *increase* in delta, beta and gamma activity compared to both MW and PP.

**Fig. 4.**
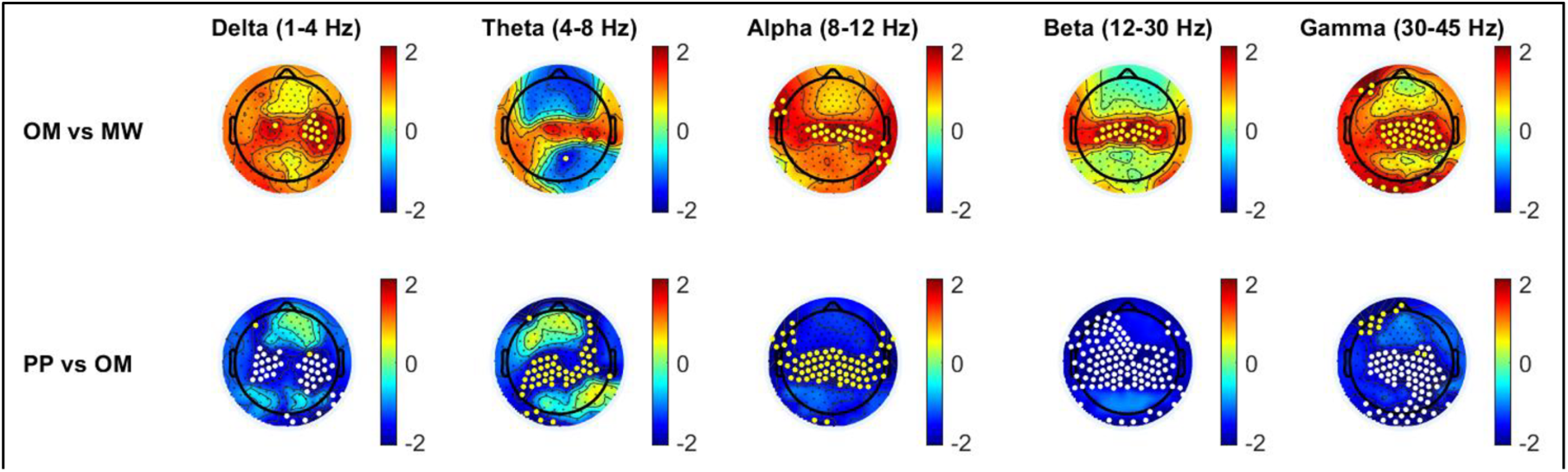
Comparison of open-monitoring (OM) to MW (first row) and PP (second row). PP data are from the same 15 LTMs displayed in Figs. 2–3, OM data from 13/15 of the same LTMs and 2 additional Vajrayana LTMs. Colors represent EEG power T value); white dots represent channels surviving SNPM-corrected p values<0.05; yellow dots represent channels surviving uncorrected p<0.05.

### PP and control states outside of meditation practice

To further assess the specificity of the neural signatures of PP, we compared this state to a set of other control conditions collected outside of meditation. Such control conditions included eyes open, eyes closed, movie watching and re-imagining, and active thinking (counting backwards) for 2–3 minutes each (see Methods). These control conditions were recorded each day of the retreat before meditation sessions. Fig. 5 displays comparison between neural signatures of PP states and those of each of these controls recorded the same day. Results show a consistent decrease in delta and gamma power during PP compared to other states, which survived SNPM correction for multiple comparisons in all states except for active thinking (uncorrected p<0.05 in both delta and gamma ranges, with PP still showing significant decrease in activity in the beta range surviving SNPM).

**Fig. 5.**
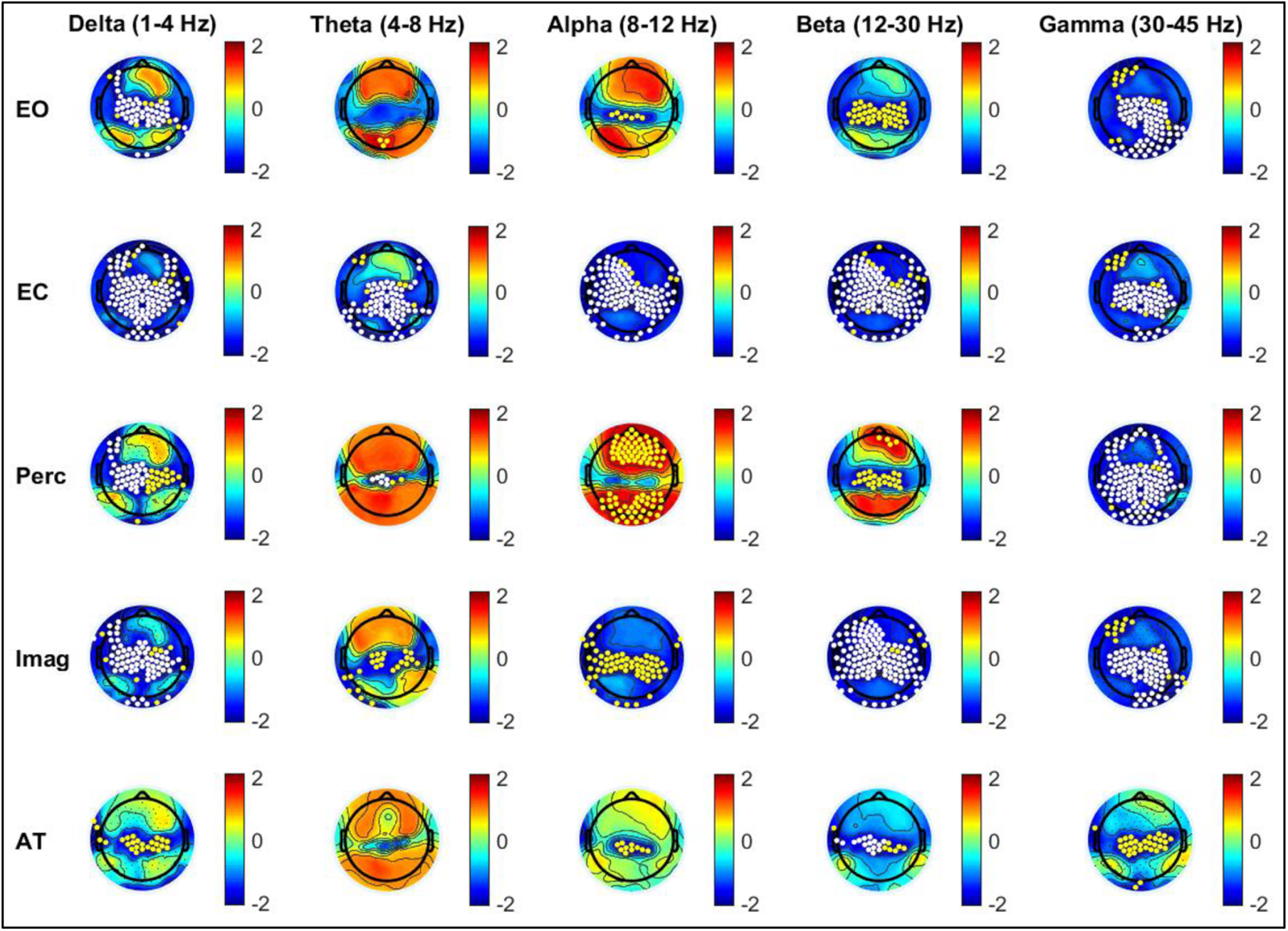
PP shows consistent delta, beta and gamma power decreases compared to other control states outside of meditation (same LTM data as displayed in Fig. 4). EO: resting with eyes open; EC: resting with eyes closed; Perc: watching a movie; Imag: imagining the same movie; AT: active thinking (counting backwards). Colors represent T values; white dots represent channels surviving SNPM-corrected p values<0.05; yellow dots represent channels surviving uncorrected p<0.05.

**Fig. 6.**
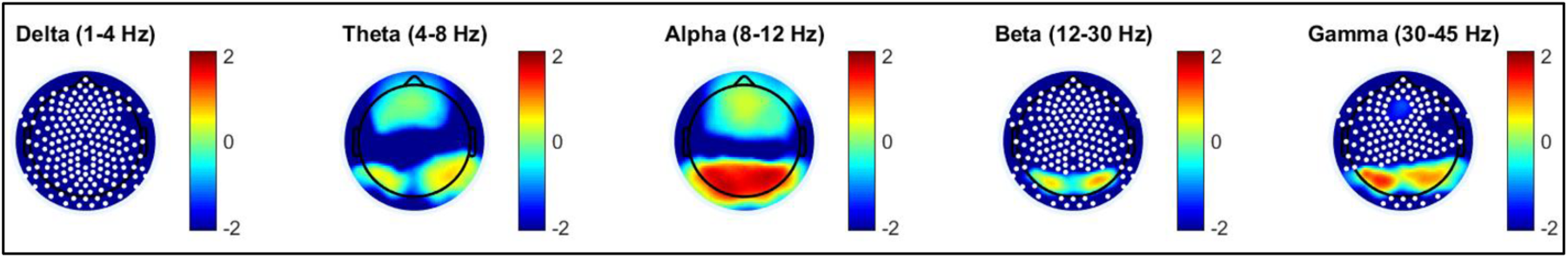
Compared to dreamless sleep, PP shows widespread decreases in EEG power, most consistent in delta, beta, and gamma ranges (PP data from the same 15 LTMs displayed in Figs. 2–4; sleep data from 2 Zen and 13 age-matched Vipassana LTMs). Colors represent T values; white dots represent channels surviving SNPM-corrected p values<0.05.

### PP vs. dreamless sleep

Finally, we compared neural signatures of PP in the same 15 LTMs described above to those of epochs of dreamless non-rapid eye movement (NREM) sleep obtained in 15 LTMs (2 Zen LTMs and 13 other age-matched LTMs from the separate Vipassana cohort), in which serial awakenings with experience sampling were performed (see Methods). Group results (displayed in Fig. 6) revealed that PP was characterized by a decrease in delta, beta and gamma power compared to epochs of dreamless sleep.

We also confirmed these results at the individual level in 2 Zen subjects in whom overnight sleep recordings could be performed at the end of their retreats (see Methods). Suppl. Fig. 10–11 show topographies of PP states vs. dreamless and dreaming epochs obtained from serial awakening from either NREM sleep (stages 1–3) or REM sleep.

### PP variants with minimal perceptual content or bliss

We further assessed the neural signatures of a state described by both Vajrayana and Zen LTMs as PP-like but with minimal perceptual content (PP-mpc), typically faint feelings of a somatosensory or visual nature. The comparison of PP-mpc to MW (Fig. 7, upper panels, 15 LTMs) shows a trend towards decreases in delta, theta, beta and gamma power, similar to PP states, with most consistent gamma decreases found in posteromedial cortices. However, these changes were not as consistent as for PP (significant at uncorrected p<0.05, but did not survive correction for multiple comparisons). Average group topographies suggested gamma activity was preserved in both central and posterior occipital cortices. Direct statistical comparison revealed a trend towards decreased gamma activity in central, posteromedial and occipital cortices during PP compared to PP-mpc (at uncorrected p<0.05, but not surviving multiple comparison correction; Suppl. Fig. 12, upper row).

**Fig. 7.**
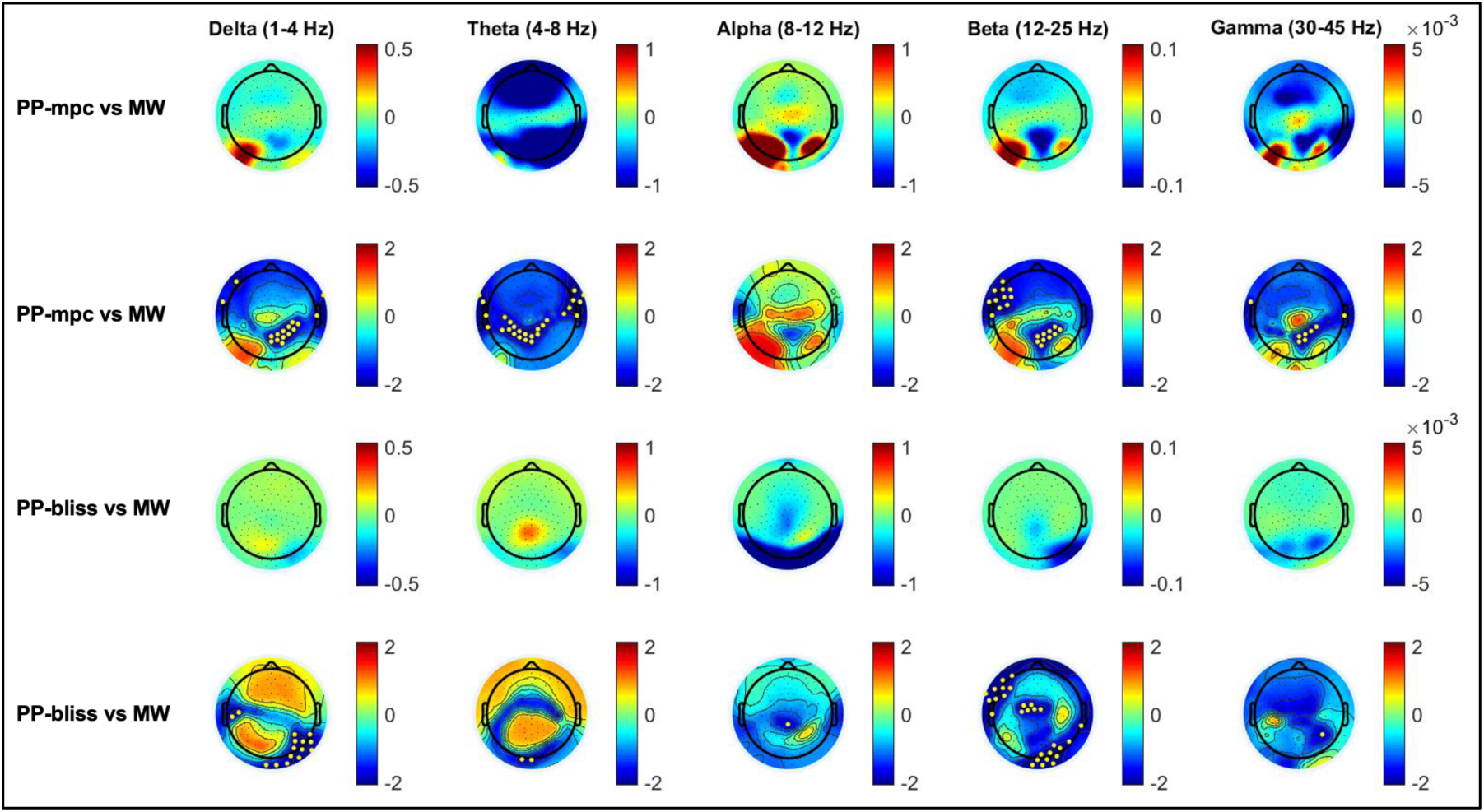
PP variants display similar signatures compared to PP states. First two rows: comparison of PP-mpc (PP with minimal perceptual contents) to MW in the same 15 LTMs who also reported PP states. Lower two rows: comparison of PP-bliss (PP with “a feeling of bliss arising from mind itself”) to MW in 9 Vajrayana LTMs who reported both states. Colors represent EEG power (first and third rows) and T values (second and fourth rows); yellow dots represent channels surviving uncorrected p<0.05.

We also investigated another PP variant traditionally described in Vajrayana as “bliss-awareness”—a PP-like state with “a feeling of bliss arising from mind itself”. The results of a group analysis in 9 Vajrayana LTMs with clean recordings during both PP-bliss and MW states (captured during 8^th^ Karmapa Guru Yoga practice) are shown in Fig. 7, lower panel. The neural signatures of PP-bliss are again similar to those of PP—with a trend for a broadband decrease in power, maximum for both delta and gamma band activity in posterior cortex, although less pronounced than during typical PP (uncorrected p<0.05, not surviving multiple comparison correction). Unlike PP-mpc, we did not find significant gamma-band activity differences between PP-bliss and PP within the same participants even at trend-level (nothing significant at uncorrected p<0.05 values; Suppl. Fig 12, lower row).

In 8 Zen practitioners, we also captured a state characterized by decreased thoughts but preserved perceptual contents during Shikantaza practice (Suppl. Fig. 13, upper panel). In line with OM findings described above, this state showed increased gamma power compared to MW. When transitioning to deeper absorption states, where thoughts stopped completely (with perceptual contents still preserved, then only minimally present (PP-mpc), then totally absent (PP)), gamma activity decreased progressively, at first in posteromedial areas, then in a more widespread pattern (Suppl. Fig. 14, three lower panels).

### PP: no-report paradigm

In a further set of experiments, we took advantage of a highly rehearsed, precisely timed practice that allowed us to assess the neural correlates of PP in a no-report paradigm and compare PP to other meditative states with richer thought-like and perceptual contents. The pre-timed 16^th^ Karmapa Guru Yoga practice in Vajrayana LTMs starts with a visualization-rich generation phase, then transitions to a mantra recitation phase, then to a dissolving (completion) PP phase. For the dissolving phase, LTMs are instructed to visualize that “all form disappears; there is now only awareness without center or limit” (see Supplementary Methods for a more detailed description). As the timing of the various phases was pre-determined, no report was needed. The results obtained in 11 LTMs by subtracting the average hdEEG topography from a pre-meditation baseline are shown in Fig. 8 (with bar graphs representing the whole-scalp average of gamma activity as the group mean +/- s.e.m. for each meditation phase).

**Fig. 8.**
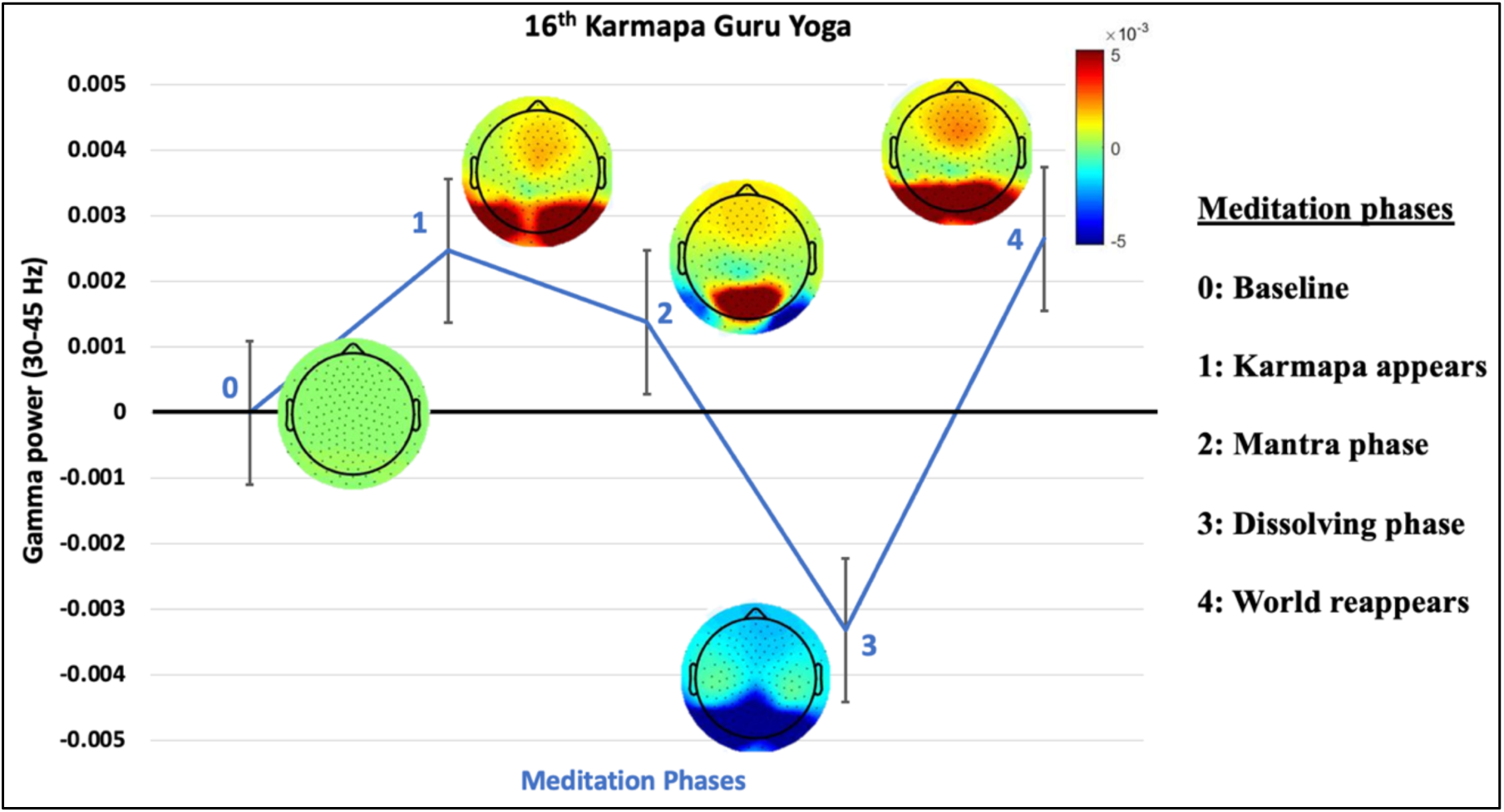
HdEEG results obtained during a highly rehearsed, pre-timed version of 16^th^ Karmapa Guru Yoga meditation (no-report paradigm). While other meditation phases were characterized by increased gamma activity compared to pre-meditation baseline (e.g., during phase 1, when Karmapa appears; during phase 2, with the mantra recitation; or during phase 4, when the world reappears), PP signatures with decreased gamma power were selectively found during the dissolving phase of the meditation (phase 3), when “all form disappears; there is now only awareness without center or limit”. Color-coded results show the group-average gamma-power topography for each phase (baseline-subtracted); bar graphs represent the whole-scalp average of gamma activity as group mean +/- sem. A detailed description of each meditation phase is presented in Supplementary Material.

As shown in the figure, the 16^th^ Karmapa dissolving phase (meditation phase 3) obtained through a no-report paradigm showed a similar neural signature as the 8^th^ Karmapa PP state obtained by delayed report. A whole-brain decrease in gamma power was indeed observed compared to baseline (p=0.026, non-parametric Wilcoxson signed-rank test); topographical analysis (Suppl. Fig. 15) confirmed that widespread gamma decreases survived SNPM-correction for multiple comparisons. At the single-subject level, the neural signatures of PP were even more consistent during the 16^th^ Karmapa than the 8^th^ Karmapa meditation in the same cohort (Suppl. Fig. 16), possibly owing to the no-report paradigm. The generation phase (meditation phase 1), which requires intense visualization, and the mantra recitation phase (phase 2) were instead dominated by a trend towards whole-brain increase in gamma activity (both p = 0.16, non-parametric Wilcoxson signed-rank test). Increased gamma activity was also seen at the end of the meditation session (phase 4, p = 0.041, non-parametric Wilcoxson signed-rank test), when LTMs are instructed to “let the world reappear”. Topographical results comparing these latter meditation phases to baseline however did not survive SNPM correction for multiple comparisons. Suppl. Fig. 16 also displays group averages for the simultaneous evolution of delta to gamma frequencies during the 16^th^ Karmapa practice.

## Discussion

We used hdEEG to investigate the neural correlates of PP, a state in which expert LTMs experience a vivid sense of spatial extendedness in the absence of any contents related to self, thought, or perceptual objects. We found that, compared to several other conditions both within and outside of meditation sessions, PP states are characterized by a distinct hdEEG signature: a pronounced and widespread decrease in gamma power, maximal in posteromedial cortex, consistent with reduced cortical activity, on a background of low delta power, consistent with high cortical arousal. In short, PP states seem to be associated with a cerebral cortex that is widely awake but minimally active. Similar findings were observed in PP variants accompanied either by minimal perceptual contents (PP-mpc) or with a feeling of bliss (PP-bliss). These results were confirmed during PP states captured using a no-report paradigm (pre-timed Guru Yoga meditation).

Low or decreased delta activity during PP, compared to both within-meditation MW and other control conditions, is consistent with a state of aroused wakefulness. The high cortical arousal of PP states was confirmed by the direct comparison between the hdEEG of PP states and that of epochs preceding awakenings from dreamless sleep in the same LTMs. It is also in line with the description of PP provided by all subjects, as well as by their traditions (19). Far from being a state of dozing off, PP is a state of intense, vivid, heightened consciousness. The signatures of PP states also differ from those of “mind-blanking” reported in meditation-naïve subjects, which seem to correspond to short bouts of drowsiness (32, 33). The phenomenology and neural signatures are thus remarkably different between “contentless” states such as PP that are meditation-induced (“cultivated”) and drowsy states that occur “in the wild” (34).

Source reconstruction revealed that the decrease in delta and gamma power was widespread but maximal in posteromedial areas. Previous work has suggested that special states of absorption or states of “thoughtless emptiness” may be accompanied by a broadband decrease in EEG power (35). Other work has shown a decreased high-frequency power in posteromedial areas as a signature of ego-dissolution achieved by LTMs (27). In our case, the decrease in gamma-band cortical activity during PP was significant not only over the precuneus and middle-posterior cingulate cortex, which are known to be involved in self-related experiential contents (26, 36, 37) and thinking (25), respectively, but also over the frontal poles, which are involved in metacognition (38), and over the temporo-parietal junctions, involved in agency (39). Gamma activity was also decreased over the right temporal lobe and sensory areas, as well as in other parts of the prefrontal cortex. These findings are in line with the phenomenology of meditative PP states—the absence of a sense of self, of thought, and of perceptual contents—and with the associated suspension of cognitive functions such as reflection, control, and working memory (17, 18).

The neural signatures of a variant PP state with minimal perceptual content (PP-mpc) were overall similar those of “contentless” PP states, with most consistent decreases in gamma power over posteromedial and right temporo-parietal cortex. However, the decrease in gamma power was less marked, and some gamma activity was preserved over medial occipital areas and somatosensory cortices. These results are compatible with reports from meditators suggesting that simple bodily sensations were last to disappear in the transition from MW to full PP experiences (see also results in Suppl. Fig 12 obtained in Zen). Another variant of PP, accompanied by “a feeling of bliss coming from mind itself” (PP-bliss), was also associated with a widespread decrease in gamma power, again less marked than in PP. This finding is aligned with the Vajrayana teachings emphasizing the inseparability of bliss and emptiness (Tib. *detong*), which is core to the Guru Yoga practice (40, 41).

The hdEEG correlates of PP were markedly different not only from those of MW episodes within the meditation session, but also from those of a variety of states outside the session. These included resting wakefulness with eyes open and eye closed, watching a movie, imagining the movie, and active thinking. Just as important, the hdEEG correlates of PP were also specific, being markedly different with respect to other meditative states achieved by LTMs. These included various meditative states during which perceptual content was not abolished, such as OM, Zen Shikantaza states with decreased thoughts but preserved percepts, and Vajrayana visualization and mantra recitation phases. For example, previous studies have reported that a variety of other meditation states, such as OM or non-dual compassion states, can be accompanied by increased gamma activity (42, 43). Overall, our findings emphasize the importance of characterizing the phenomenology of various meditative states to identify their specific and reproducible neural signatures (34). The distinctive EEG signature of PP is also in line with the ability of practitioners to distinguish such states from other meditative states as well as from dreamless sleep.

This study was undertaken to assess a counterintuitive prediction of integrated information theory (IIT): that the neural substrate of consciousness can support a vivid experience even when it is minimally active, as long as its causal powers are preserved (44). The phenomenology of PP states is described by meditators as a vivid feeling of “presence,” hence the name, further characterized as a sense of vast extendedness, devoid of self, thoughts, and perceptual objects. According to IIT, such a vividly conscious experience should be supported by a hierarchical, lattice-like substrate that is virtually silent, but with preserved causal powers. Therefore, IIT predicts that, to the extent that EEG gamma power can be taken as a reflection of neural activity, it should be low over the likely substrate for conscious contents in posterior-central areas of the cortex (45). Importantly, EEG delta power over the same regions should also be low, at least to the extent that its increase reflects a breakdown of causal interactions (46). Specifically, it should be as low as in other vividly conscious states, rather than high, as observed during dreamless sleep and anesthesia (22).

In line with these predictions, the results presented here show that vivid states of PP are associated with a hdEEG background of low gamma power and low delta power. Widespread low gamma power, which was lower than in all other awake conditions tested, is suggestive of low neural activity throughout the cortex. Low delta power—much lower than in dreamless sleep and as low or lower than in MW, movie imagining, and movie watching—indicates that causal links within the cortex are in working order, rather than interrupted by the bistability that characterizes slow wave sleep (46),(47). Overall, this pattern contrasts with what would be expected in a drowsy state (high delta) or a state of widespread activation (high gamma), thereby aligning with IIT’s predictions.

Of course, EEG cannot provide a direct readout of neuronal activity, so it cannot provide definitive evidence for IIT’s counterintuitive prediction. However, had the findings been different, say an increase of gamma activity in PP vs. MW, they would have provided evidence *against* IIT’s prediction. Thus, despite their limitations, the present experiments constitute a valuable test of IIT’s predictions and provide evidence supporting it. By the same token, the present results pose a challenge to current computational/functionalist or dynamical system accounts, according to which being conscious requires some kind of “computation” or “information processing” (7). Indeed, our findings suggest a likely disconnect between the vividness of conscious experiences and the level of neuronal activity.

Besides stipulating the necessary and sufficient conditions for being conscious, IIT also aims at accounting for the quality of an experience, which it identifies with the cause–effect structure specified by its neural substrate in its current state (8, 11). In particular, previous work has shown that lattice-like substrates, such as those found in posterior cortex, specify cause–effect structures that capture the feeling of “extendedness”. A phenomenal analysis of the properties of spatial extendedness indicates that the visual field is composed of phenomenal distinctions (spots) bound by the phenomenal relations of inclusion (every spot is included by or includes some other spot), connection (every spots partially overlaps with some other spot), and fusion (the union of connected spots is another spot, (10)). As shown by computer models based on IIT, a lattice-like substrate specifies a kind of cause-effect structure, called an “extension”, that is composed of causal distinctions bound by causal relations of reflexivity, inclusion, connection, and fusion. Such a structure can account for the corresponding phenomenal properties of spatial extendedness, as well as for derived spatial properties such as regions, locations, and distances (11). Notably, a grid-like lattice in a state where all units are inactive (but not ‘inactivated’ by bistability) specifies such a structure just as much as in other states, where some units are active. The only difference is that the extension is completely homogenous in the case of inactivity, while it is warped by inhomogeneities when some units are active and others inactive (11). Remarkably, LTMs of various traditions, including our subjects, describe PP states as a vast, homogeneous expanse, sometimes qualifying the experience as “mirror-like”, “panoramic”, or “luminous” (please see examples in the two phenomenological interviews attached in Supplementary Material). It remains to be determined whether some phenomenological descriptions of PP states may differ systematically across different contemplative traditions.

### Limitations of the study

While the present work demonstrates that a vivid experience of PP is accompanied by a widespread reduction of EEG power, especially in the gamma range, it cannot demonstrate directly that neuronal activity in the underlying cortex is in fact reduced or minimal. In both monkeys and humans, increases and decreases in cortical gamma power are tightly correlated with local increases and decreases in neuronal population firing (14–16). However, it is certainly possible that, at the level of individual neurons or small local cortical populations, activity may not decrease during PP—in fact, it might even increase— without being detected by the hdEEG. Moreover, neuronal activity in some cortical regions located far from the scalp, are often invisible to the EEG. Intracranial recordings in humans during PP-like states may shed light on these possibilities.

PP states are challenging to capture with neuroimaging. Indeed, vivid meditative PP states like those obtained in the present study are typically a few second long, and they could be achieved only in 22 out of 38 meditators, even though each meditator was recorded over 5-7 days of retreat. Data reduction to 15 LTMs for our main analysis was further necessary due to strict data quality requirements for including uncontaminated EEG gamma-band activity.

Another limitation of this study is that PP was often collected *after* MW. However, the lack of any difference in delta power between the two conditions suggest that differences in other frequency bands are unrelated to circadian or homeostatic changes in drowsiness. There are also potential order effects that could be remediated in larger studies. Our sample size of 22 LTMs is higher than most recent reports presenting brain activity during deep meditation states (e.g. (48–50)). Even so, research in larger cohorts could also shed light on potential effects of meditation expertise and differences in cultural background.

Several exploratory findings were reported here even when they did not survive multiple comparisons, in line with recent recommendations to include not only statistical peaks but also near-threshold neuroimaging results in order to enhance reproducibility and cross-comparison of studies (51). While informative, such exploratory findings should be validated by further studies. Finally, we only obtained two formal phenomenological interviews about PP states—one in a Zen participant and one in a Vajrayana participant. Further phenomenological validation of PP states should be performed in larger cohorts.

Lempel-Ziv complexity, computed on continuous EEG signals, did not significantly differ between PP and MW states. It should also be mentioned that, according to IIT, it is the causal complexity, rather than observational complexity measures, that matters for consciousness (52). In general, the causal complexity that characterizes states of PP is expected to be comparable to other vividly conscious states, despite the absence of usual contents such as objects, thoughts and self in PP states (52). By contrast, IIT predicts that complexity should decrease significantly from conscious to unconscious states, as confirmed by studies employing perturbational approaches (*52*).

In future work, it will be instructive to compare the neural signatures of meditative PP with those achieved by LTMs during “clear light” practices in lucid dreams (53) and with certain states induced by psychedelics occasionally described as being phenomenologically similar (54). The correlation between the occurrence and depth of PP states and measures of mindfulness, decentering (55), fear of death (56), hedonic and eudaimonic well-being should also be explored. It will also be important to investigate the neural mechanisms underlying LTMs’ ability to reach states of PP, including the role of attentional training. Finally, it will be important to replicate and extend the present findings by targeting meditative states similar to PP using intracranial EEG, ideally combined with single-unit recordings, in subjects with epilepsy (57). Towards this goal, ongoing work aims to induce PP-like states in meditation-naïve subjects using paradigms inspired by millennia-old traditions, such as a fading meditation bell (58).

## Materials and Methods

### Participant selection

We recruited Vajrayana and Zen participants for the present study. Inclusion criteria consisted in at least 3000 hours of lifetime meditation experience, current meditation practice of several times a week, and previous meditation experience in a retreat setting. Study procedures were approved both by UW–Madison Health Sciences’ and University of Granada’s Institutional Review Boards.

Vajrayana practitioners were recruited from the Tibetan Karma Kagyu lineage (Diamond Way Buddhism organization) because their Guru Yoga practice explicitly targets the achievement of PP states during the “dissolving phase” of the meditation. Recruited participants had all completed a series of intensive practices called the *Ngöndro* (Tib.) at least once and performed both 8^th^ and 16^th^ Karmapa Guru Yoga as their main practices (see Supplementary Material: Vajrayana practice details).

Because PP states are not an explicit target of Zen practice, Zen participants were recruited if they indicated they had had at least one PP experience and thought it was possible to replicate this in an experimental setting during a seven-day meditation retreat. They also all had experience with Shikantaza or equivalent meditation practice.

### Retreat and hdEEG acquisition schedule

All retreat experiments took place in naturalistic settings, in meditation centers belonging to the LTMs’ respective spiritual schools. Up to three LTMs were recorded during the same week. A member of the research team stayed on-site with the LTMs and ensured that their subsistence needs were met so that they could fully focus on the practice. Retreat schedules and rating scales were adapted for Zen and Vajrayana traditions (see Supplementary Methods). After informed consent was obtained, questionnaires were administered to quantify the participant’s total hours of experience for each type of practice, including in retreats. LTMs were then fitted with a 256-channel hdEEG net and given control tasks, including resting with eyes open and eyes closed (2 min.), a series of video clips (1 min) that were then followed by an imagination task where they closed their eyes and imagined the same video they had just watched (1 min); this sequence was repeated three times and the videos were randomized from a pool of six videos each session. Additionally, subjects performed an active thinking task where they were instructed to count backwards starting at 300 by sevens (3 min), and a short open-monitoring meditation session where they wore an eyeshade and earplugs to reduce ambient stimulation and were told to just allow thoughts to come and go without paying any attention to them but also not trying to reduce them (3 min). After the control tasks, subjects were then trained to meditate while rating their experiences using a mouse click, after emerging from various absorption states. At the end of each meditation session, participants reported on their sleepiness (0–4 scale) and were invited to provide open-ended reports about their meditative states. To ensure accurate understanding and communication of the target phenomenal states to be signaled within each meditation practice, LTMs of the two traditions who were members of the research team assisted with pre-briefing, de-briefing, and assistance throughout the retreats.

For Vajrayana participants, the hdEEG net was applied on the day of arrival to familiarize participants with the controls and the rating procedure, then for 4 hours per day every day of the 5-day retreat, while the rest of the time was spent meditating without the hdEEG net. After net application, control conditions were performed, then a pre-timed 16^th^ Karmapa Guru Yoga session (no-report paradigm). Then, participants were instructed to perform 8^th^ Karmapa Guru Yoga, during which they signaled target phenomenal states with mouse clicks. All participants were instructed to keep their eyes open throughout the meditation recordings (including mind-wandering epochs), even if their usual practice was to at times keep their eyes closed at home. Participants were also instructed to keep eyes open in all control conditions, except of course the ‘resting eyes-closed’ one.

For Zen participants, the hdEEG net was applied on the first day and during the last three days of the week-long retreats. Participants used mouse clicks to signal target phenomenal states during meditation sessions. Control conditions were repeated at the beginning and the end of each recording day (for analysis, the cleanest session of the two was kept for analysis in each LTM; yielding 6 morning and 2 evening sessions). The meditation rounds were all 50 min long followed by 10 min of walking meditation. All meditation rounds were timed by the researcher(s) running the study. These 50 min rounds were repeated throughout the day for an average of 8–9 rounds totaling 6∼7hrs of formal meditation each day.

Two LTMs (2952.83 ± 468.34 lifetime hours of practice; age, 41.87 + 6.95; age range, 38–52; 1M 1F) in the Zen tradition that had reported a PP state by the end of day 6 were recorded at WI Sleep for an overnight Serial Awakening (SA) recording. Once they arrived at the sleep lab, we applied a 256 HdEEG net (Electrical Geodesics, Inc., Eugene, Ore.) for sleep recordings and additional submental electromyogram and electrooculogram to perform sleep staging using Alice^®^ Sleepware (Philips Respironics, Murrysville, PA) Throughout the night, they were awakened repeatedly (59) and asked to report whether, just before the awakening, they had been experiencing anything (DE), had been experiencing something but could not remember the content [DE without recall of content (DEWR)] or had not been experiencing anything [no experience (NE); (59)]. The serial awakening protocol has previously been reported in detail (59). The group from the Vipassana tradition underwent the same protocol. To summarize, participants were awakened periodically throughout the night using a computerized tone delivered through headphones lasting 1.5s. They were then asked to report, “what was the last thing going through your mind before the alarm?”. If they reported something (DE), they then underwent a structured interview through an intercom where they were asked to describe details of the report followed by a series of questions probing the duration, complexity, whether the experience was self-centered vs environment centered, how much they were perceiving, thinking, or imagining, the degree of positive or negative affect, and a few other questions regarding control of dream content and specific contents of the dream. Awakenings were performed at intervals of at least 20 min, in N2 or REM sleep. Participants were required to have been asleep for a minimum of 10 min and must have been in a stable sleep stage for a minimum of 5 min for an awakening to occur. At the conclusion of the awakenings, subjects were allowed to sleep adlib.

### High-density EEG preprocessing

Data from days 5–7 of retreats were kept in the analysis. We limited the analysis to these days because of the expected occurrence of longer and more stable PP states towards the end of the retreat compared to during the first days (where PP occurrence was expected to be both short and rare). We applied a 0.1–Hz first-order high-pass filter and focused our analysis on 185 channels with highest signal-to-noise ratio, excluding the neck and face. We then applied a linear finite impulse response bandpass filter between 1–50 Hz (using EEGLab pop_eegfiltnew FIR filter algorithm), removed the noisiest sections and bad channels visually, and then ran adaptive mixture independent component analysis (AMICA) (28). We inspected the AMICA decomposition and removed segments with artifacts in which ICA decomposition failed to separate brain activity from physiological noise time courses (Supplementary Fig. 1, upper panel) and then re-ran the AMICA algorithm without remove components. Once the AMICA decomposition showed a clear separation between the first twenty brain and artifactual components (comprising most of the signal variance; see Supplementary Fig. 1, lower panel), ICA components contaminated by artifacts were removed. We then interpolated bad channels using spline interpolation in EEGLAB and average-referenced the signal. We then extracted epochs preceding by 8 to 3-seconds prior to the reports of events of interest (e.g. reports of PP, PP-mpc, PP-bliss and MW during meditative states). For the 16th Karmapa Guru Yoga sessions, we extracted epochs 5–10 seconds after the end of each instruction (these time points corresponded to our estimation of reliable epochs capturing the phenomenology of target states in the report vs no-report cases, based on our previous studies tracking quantifying changes in power in 256-electrode high-density EEG recordings during both wakefulness and sleep dreams (22, 25)). For the OM and all other controls tasks, we extracted epochs of 5 seconds of continuous clean data starting at least 5 seconds after beginning the task and no more than 5 seconds before the end of tasks. For both the 2 Zen meditators and the 13 Vipassana LTMs that had serial awakenings (Fig. 5), we extracted 5 second epochs immediately before each serial awakening from N2–N3 sleept hat was reported as dreamless. Data were similarly extracted in 1 Zen participant who also had serial awakenings with reports of dreams from both NREM sleep and REM sleep (Suppl. Fig. 7). We extracted epochs -5–0 seconds before each serial awakening. In one Zen participant, we also extracted 5 seconds of continuous clean data from stages N1, N2, N3, and REM sleep and displayed them on the same power scale for comparison to PP and MW. Topographies of delta (1–4 Hz), theta (4–8 Hz), alpha (8–12 Hz), beta (15– 25 Hz) and gamma (30–50 Hz for Zen data, 30–45 Hz for Vajrayana data) were computed in each condition for statistical analysis. Of note, the gamma frequency range was kept below 45/50 Hz to stay away from line noise artifacts (60 Hz for USA data, 50 Hz for Spain data) while applying FIR filters. We did not analyze higher EEG frequency data because of expected lower signal-to-noise ratio for high-gamma. Given the importance of EEG gamma activity to be of neural origin to properly test our hypothesis, we only included data from participants where no residual muscular or eye movement artifacts could be seen in post-AMICA EEG topographies (manifesting as focal gamma activity spanning <3-5 channels, see many examples of 256-electrode muscular components in Suppl. Fig. 2). Suppl. Table 1 indicates which participants were kept in each condition across report and no-report PP conditions, additional controls, and sleep.

In order to calculate Lempel-Ziv complexities between the MW and PP conditions for all 15 subjects, the first 5 seconds of MW and the last 5 seconds of PP were extracted for each subject. We then took the average of all 185 channels, calculated the mean activity value, then binarized the data into a string of 0 and 1s (if below or above the mean), which we then used to calculate the Lempel-Ziv complexity of each condition using the LZ76 algorithm (60) as implemented in Matlab (https://www.mathworks.com/matlabcentral/fileexchange/38211-calc_lz_complexity). We then averaged all individual complexities scores per condition and compared them across groups.

Source-reconstruction was performed for MW and PP conditions in the 15 LTMs who reported both states. Forward model used a realistic head template with boundary element methods (BEM) and sLORETA priors, as implemented in Brainstorm (www.neuroimage.usc.edu/brainstorm). We created separate source-space topographies for EEG gamma to delta frequency bands for PP and MW.

### Statistical analysis

Except when specified below, hdEEG topographical statistics were performed using a within-subject design, using paired t-tests testing for within-subject differences across conditions in each frequency band of interest. As in previous work, scalp-level statistics were corrected for multiple comparisons using statistical non-parametric mapping (SNPM), and source-space statistics using false-discovery rate (22).

The comparison between PP and controls (Fig. 5) used similar participants in 13/15 subjects (two Vajrayana participants who achieved PP during 8^th^ Karmapa practice had noisy control data, and they were replaced by two other participants who also had clean 16^th^ Karmapa data). Unpaired t-tests were used to compare 16^th^ Karmapa phases to baseline, as 10/11 subjects had clean data in all phases, but some data were noisier in one subject both for phases number 2 and 4 (‘Karmapa appears’ and ‘the world reappears’). Because the comparison between PP and dreamless sleep involved different subjects, unpaired t-tests were performed testing for between-group differences in hdEEG power (with one group of LTMs in PP states, and one group of LTMs in unconscious sleep states). Again, all scalp-level statistics were corrected for multiple comparisons using SNPM.

## Acknowledgments

This work was made possible thanks to the generous support of the Templeton World Charity Foundation and the Tiny Blue Dot Foundation to MB and GT. We thank Jeremiah Hendren for helpful comments on the manuscript. Study data will be shared with other investigators upon reasonable request. GT is Board Member and has a financial interest in Intrinsic Powers Inc.

## Supplementary material

### Supplementary discussion

#### Additional methodological considerations

The results presented here focus exclusively on gamma frequencies below 50 Hz. We had previously validated this frequency band to localize conscious contents with hdEEG (1, 2). Moreover, based on our extensive experience with 256-channel hdEEG, we are confident in our ability to isolate neural sources from muscular or ocular artifacts frequencies below 50 Hz through a rigorous pipeline, but much less confident above 50 Hz. Previous studies have shown that most gamma activity picked up by raw EEG signals in the awake state is likely of muscular origin (3). Thus, we made every effort to optimize the separation of muscular from cortical sources of gamma power through our custom preprocessing pipeline combined with AMICA (Suppl. Figs. 1-2). This allowed us to unequivocally identify a large number of components that were related not only to eye movements, heartbeat, muscles, head movements, and single-channel popping artifacts, but also a small number of components with no obvious contamination by muscle in either topography or time course. To ensure that differences in gamma activity that we identified were due to differences in cortical signals rather than to artifacts, especially muscle activity, we also excluded components for time courses that appeared mixed. Crucially, the ICA decomposition was performed on the entire dataset, and the same components were removed in both the PP and MW conditions. (Some LTMs of the Vajrayana group who achieved PP during 8^th^ Karmapa practice could not be included in the analyses because their data displayed persistent muscular/ocular artifacts in post-AMICA topographies).

Transfer entropy could not be reliably computed on the present dataset, because it would require longer epochs than those we can reliably assume to correspond to specific experiential states (based on our previous studies, we chose 5-second long epochs (1, 2)). Similar considerations apply to continuous EEG spectrogram analysis, which has been employed to target more enduring differences in the level of consciousness, such as continuous propofol infusion (4). Such methodologies may be more appropriate for more prolonged PP-like states, like those induced by 5-MeO-DMT (5).

Participants were instructed to keep eyes consistently open during meditation recordings. Their compliance with this instruction during PP states is further confirmed by the presence of similar occipital alpha power during PP compared to MW (Fig. 2, with if anything, a decrease in alpha power during PP at uncorrected p<0.05), a significant decrease alpha power in occipital channels during PP compared to eyes-closed resting condition (surviving SNPM correction), and no difference in alpha power with resting eyes-open condition at uncorrected p<0.05.

The attainment of PP is reported retrospectively by trained LTM upon emerging from the state. It remains to be elucidated through which neural mechanisms the occurrence of PP experiences can be encoded in short-term memory and then recalled. Future studies may also aim to relate the phenomenology of PP states to changes in physiological variables such as breathing rate.

#### Comparison of meditative PP to psychedelic-induced altered states

Similarities and differences in phenomenology should generally translate into similarities and differences in neural correlates. For example, changes in contents of experience during psilocybin infusion typically combine ego-dissolution with decreased cognitive control (6) and decreased integration of bodily signals (7). On the other hand, in contrast to PP, top-down visual imagery may increase (8). Correspondingly, significant decreases in gamma power were reported after psilocybin infusion, which were most prominent in sensorimotor and prefrontal regions, as well as in some smaller areas within the superior parietal cortex and precuneus (9). However unlike in our PP study, gamma activity did not decrease in visual regions, consistent with phenomenology. Future studies should directly compare brain activity during meditative PP states with those of PP-like states induced by other psychedelics/transcendelics such as 5-meo-DMT (5).

#### Zen practice details

All Zen participants practiced Shikantaza meditation during the retreat. Although the participants had various Zen background and teachers, all but one had over 10 years of continuous meditation practice and a history of multiple in-person retreats. They also all indicated they had at least one prior experience of pure presence and thought there was a good chance they could replicate it during our study. Shikantaza is a Japanese term often translated as “just sitting” described here:

> Shikantaza, or “just sitting,” is meditation without a goal. It is boundless—a process that is continually unfolding… One way to categorize the meditation practice of Shikantaza, or “just sitting,” is as an objectless meditation. This is a definition in terms of what it is not. One just sits, not concentrating on any particular object of awareness, unlike most traditional meditation practices, Buddhist and non-Buddhist, that involve intent focus on a particular object…But objectless meditation focuses on clear, nonjudgmental, panoramic attention to all of the myriad arising phenomena in the present experience (Loori, 2012)

The word Shikantaza is similar in meaning as the Chinese word for “just sitting” translated in English as “silent illumination”. The equivalent in the Chinese school of meditation practice known as Chan (Zen originally came from the word Chan both meaning meditation) of “silent illumination” is best described here by Chan Master Sheng-yen:

> The style of meditation called “silent illumination” is one of the great practices of the Chan (Zen) tradition. Silent illumination originated around the eleventh century, and its greatest advocate was Master Hung-chih Cheng-chueh (Hongzhi Zhengjue) of the Ts’ao-tung (Caodong) sect, which became the Soto sect in Japan. In Tibet, the mahamudra practice is very similar. The practice originated in India, where it was called shamatha-vipashyana, or serenity-insight. The aim of this practice is a mind unburdened with thoughts. This leads the mind to profound awareness about its own state (Loori, 2012).

But this is not a “dead” sitting where one is unalert. This alert awareness is described in James Austin’s book, Zen and the Brain as an alert condition, performed erect, with no trace of sluggishness or drowsiness. During retreats in particular, Shikantaza can be shifted, or shifts itself, into long moments of extra-attentiveness. This means a kind of listening as though one were blind, of looking as though one were deaf, of feeling as though all one’s pores were open and receptive. The senses seem to stretch out to close the gap between stimulus and perception, that interval which had once been occupied by the old judgmental barriers of interpretation. At such times, the meditator enters a state of high perceptual expectancy. It is the way one listens, knowing that a tiger lurks in the jungle nearby (Austin, 1998). This extra-attentive state, as Austin describes it, leads one into a type of samadhi (VI-B Absorption with Senate Loss: Internal Absorption. Table 10 pg 302 (Austin, 1998) where eventually the meditative absorption leads one’s experience of body and mind to drop away leaving just the experience of pure presence with no objects remaining, including the self.

Even during long retreats this can be a fairly rare experience and is not sought after but arises through deep meditative absorption (samadhi) as the practice progresses. Half of those recruited (8) reported this state at least once during the retreat by pressing a silent mouse button when they came out of the state and found themselves returning to perceptual/thought like awareness. In all cases, this only occurred during the last 2-3 days of the retreat. They were initially trained over the first 3-4 days of the retreat to report when they came out of various states of emptiness as indicated on the Zen Pure presence rating scale in the supplemental. There was also an opportunity to speak with an experienced, long-term meditator who is familiar with such states in order to clarify any confusion that may have arisen during the retreat.

## Vajrayana practice details

All Vajrayana Buddhist participants were practitioners of the Tibetan Buddhist Karma Kagyu lineage (Diamond Way Buddhism organization) and followed the same system of meditation practices. For all of them, the Guru Yoga meditation on the 16^th^ Karmapa had been the general meditation they had been practicing regularly for years—many of them daily. They had also all completed a series of practices called the *Chag Chen Ngöndro* (Tib.; Eng. “Preparation for Mahamudra Practice,”)—known in English as the “Four Foundational Practices” (Nydahl, 2019), the “Four Uncommon Preliminaries” (Karmapa Wangchug Dorje, 2009), or the “Four Special Foundations” (Kongtrul, 1977)—and were all currently practicing the advanced Guru Yoga meditation of the 8^th^ Karmapa Mikyö Dorje (Tib. *Tun Shi Lame Naljor*; Eng. “Meditation on the Lama in Four Sessions”). Following the established path of the Karma Kagyu lineage towards the ultimate Mahamudra practice, they had all adopted this Guru Yoga of the 8^th^ Karmapa as their main, regular meditation practice.

Generally speaking, Guru Yoga meditations focus on awakening the qualities of an enlightened mind, as they are represented or expressed by one’s teacher (Magnussen, 2003; Malinowski, 2013). Within Tibetan Buddhist traditions, the Guru Yoga is hailed as the most effective meditation method to achieve full realization of these qualities (Sobisch, 2006). For example, the contemporary Buddhist master Dzongsar Jamyang Khyentse Rinpoche summarizes the practice in his preface to the book *Guru Yoga* (Dilgo Khyentse, 1999, p. 18): “Guru Yoga is the quickest, most effective method for attaining enlightenment and is the one path in which all other paths are complete. Guru Yoga includes renunciation, bodhicitta, development (*Kyérim*) and completion (*Dzogrim*) meditation, and mind training (*Lojong*), which is why we can say Guru Yoga is the embodiment, or the essence, of all paths.”

Given the centrality of the Guru Yoga for Vajrayana practitioners, our research focused on their two main Guru Yoga meditations: the Guru Yoga of the 8^th^ Karmapa (Mikyö Dorje; see Rheingans, 2012, 2017) and of the 16^th^ Karmapa (Rangjung Rigpe Dorje; see Bausch, 2018, 2021). With growing meditation expertise, all practitioners increasingly emphasize the dissolving or completion phase of these meditations (Tib. *dzog rim*). This phase was the perfect target for our study because practitioners aim to rest mind in a state of pure presence.

The 8^th^ Karmapa Guru Yoga was the main practice all participants engaged with during their week-long retreats. They practiced it in a flexible, self-directed manner with no restrictions in terms of timing or progression through the various stages of the practice. They were instructed to report specific states of awareness during the dissolving (or completion) phase of this practice (see Methods).

In addition, we used the 16^th^ Karmapa Guru Yoga meditation to quantify the neural signatures of pure presence using a no-report paradigm. Rather than engaging in self-paced meditation practices, here the participants followed a scripted, pre-recorded and precisely timed guided meditation. Compared to the 8^th^ Karmapa Guru Yoga, this practice has a relatively condensed and simple structure. It was composed by Karmapa Rangjung Rigpe Dorje to be easily accessible and easy to practice. It emphasizes the key points of Guru Yoga in a compact fashion, making the fixed timing we employed feasible. The fixed timing allowed us to know when exactly the practitioner enters the dissolving phase, without relying on self-report. To reduce the impact of the auditory instructions and to ensure that the different phases of the meditation are of equal length, we recorded an abridged version of the guided meditation instructions, which had been approved by Lama Ole Nydahl (for the full text of the meditation, see p. 147ff in Nydahl, 2009). Here our main question was whether neural signatures of PP can also be detected when externally instigated, and whether the neural signature is similar to self-reported states of PP as detected during the 8^th^ Karmapa Guru Yoga.

|  | 16<sup>th</sup> Karmapa Guru Yoga key meditation phases | Brief description and representative quotes (original quotes are indicated) |
| --- | --- | --- |
| 0 | Pre-meditation baseline | Resting with eyes open until meditation starts |
| 1 | The form of the 16<sup>th</sup> Karmapa appears | The meditation starts by contemplating the four basic thoughts, taking refuge and awakening *bodhicitta* (Skt.), the altruistic motivation for one's practice. Then, the meditator calls the form of Karmapa to mind, who appears in front of them in space, and focuses on awakening the enlightened qualities of body, speech, and mind: "In front of us appears the golden, transparent form of the 16<sup>th</sup> Karmapa, a radiant field of energy and light." |
| 2 | Mantra phase | Participants are instructed to repeat the mantra only internally / in their mind (i.e., without verbalization): "We now use the mantra *karmapa chenno*. It means, 'Power of all buddhas, work through us.'" |
| 3 | Dissolving phase | "All form disappears; there is now only awareness without center or limit." |
| 4 | World reappears | "Our surroundings, this world and all worlds appear, perfect and pure." |

## Supplementary Table

**Suppl. Table 1.**
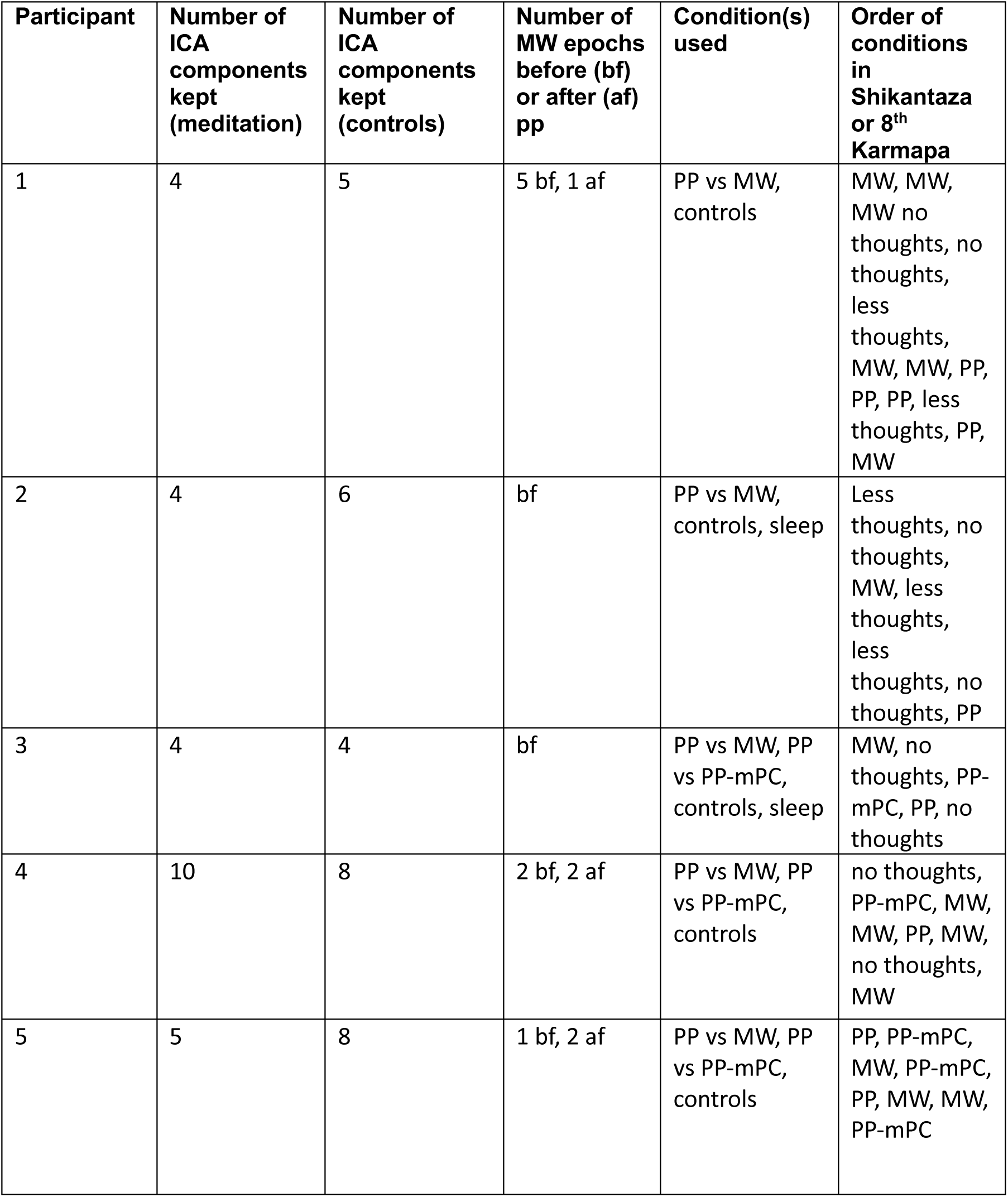
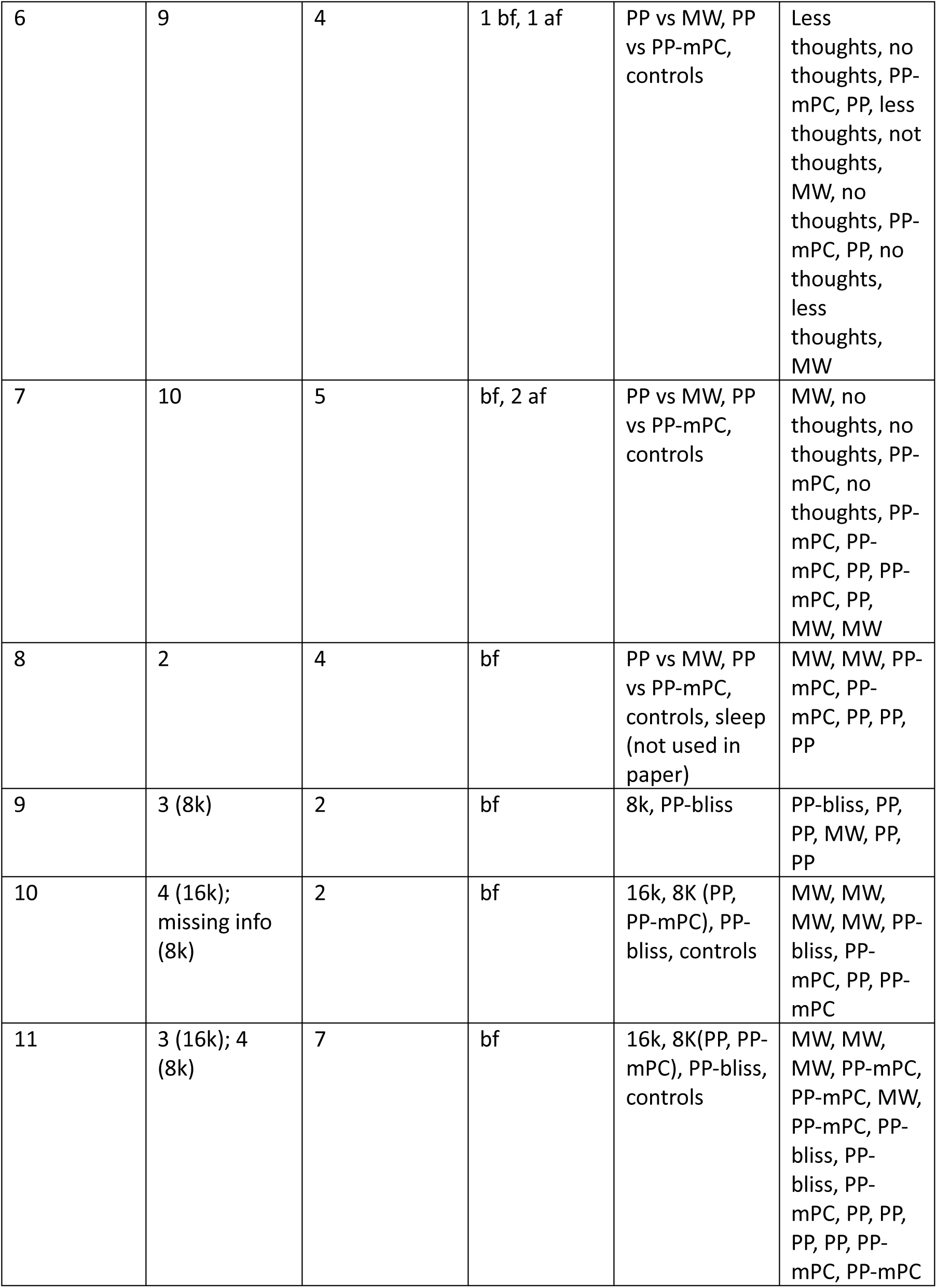
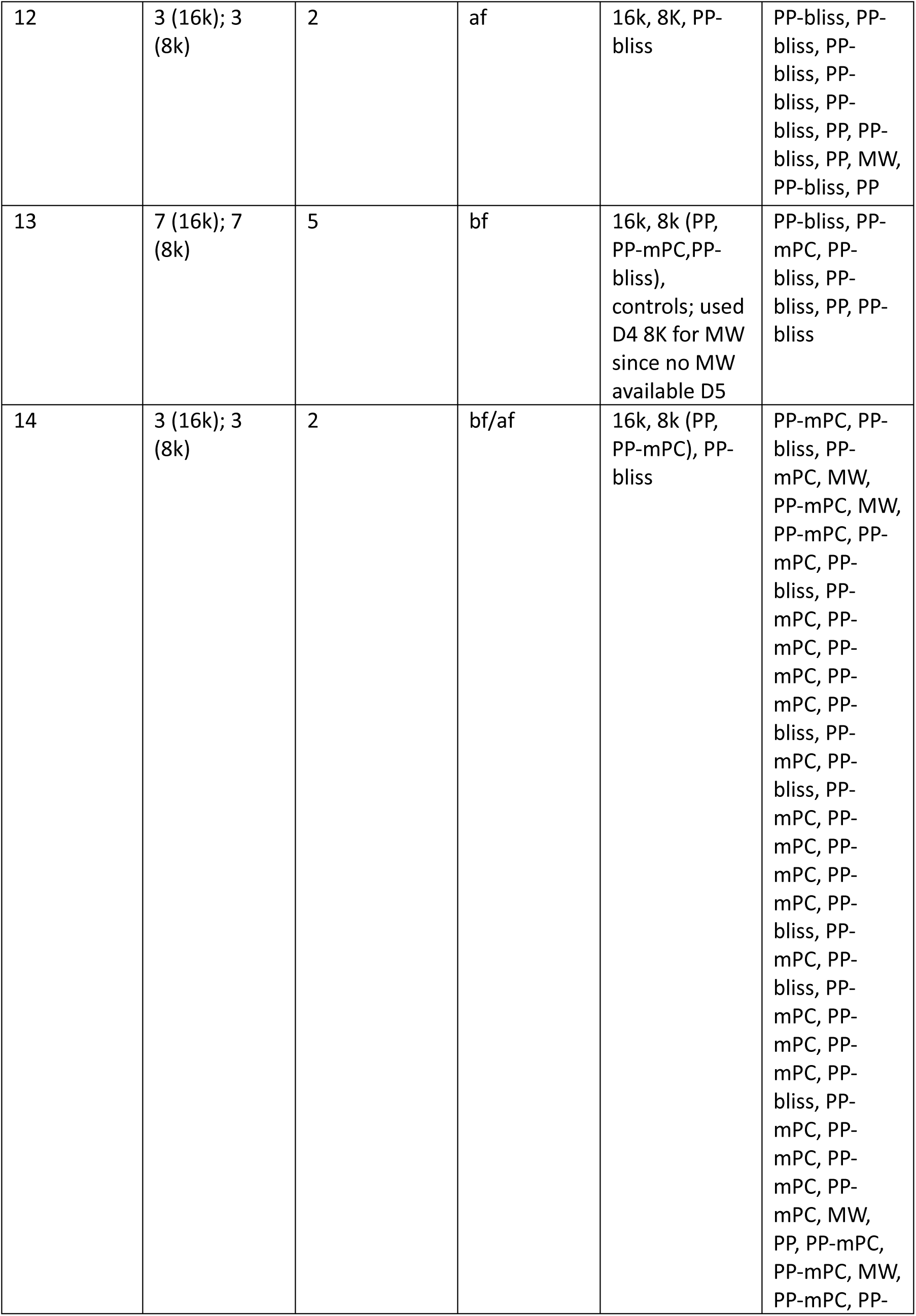
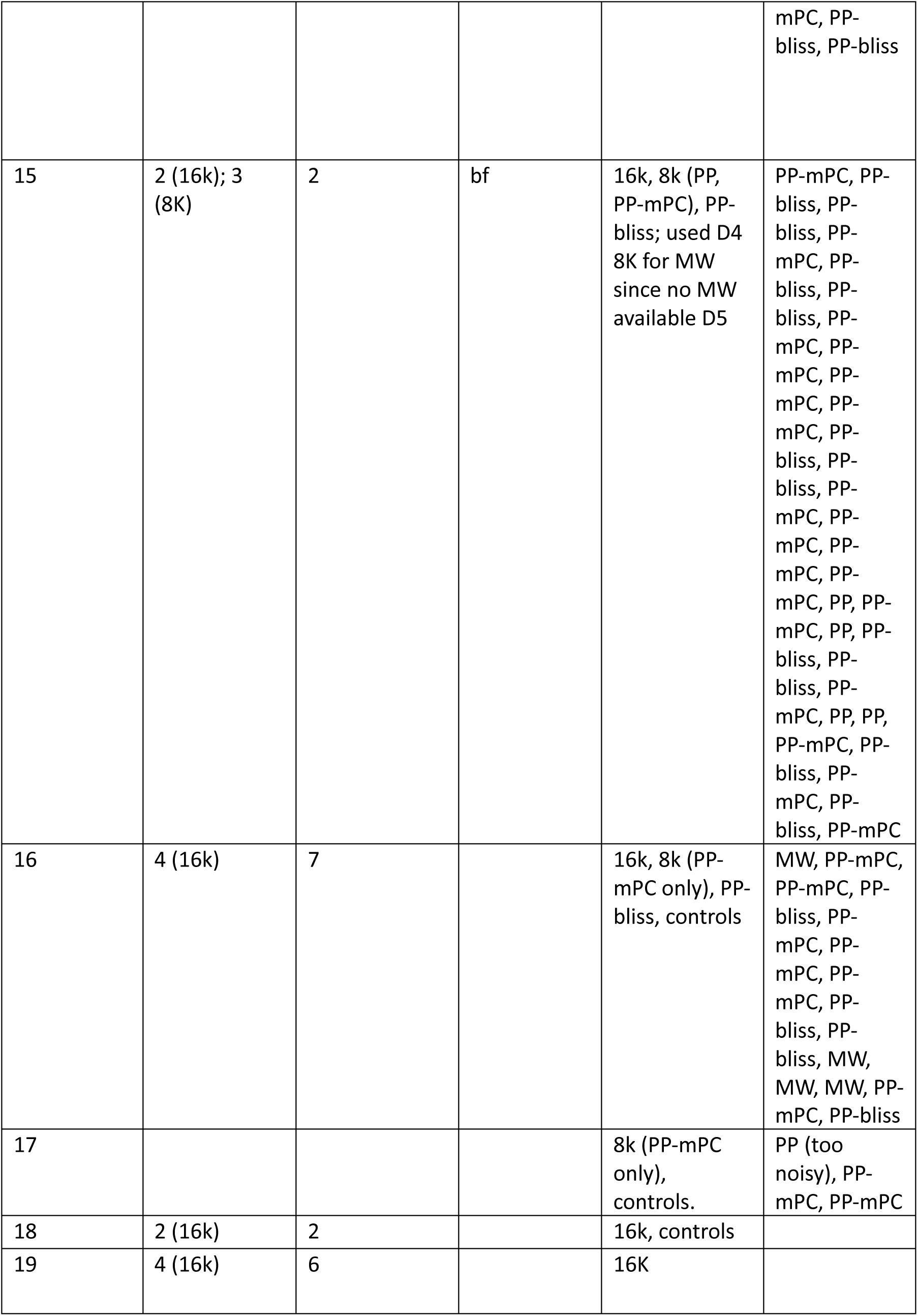
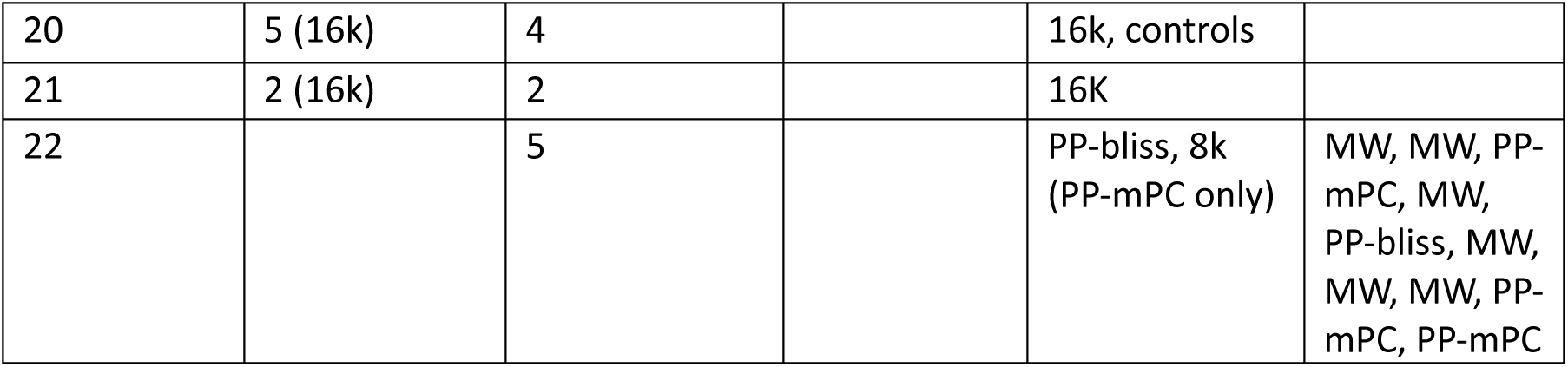
List of participants included in each condition, with condition order.

## Supplementary Figures

**Suppl. Fig 1.**
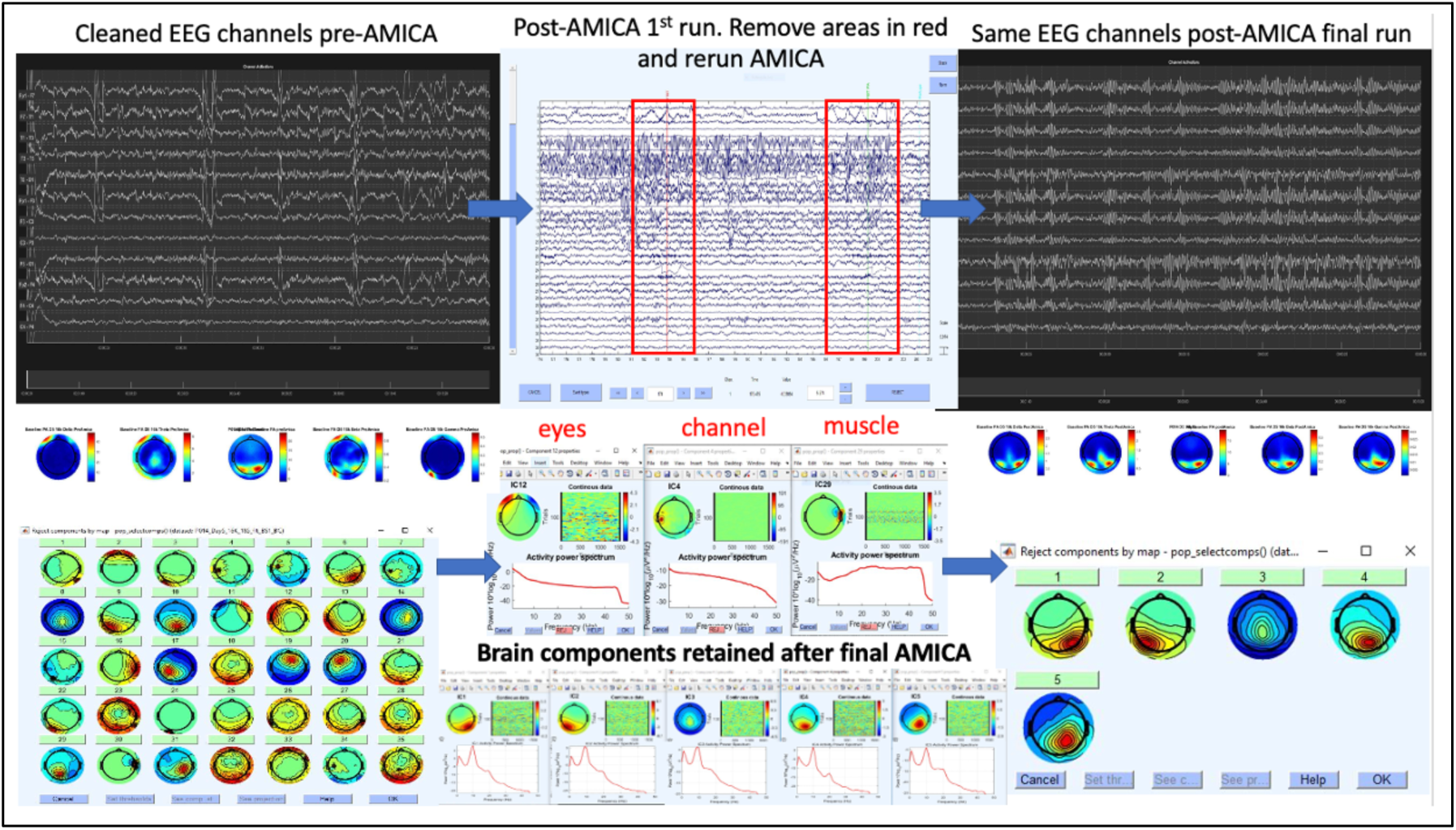
Clean epoch selection was aided by inspection of AMICA component time courses. Epochs where AMICA decomposition failed (i.e. when artifacts were found to contaminate many ICA components rather than remain contained in their respective muscle, eye or EKG components) were removed from data and AMICA was ran again. This allowed to obtained (right panel) smooth brain activity topographies, as expected by spatial filtering from the skill, and well separated from sparse and focal extracranial noise sources such as eye movements or muscle, with data quality assessment greatly favored by high spatial resolution of 256-electrode high density EEG.

**Suppl. Fig 2.**
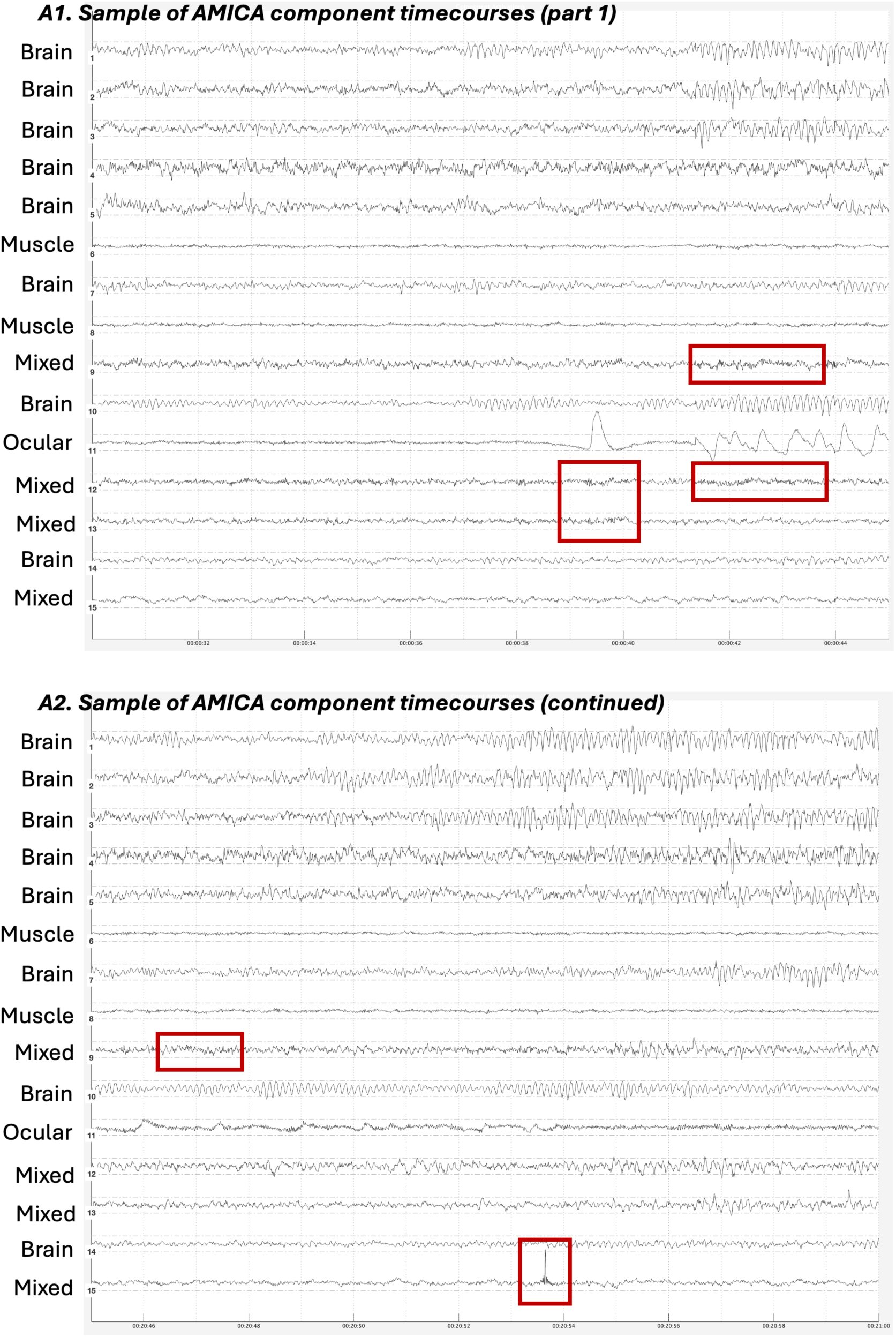
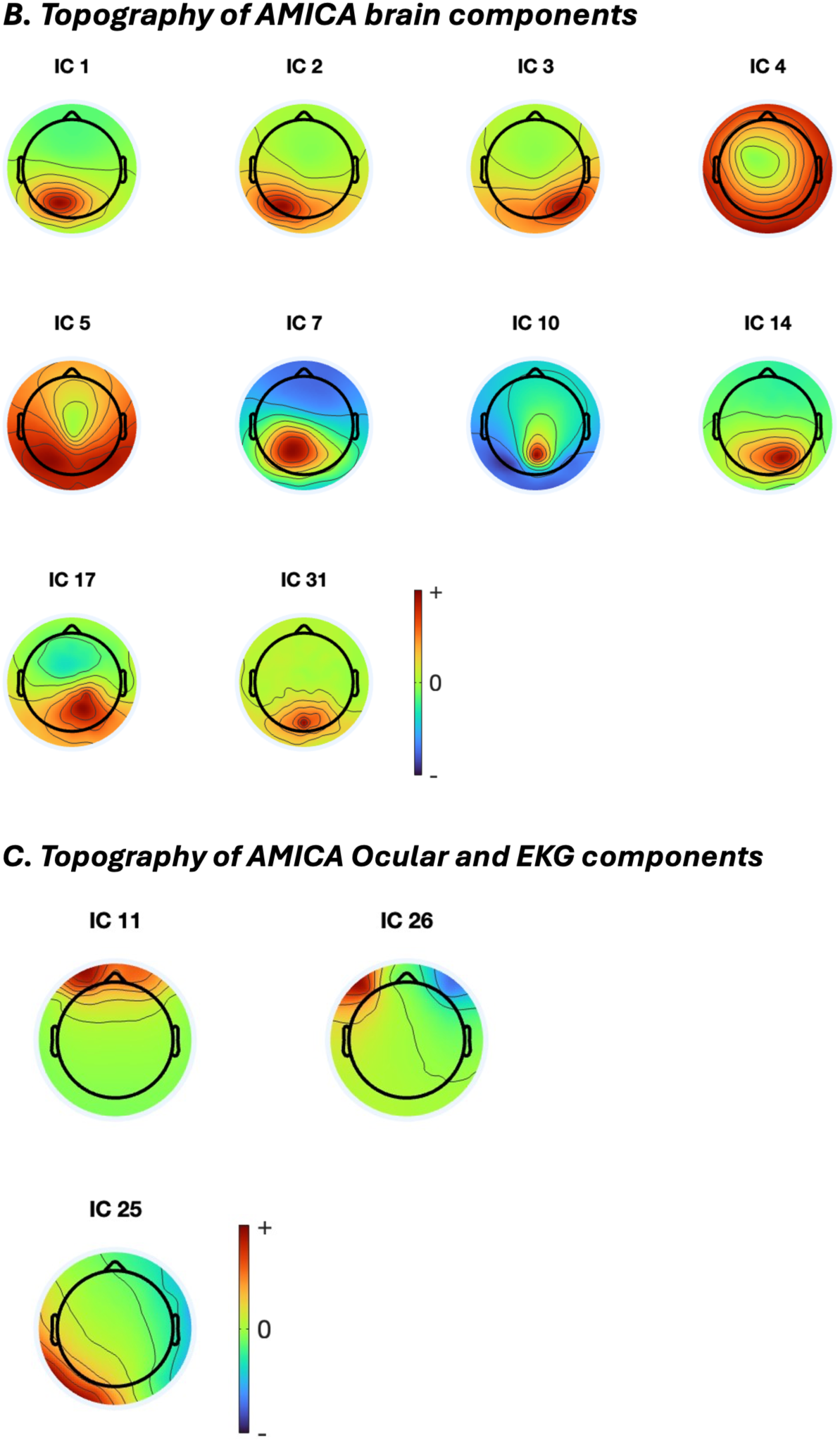
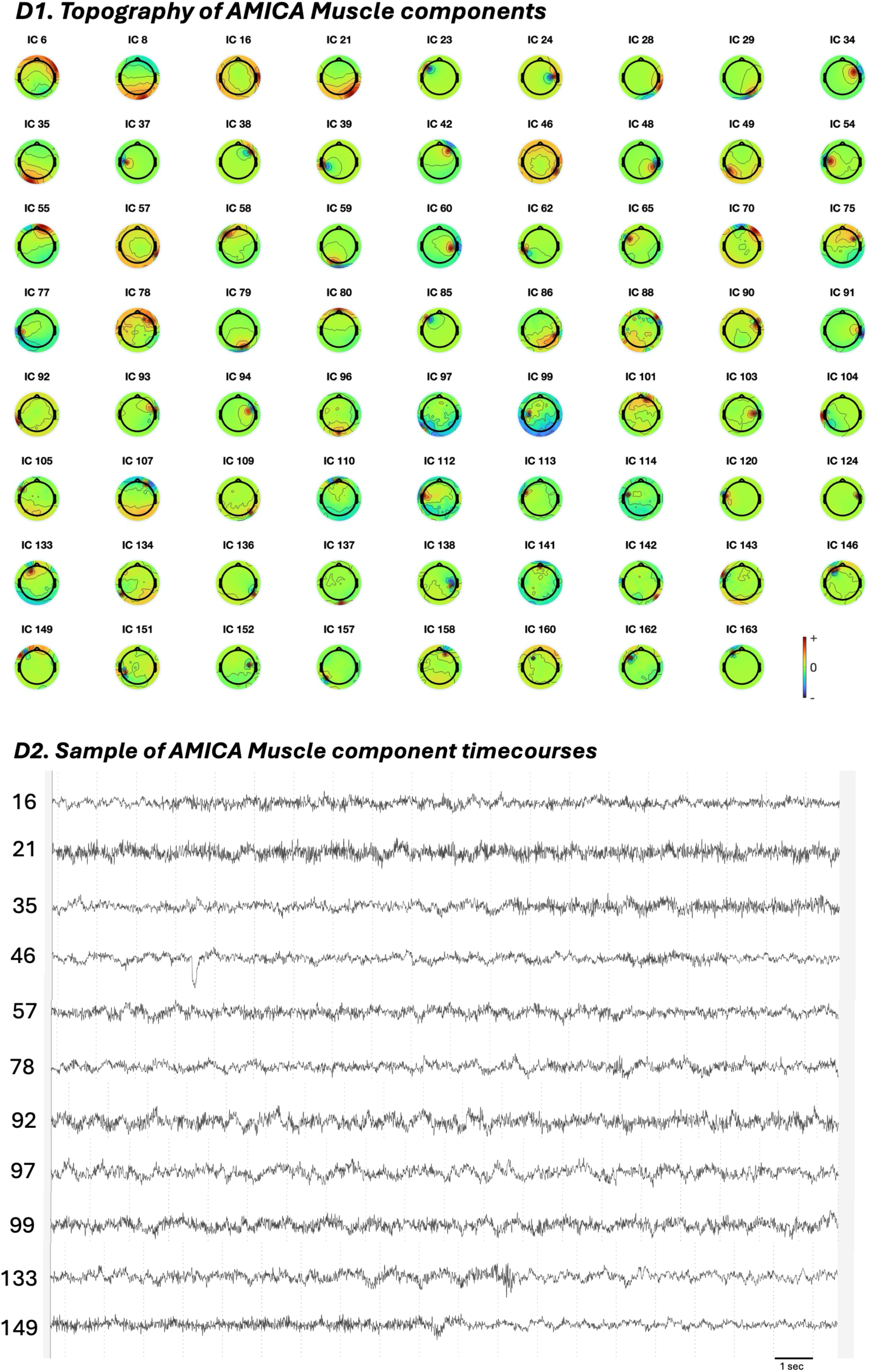
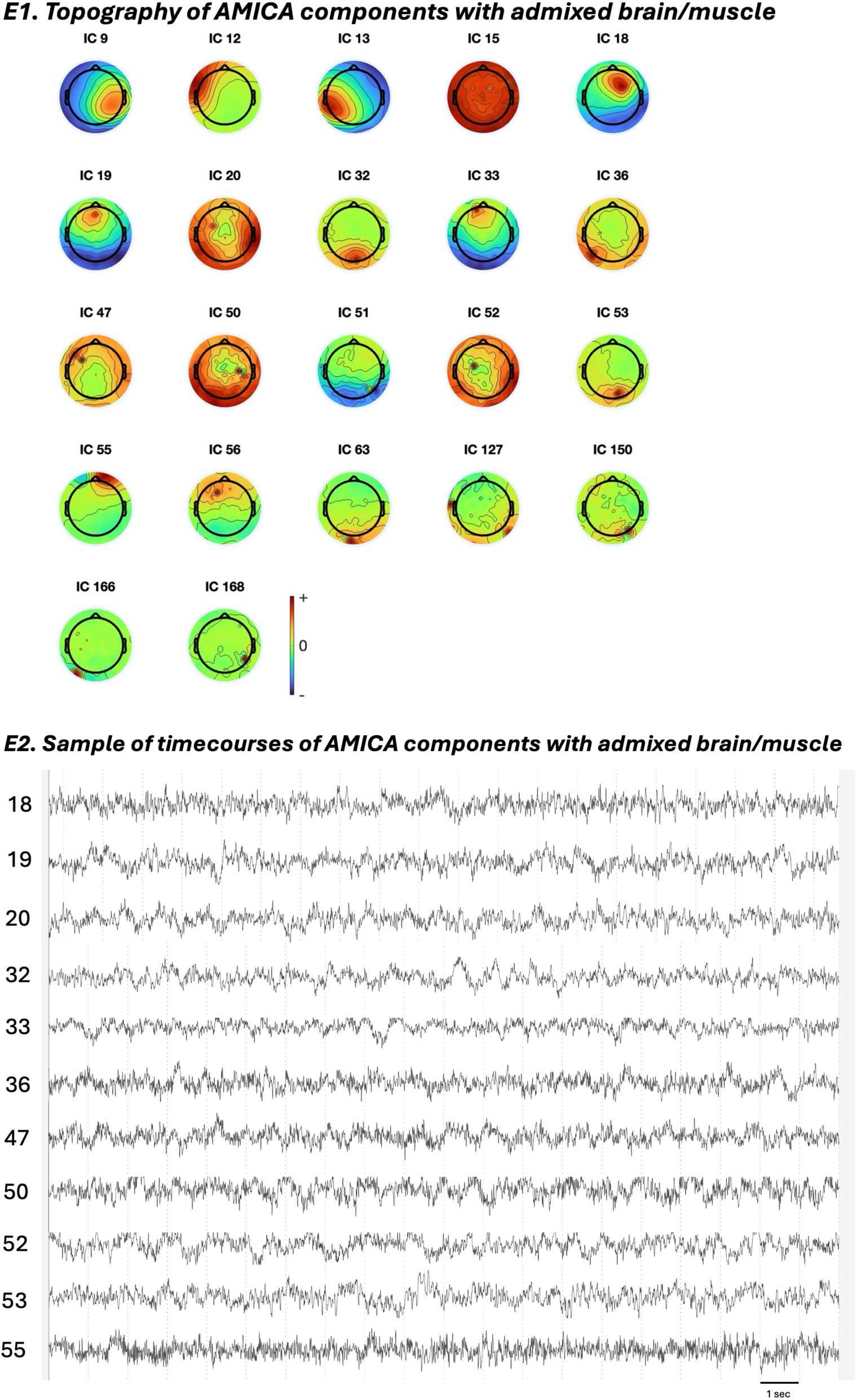
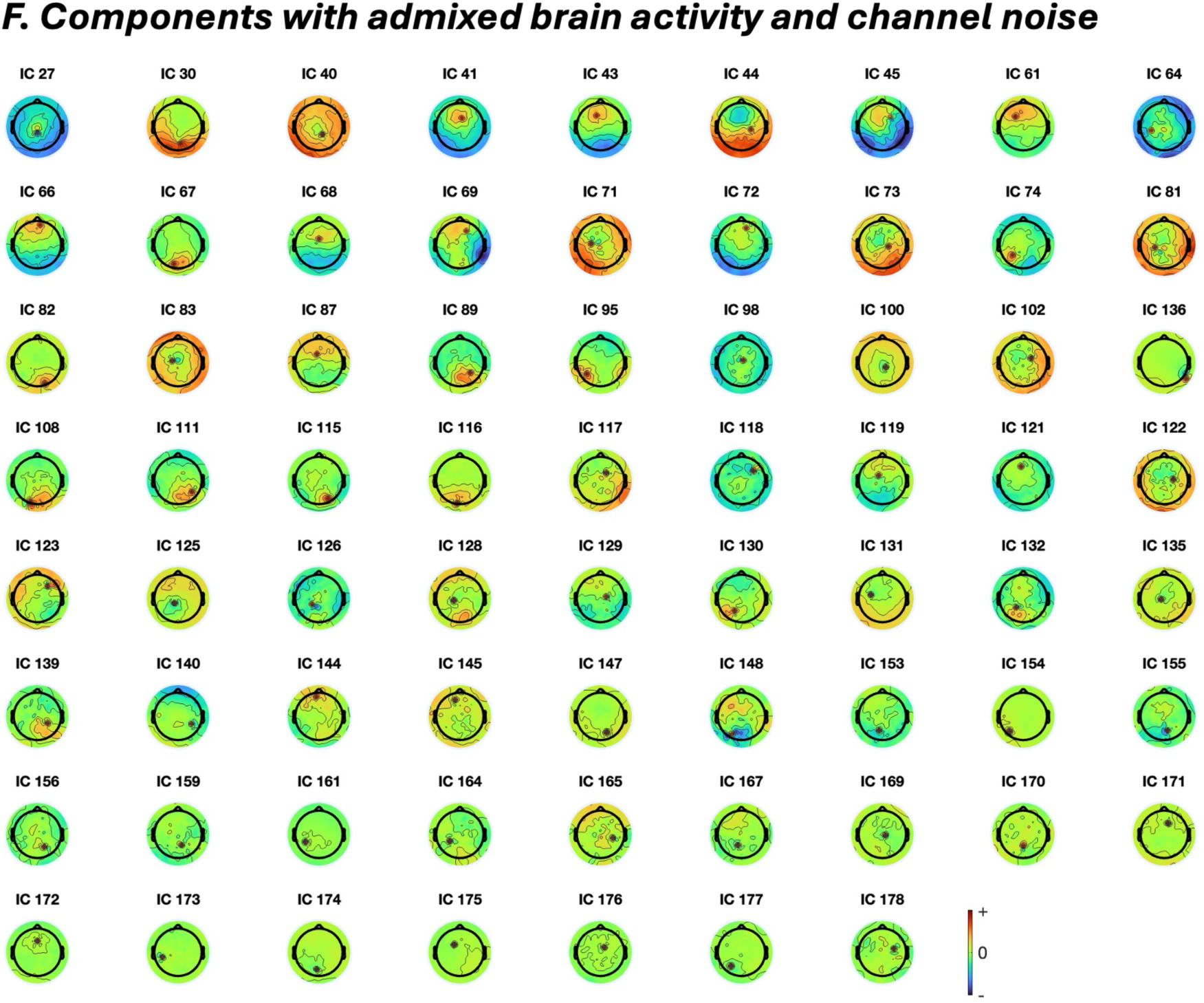
Example of enhanced ICA decomposition enabled by 256-electrode hdEEG combined with AMICA in a representative participant (Zen7). A1-2. Representative epochs of the first 15 components, capturing the highest variance, classified as Brain, Ocular, Muscle, or Mixed. Example of muscular artifacts contaminating brain activity within mixed components is highlighted in red. B. Topography of the ten AMICA Brain components included in the PP vs MW analysis. C. Topography of two Ocular and one EKG AMICA components identified. D1. Topography of 71 Muscle components identified. D2. Sample time courses of AMICA Muscle components. E1. Topography of AMICA components with admixed muscular and brain signals. E2. Sample time courses of AMICA components with admixed muscular and brain signals. F. Topography of AMICA components with admixed brain and channel noise signals.

**Suppl. Fig 3.**
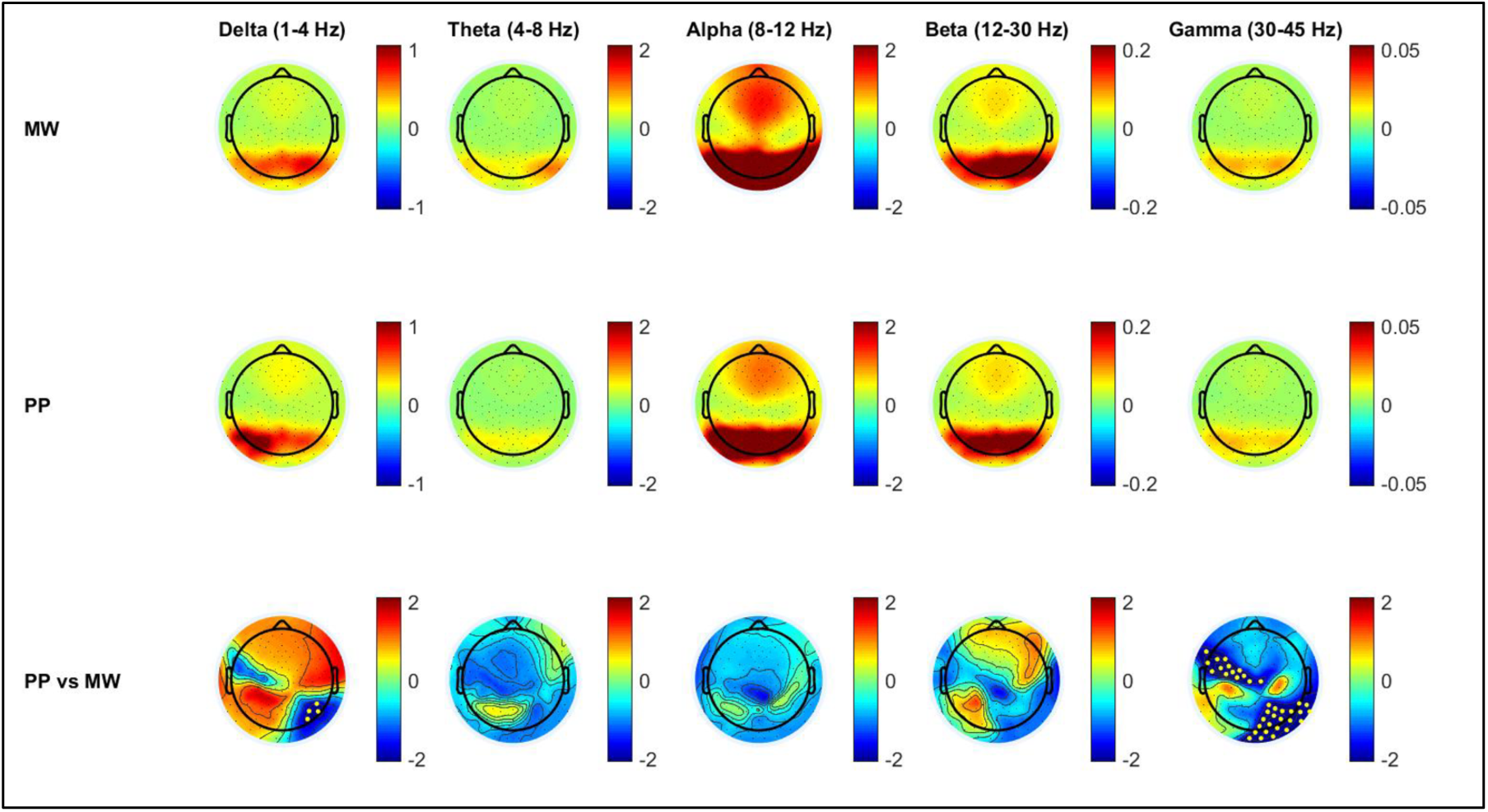
Neural correlates of pure presence (PP) compared to within-meditation mind-wandering (MW) – subgroup analysis in 7 Vajrayana LTMs in delta to gamma frequency ranges. First row: average group topography for MW. Second row: average group topography for PP. Third row: average group difference for PP compared to MW (within-subject subtraction of MW from PP). Last row: statistical non parametric mapping (SNPM) statistics for group difference between PP and MW. Colors represent EEG power (first to third rows) and T values (last row). Yellow dots represent channels surviving uncorrected p<0.05 values.

**Suppl. Fig 4.**
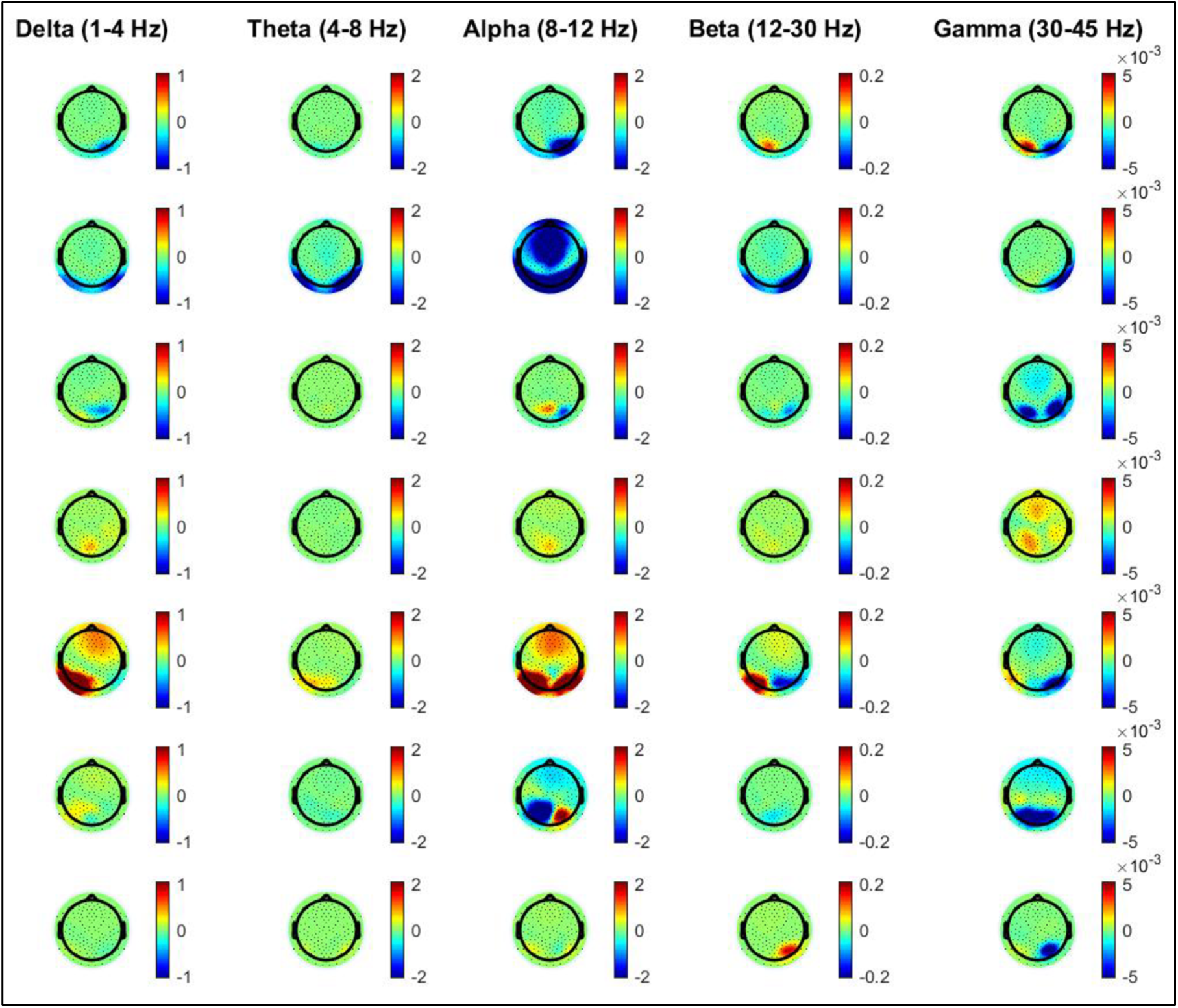
Neural correlates of pure presence (PP) compared to within-meditation mind-wandering (MW) – individual analysis in 7 Vajrayana LTMs in delta to gamma frequency ranges. Colors represent EEG power.

**Suppl. Fig 5.**
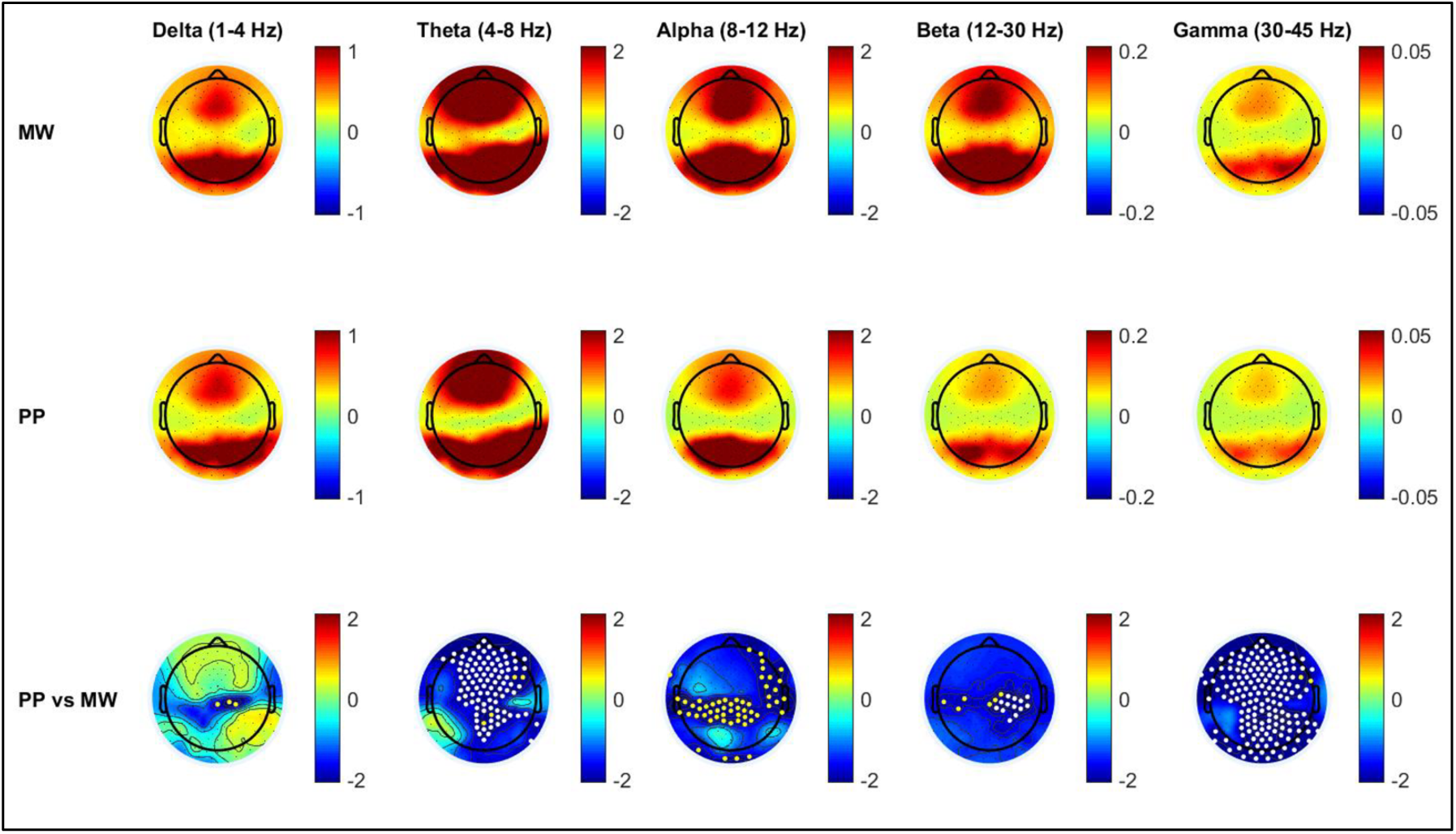
Neural correlates of pure presence (PP) compared to within-meditation mind-wandering (MW) – subgroup analysis in 8 Zen LTMs in delta to gamma frequency ranges. First row: average group topography for MW. Second row: average group topography for PP. Third row: average group difference for PP compared to MW (within-subject subtraction of MW from PP). Last row: statistical nonparametric mapping (SNPM) statistics for group difference between PP and MW. Colors represent EEG power (first to third rows) and T values (last row). Yellow dots represent channels surviving uncorrected p<0.05 values.

**Suppl. Fig 6.**
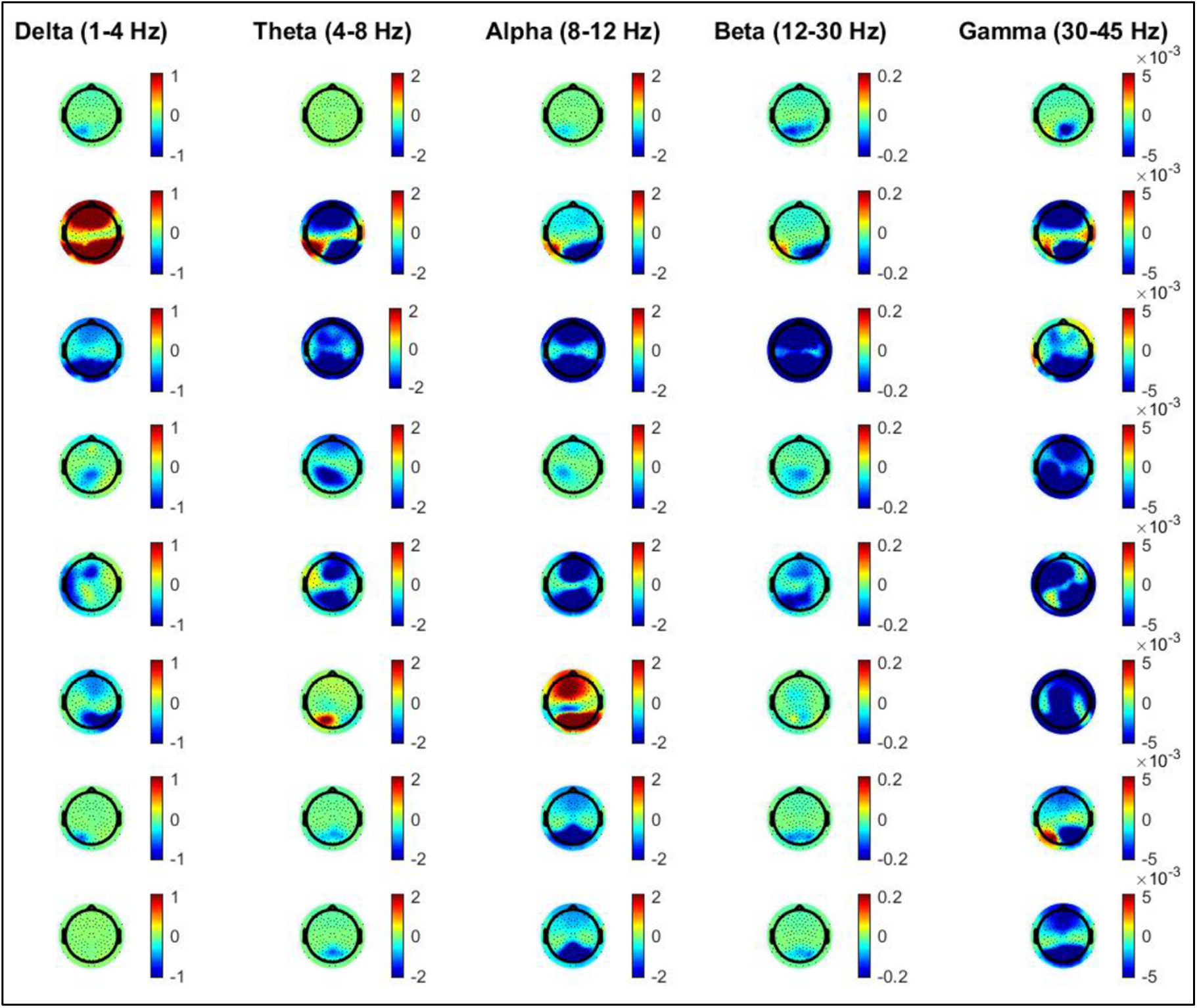
Neural correlates of pure presence (PP) compared to within-meditation mind-wandering (MW) – individual analysis in 8 Zen LTMs in delta to gamma frequency ranges. Colors represent EEG power.

**Suppl. Fig 7.**
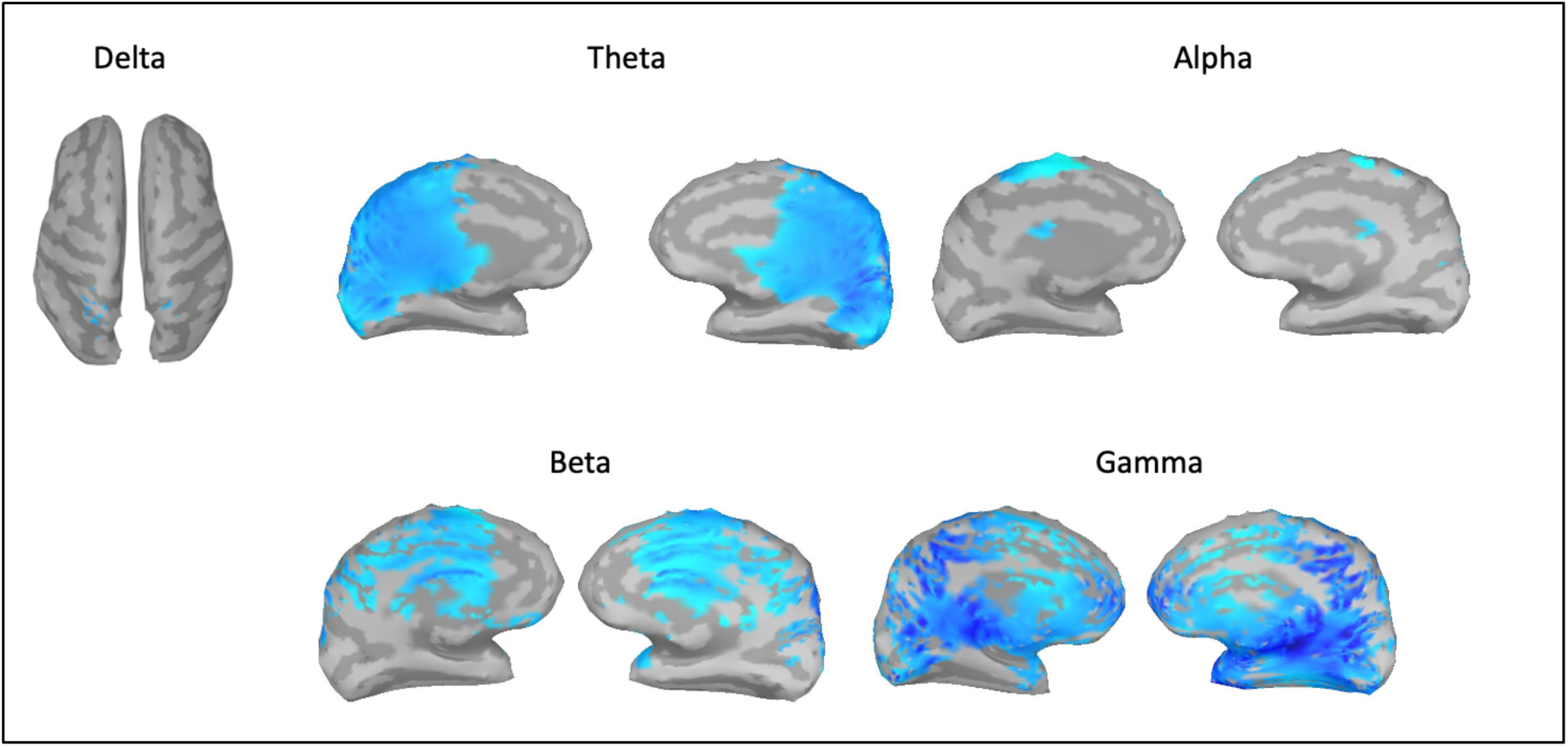
Source reconstruction results for gamma decrease in PP vs MW, uncorrected p<0.05. LL: left lateral; RL: right lateral; LM: left medial; RM: right medial. Colors represent T value for changes in source-level in delta (1-4 Hz), theta (4-8 Hz), alpha (8-12 Hz), beta (15-30 Hz) and gamma (30-45 Hz) power during PP compared to MW in the same 15 LTMs displayed in Figure 1. Darker blue colors represent stronger decreases.

**Suppl. Fig 8.**
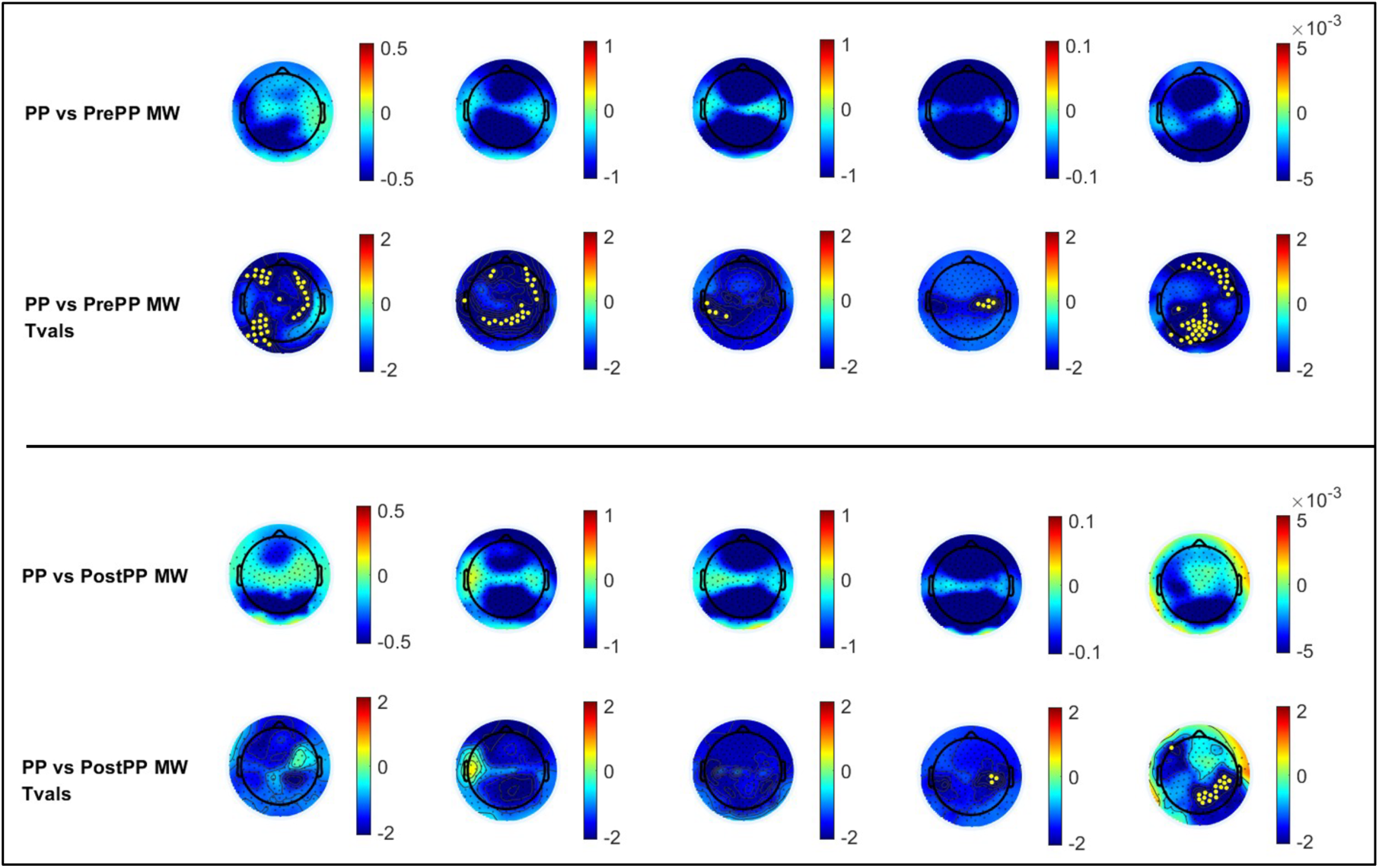
Comparisons of neural correlates of pure presence (PP) contrasted within-meditation mind-wandering (MW) occurring before vs after PP in Zen LTMs. Topographies are displayed in delta to gamma frequency ranges (same order and frequency bands as in main manuscript). Color scales represent EEG power (upper rows) and T values (lower rows). Yellow dots display electrodes surviving t-tests between PP and MW at uncorrected p<0.05.

**Suppl. Fig 9.**
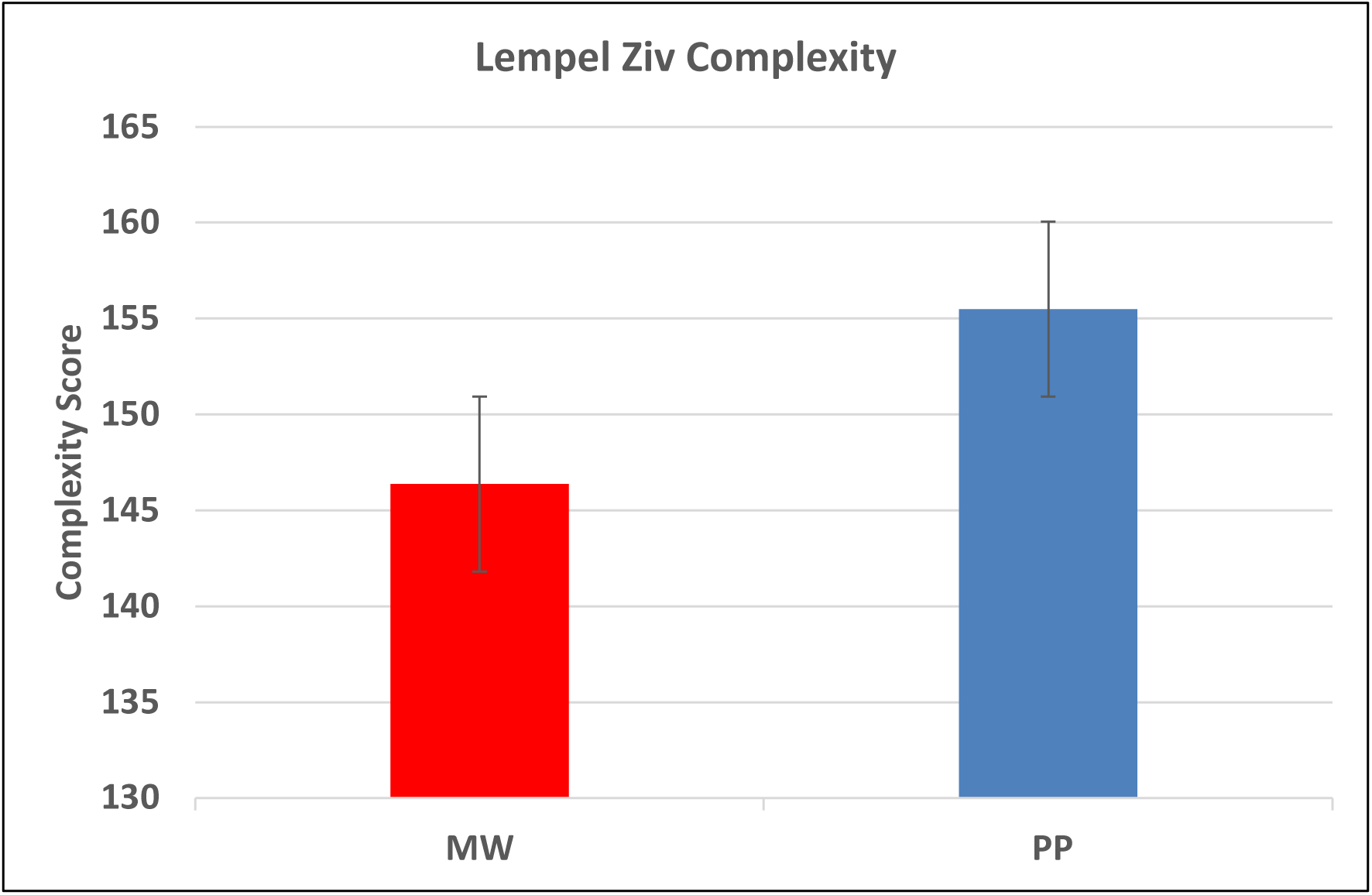
Comparisons of Lempel-Ziv complexity values during within-meditation mind-wandering (MW) vs. states of pure presence (PP). Values displayed are group mean and standard error of the mean in each condition. Statistical comparison with a t-test did not reach significance (p = 0.11).

**Suppl. Fig 10.**
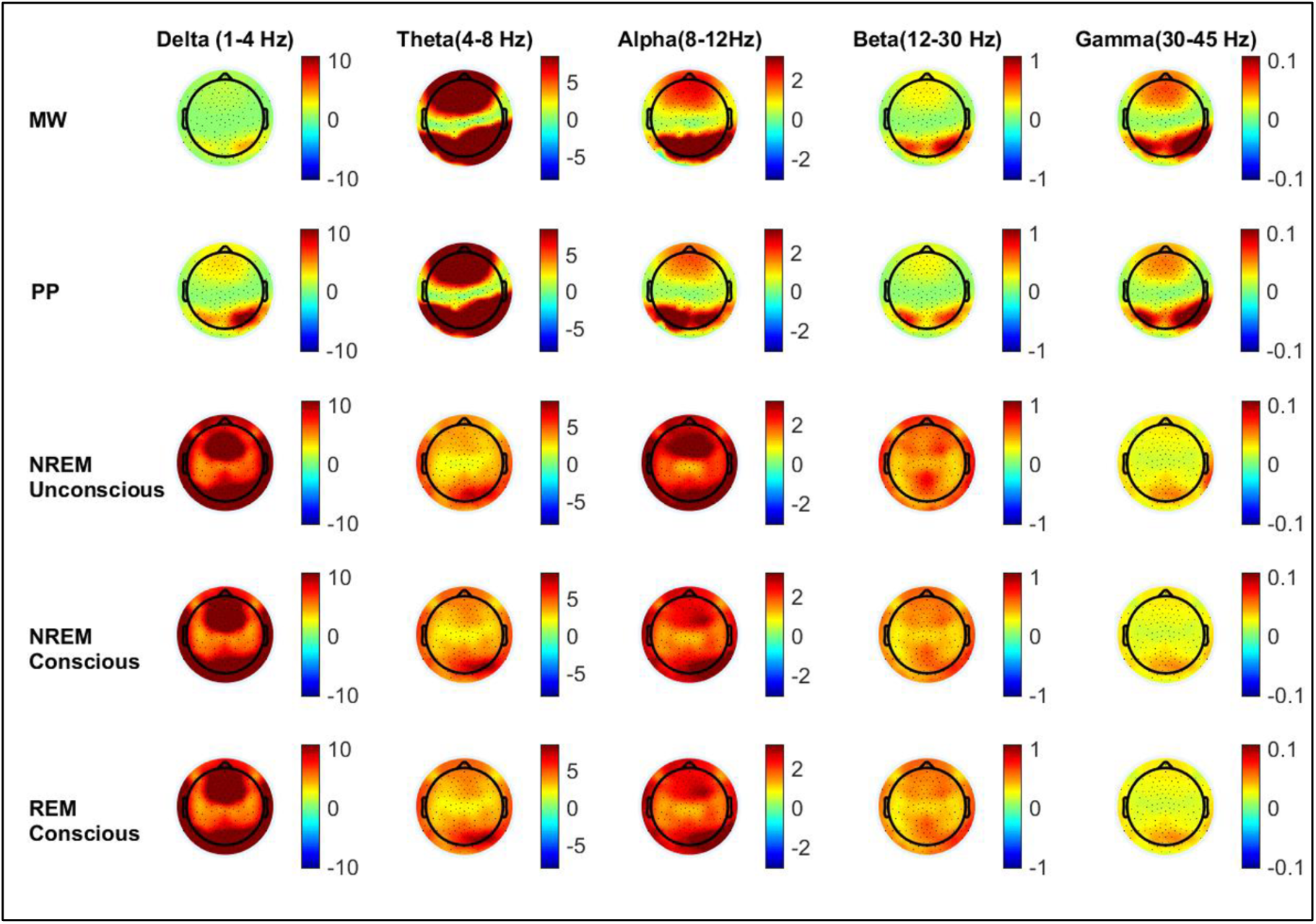
Individual analysis of a Zen LTM during MW, PP vs serial awakenings from NREM (either conscious or unconscious) and REM sleep (conscious). Colors represent EEG power.

**Suppl. Fig 11.**
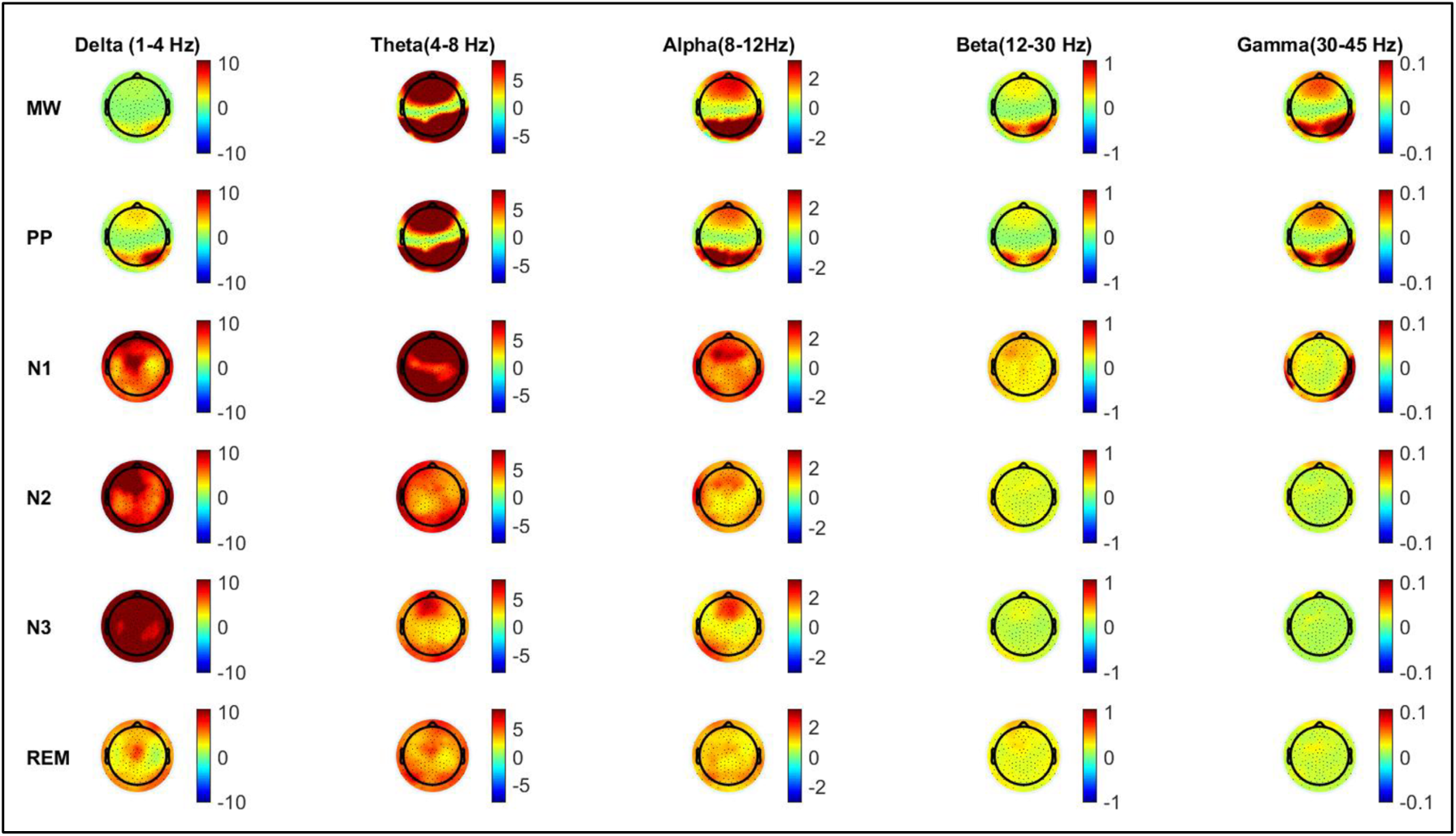
Individual analysis of a Zen LTM during MW, PP, NREM sleep stages 1, 2 and 3, and REM sleep. Colors represent EEG power.

**Suppl. Fig 12.**
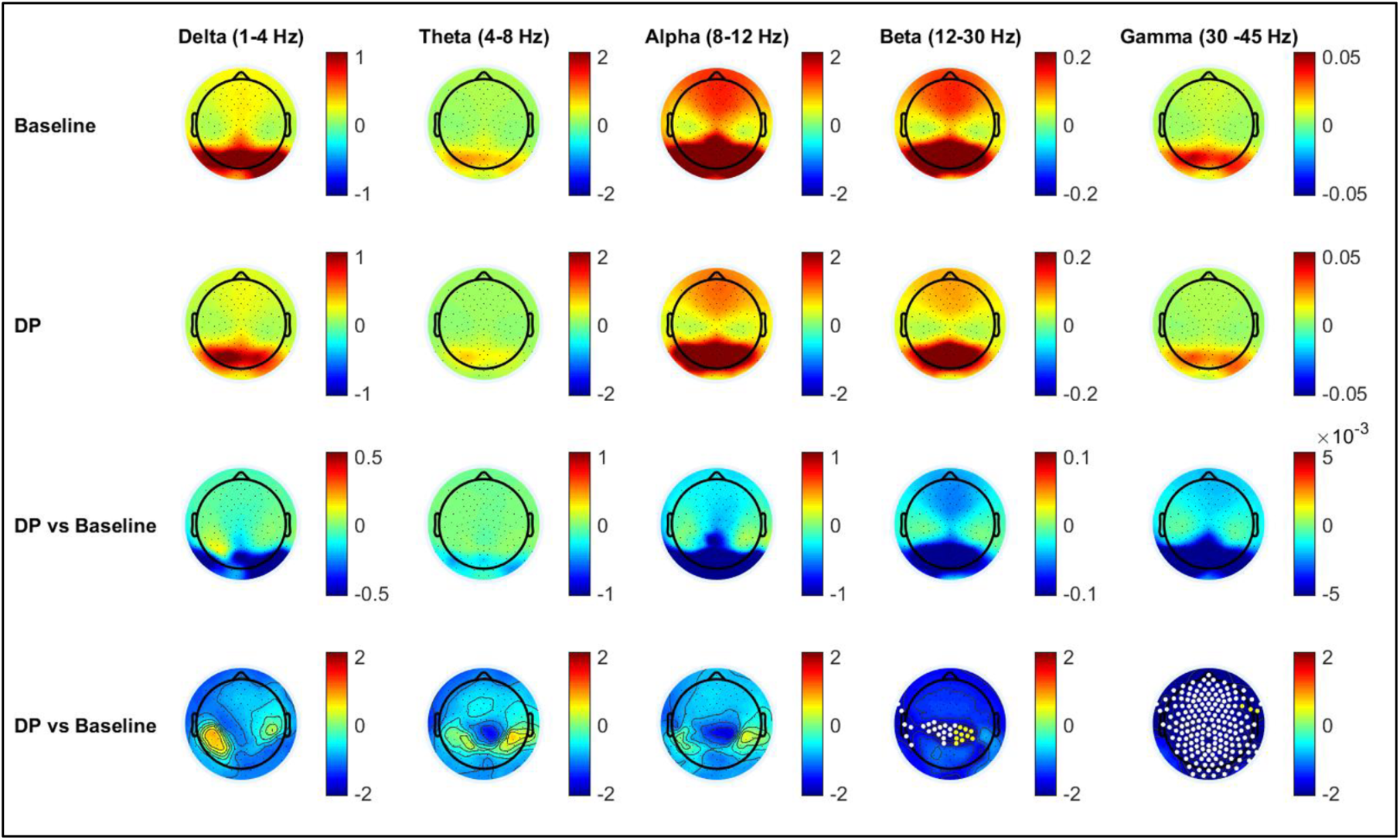
Neural correlates of pure presence (PP) using a no-report paradigm during the Dissolving phase (DP) of 16^th^ Karmapa Guru Yoga compared to baseline (11 Vajrayana LTMs). First row: average group topography for baseline. Second row: average group topography for DP. Third row: average group difference for DP compared to baseline (within-subject subtraction of baseline from DP). Last row: statistical non-parametric mapping (SNPM) statistics for group difference between DP and baseline. Colors represent EEG power (first to third rows) and T values (last row). White dots represent channels surviving p<0.05 corrected for multiple comparisons using SNPM; yellow dots represent channels surviving uncorrected p<0.05 values.

**Suppl. Fig 13.**
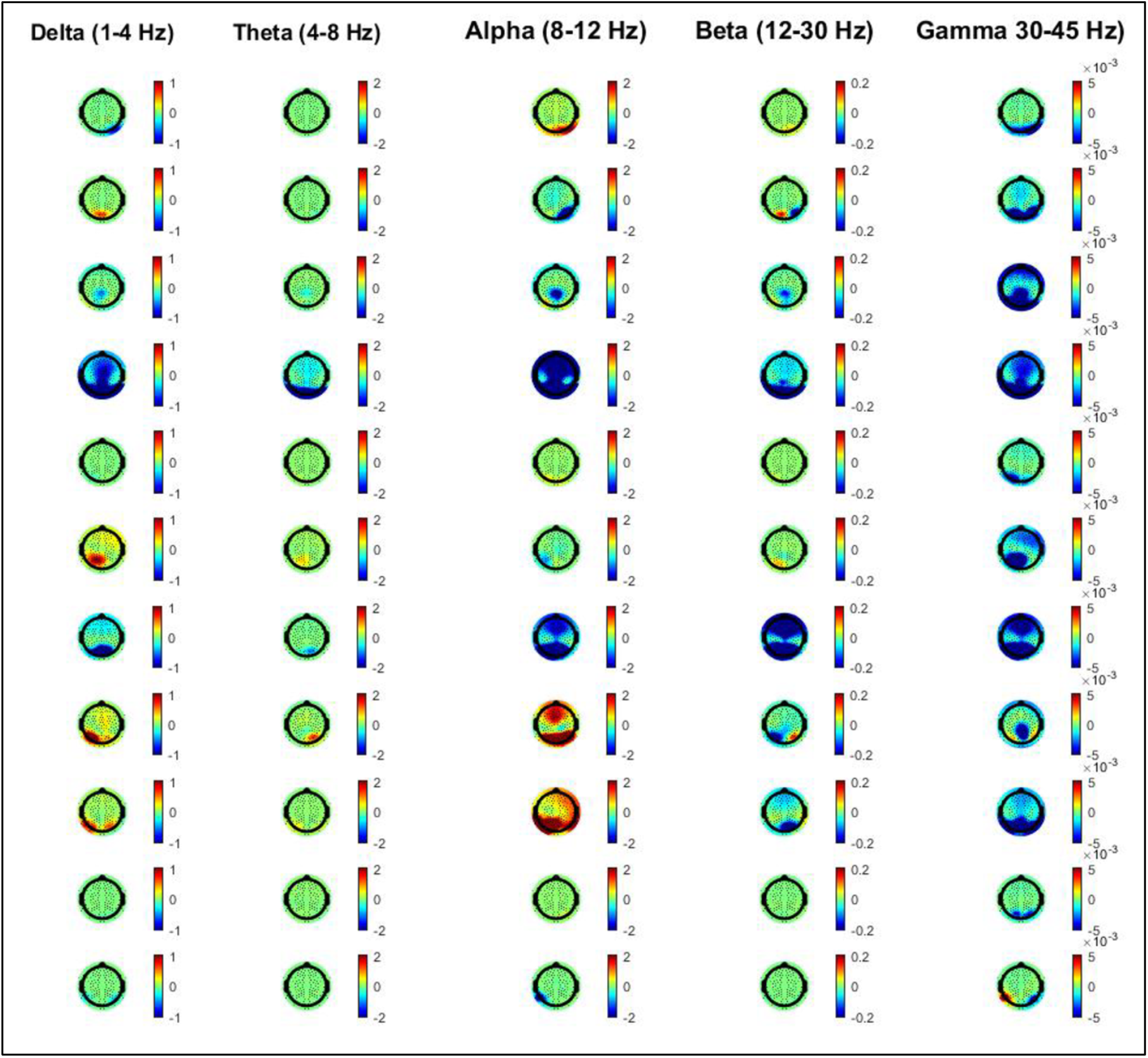
Neural correlates of dissolving phase (PP) compared to pre-meditation baseline – individual analysis in 11 Vajrayana LTMs in delta to gamma frequency ranges. Colors represent EEG power.

**Suppl. Figure 14.**
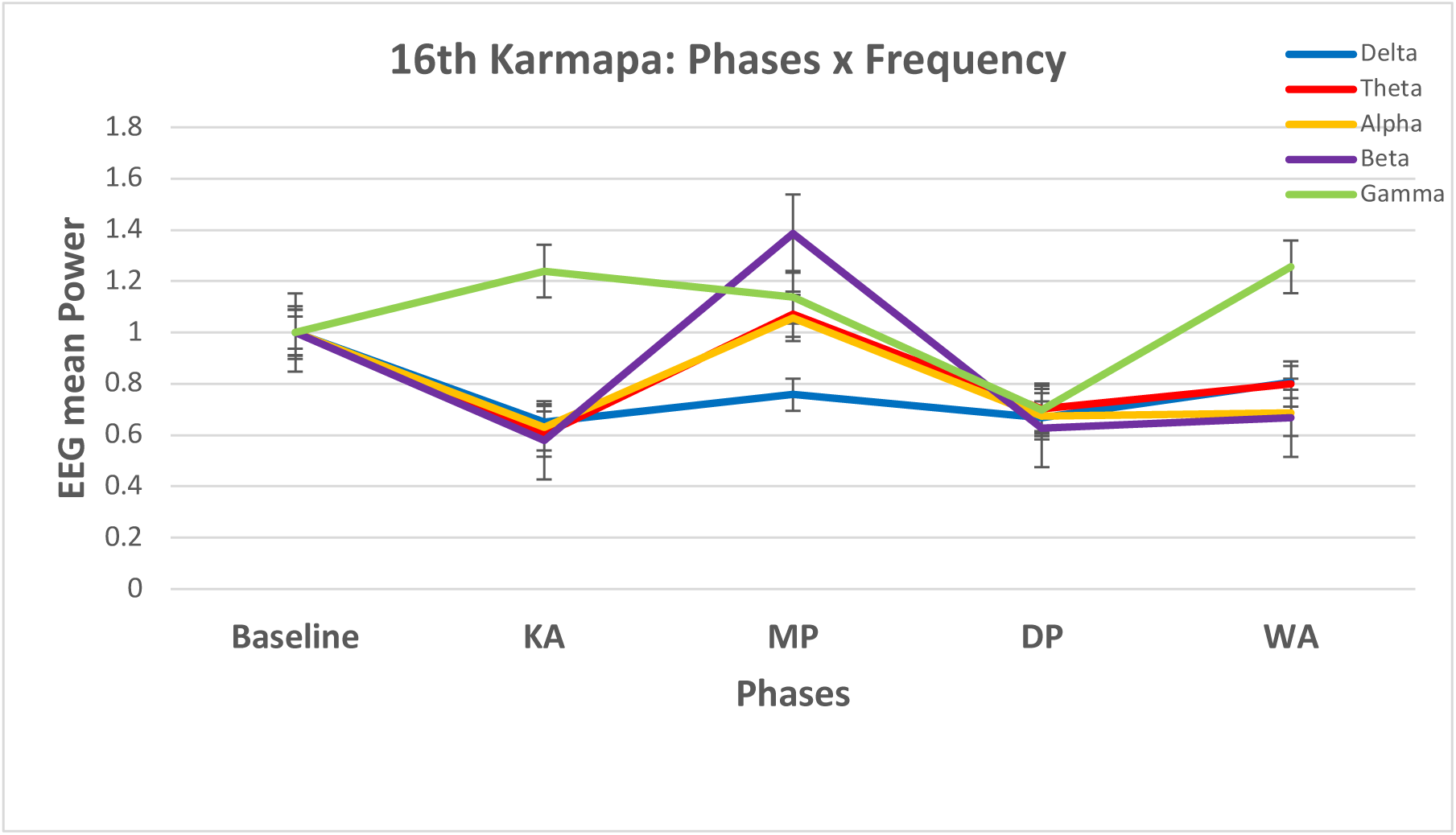
Temporal evolution of delta to gamma frequency bands during the 16^th^ Karmapa meditation phases (KA: Karmapa Appears; MP: Mantra Phase; DP: Dissolving Phase; WA: World Appears). Data represent group mean /- standard error of the mean for data of each phase normalized by baseline.

**Suppl. Fig. 15.**
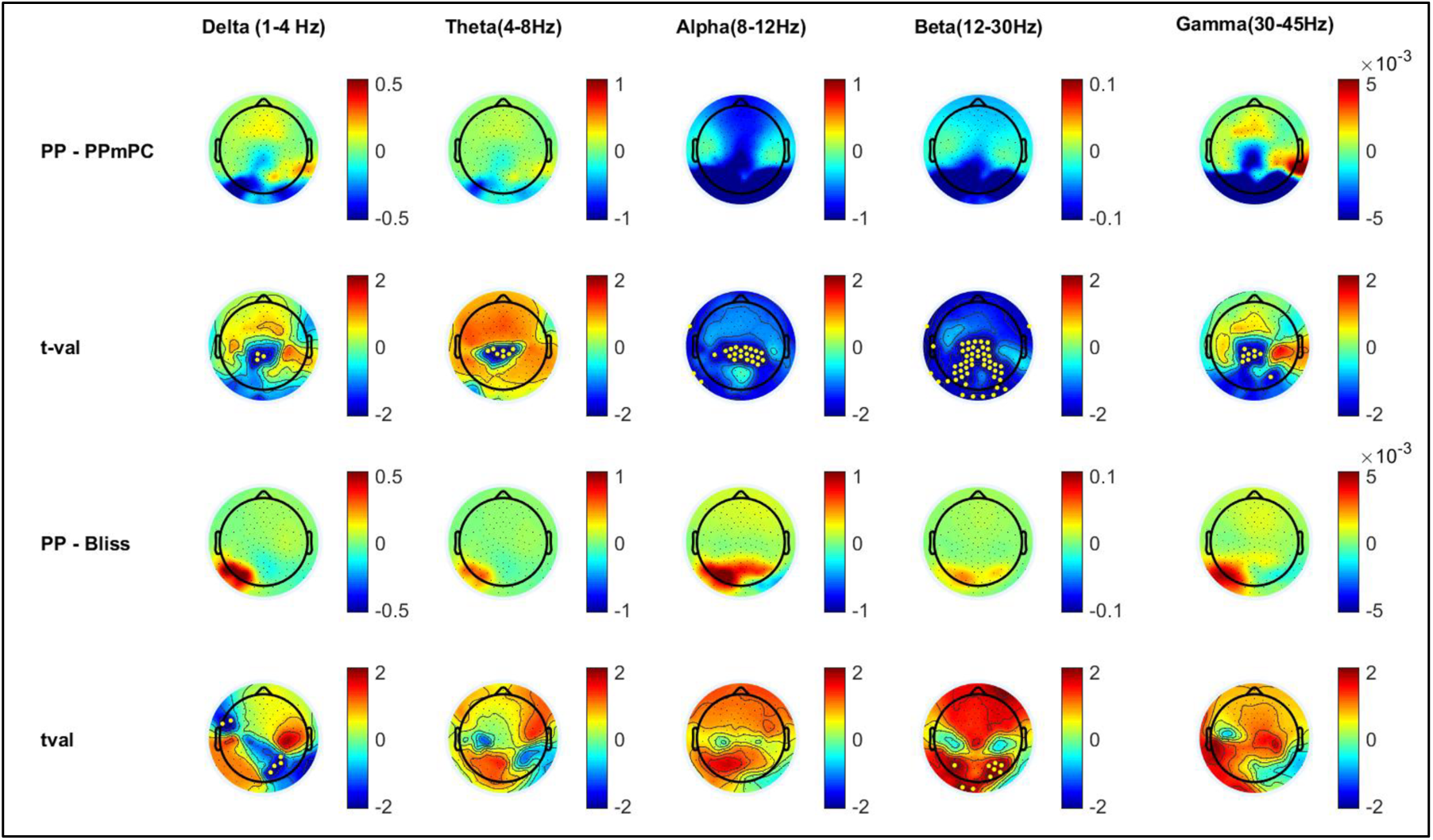
Upper two rows: comparison of PP vs PP-mPC in 15 subjects (uncorrected p<0.05). PP displays decreased delta and decreased gamma in central, posteromedial and occipital cortices. Lower two rows: comparison of PP vs PP-bliss in 9 subjects (uncorrected p<0.05). PP displays slightly less delta than PP-bliss, but no significant changes in gamma are seen. Colors represent EEG power (first and third rows) and T values (second and fourth rows). Yellow dots represent channels surviving uncorrected p<0.05 values.

**Suppl. Fig 16.**
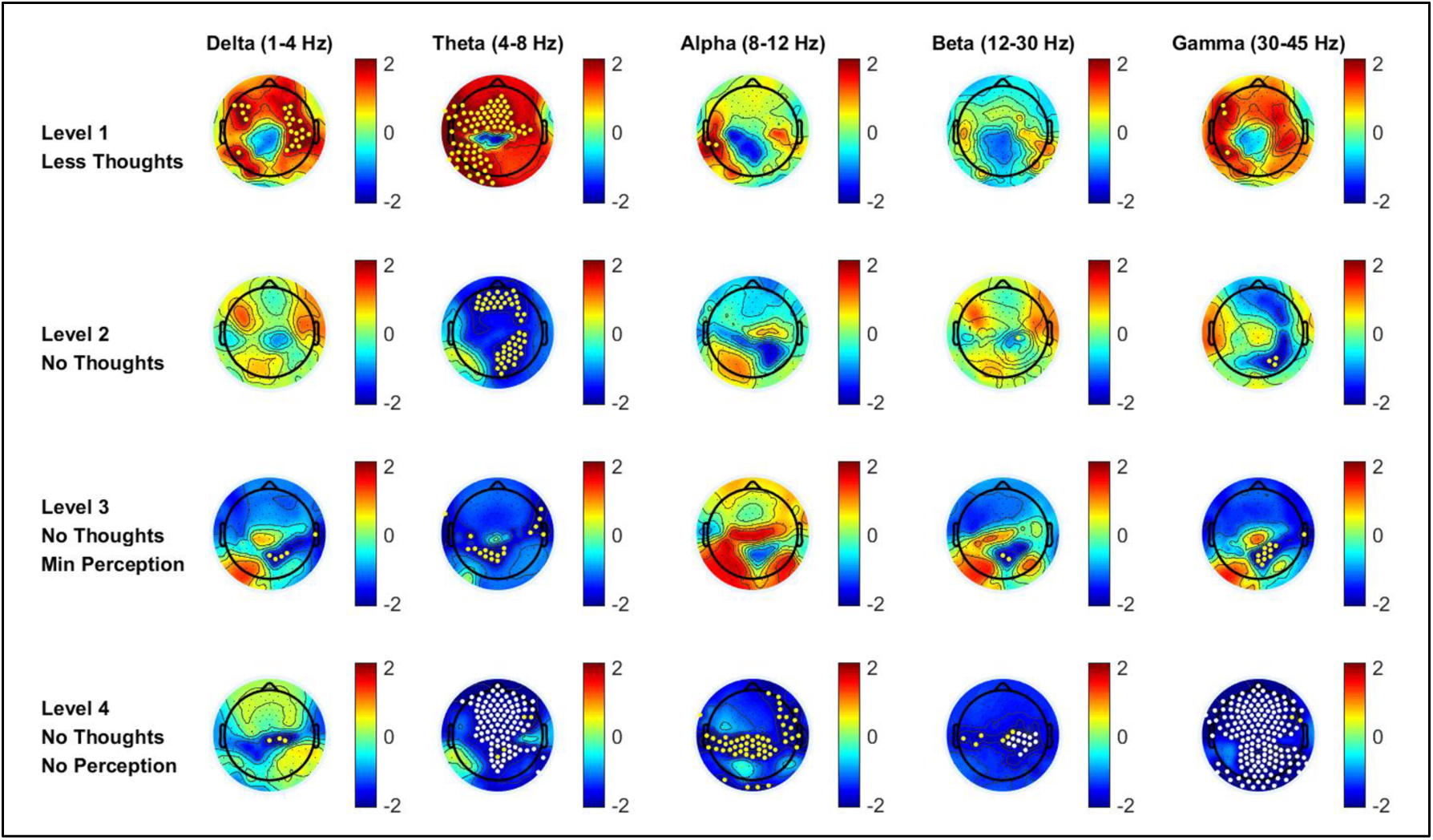
Progressive decrease in gamma with levels of absorption in 8 Zen LTMs (no thoughts, less thoughts, PP-mC then PP). Colors represent T values. Yellow dots represent channels surviving uncorrected p<0.05 values. White dots represent channels surviving p<0.05 corrected for multiple comparisons using SNPM; yellow dots represent channels surviving uncorrected p<0.05 values.

## Supplementary Methods

### 1. Zen Science–Meditation Retreat Schedule

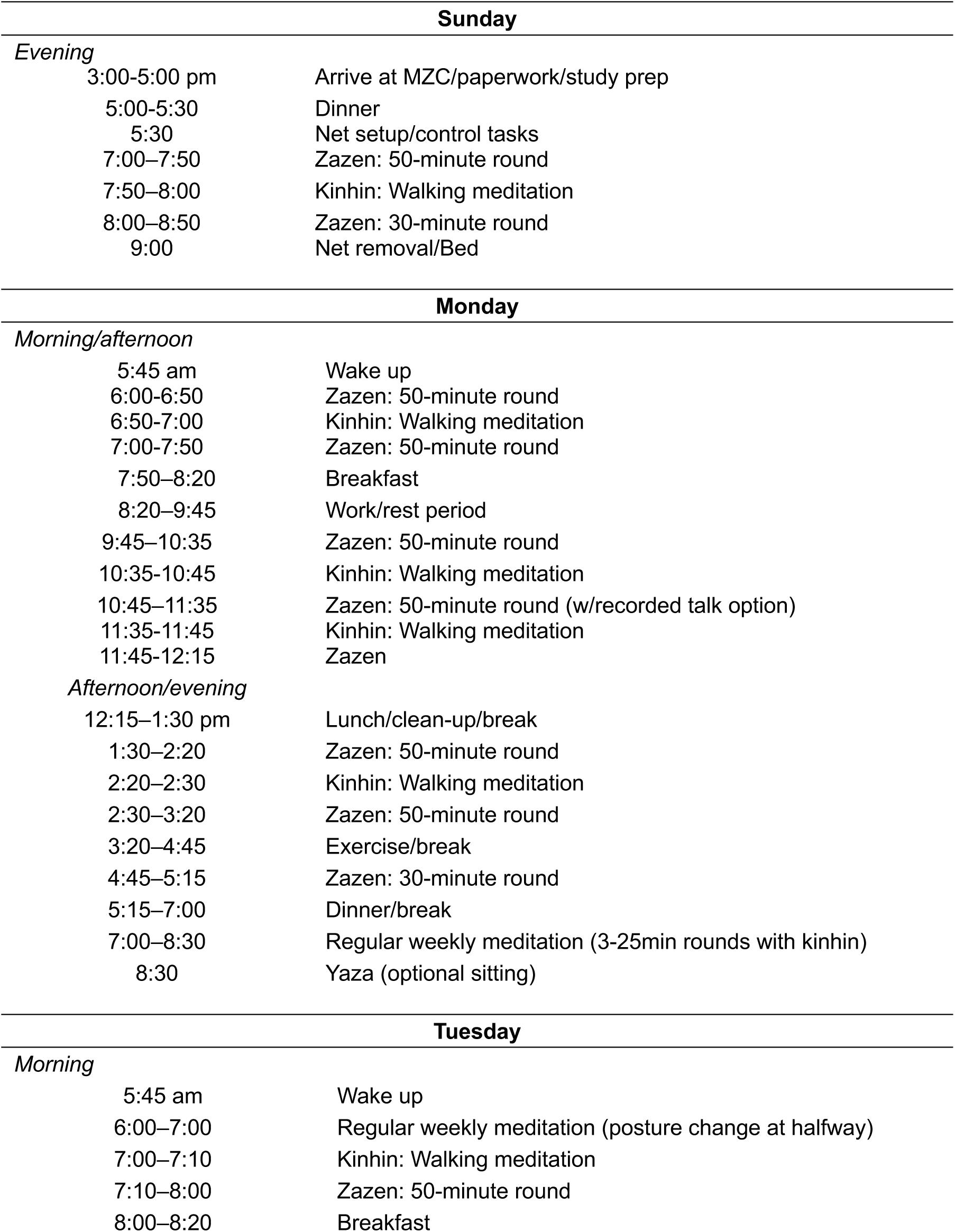
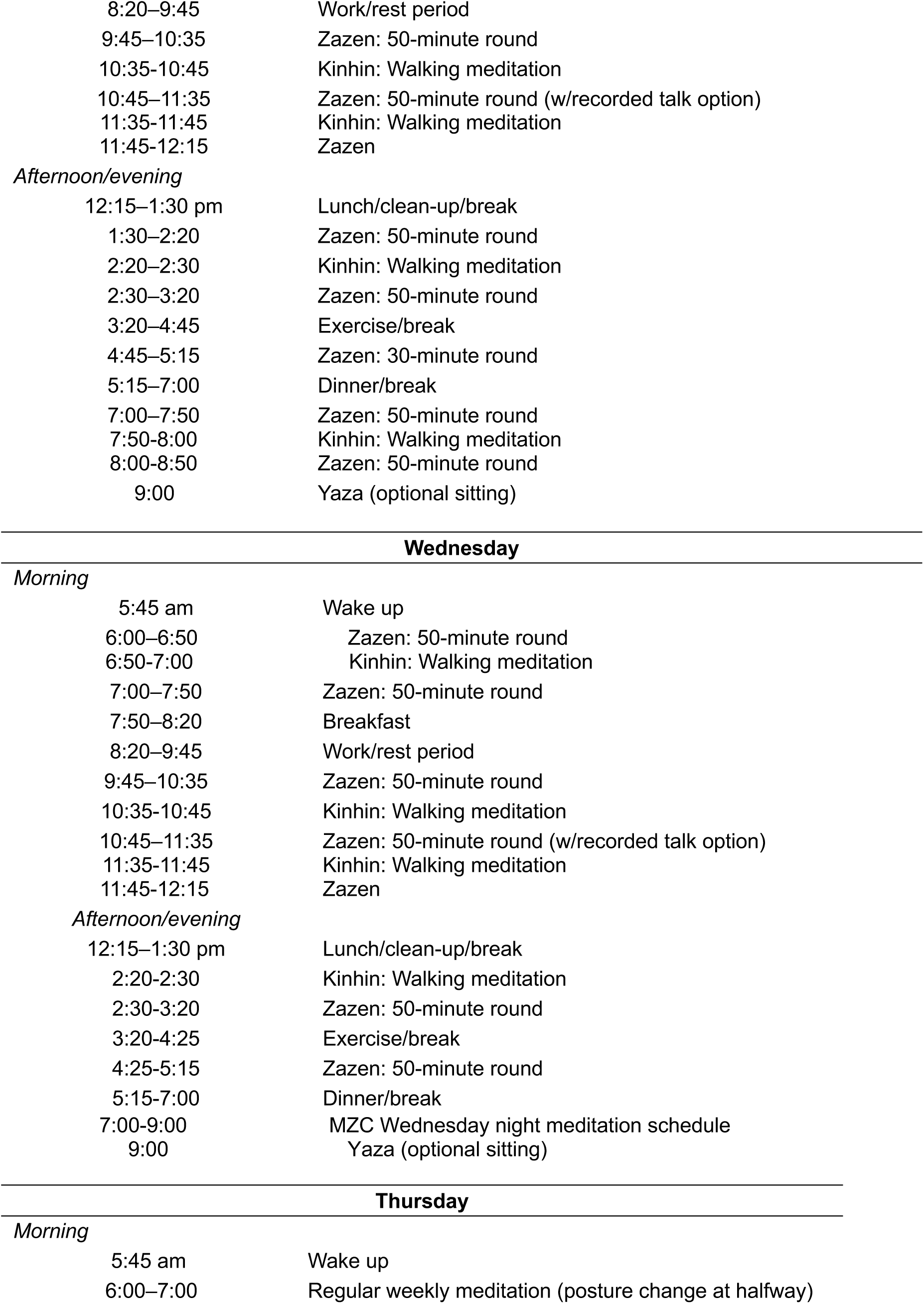
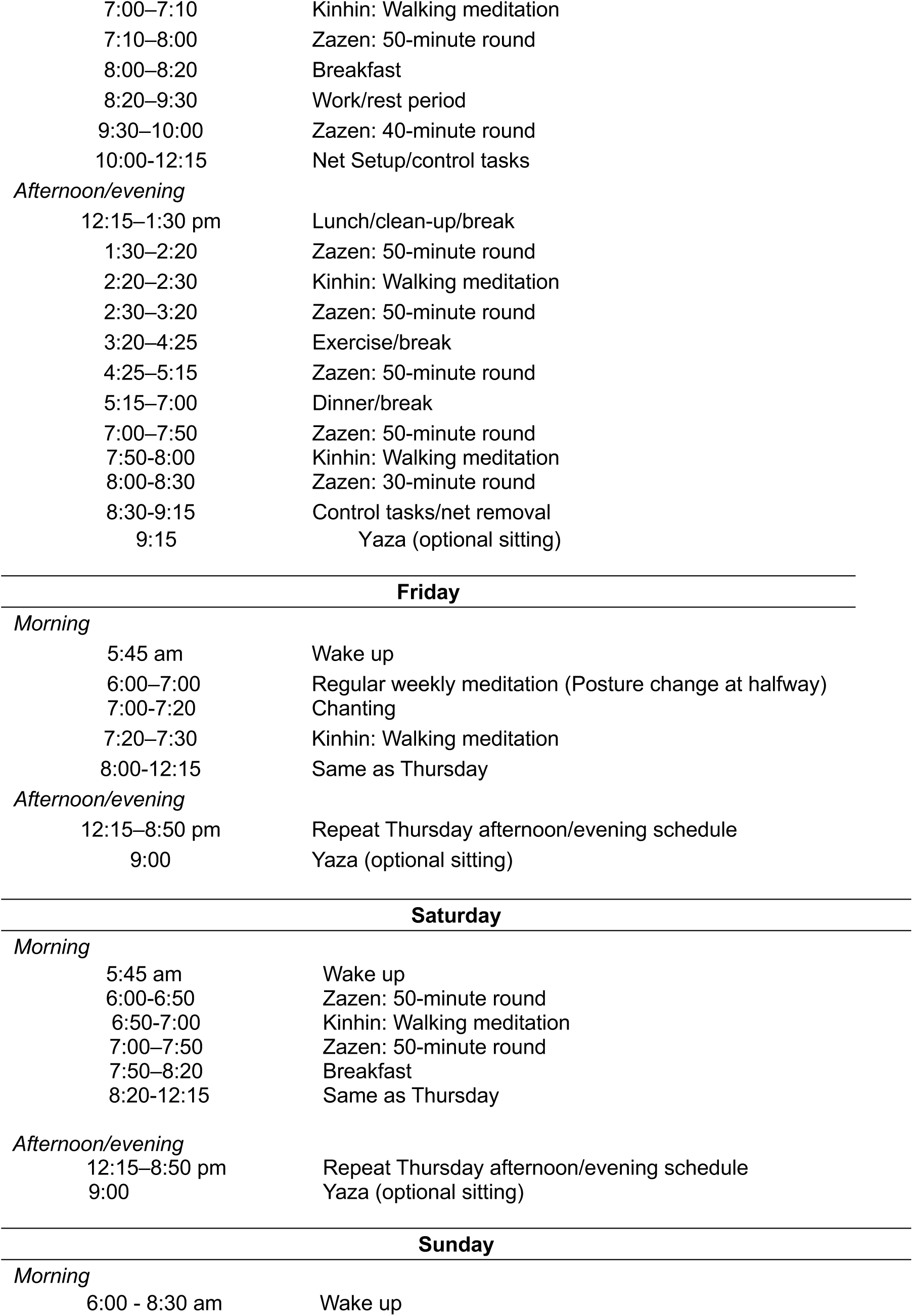
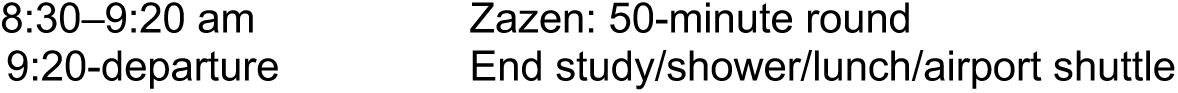

### 2. Vajrayana Science–Meditation Retreat Schedule

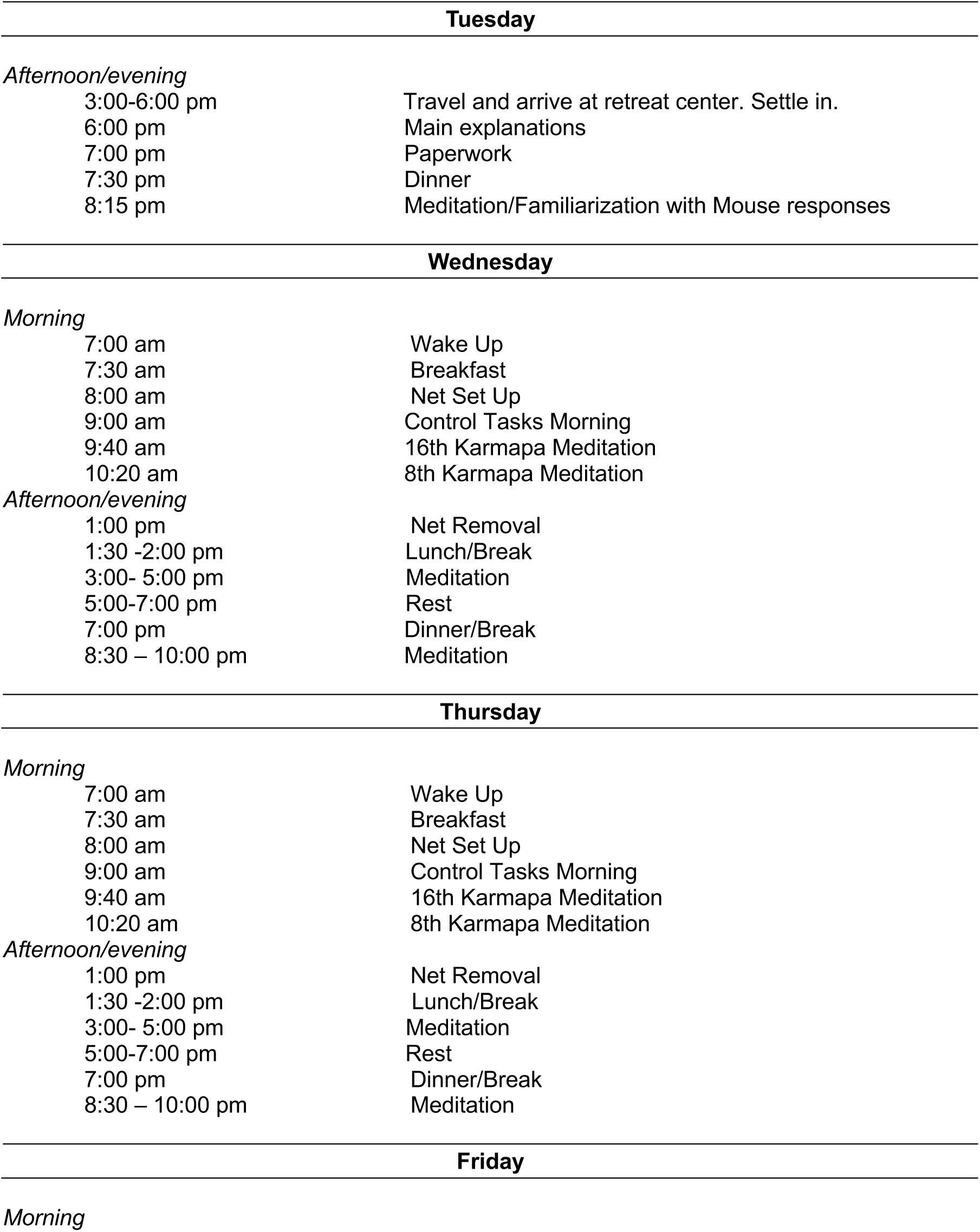
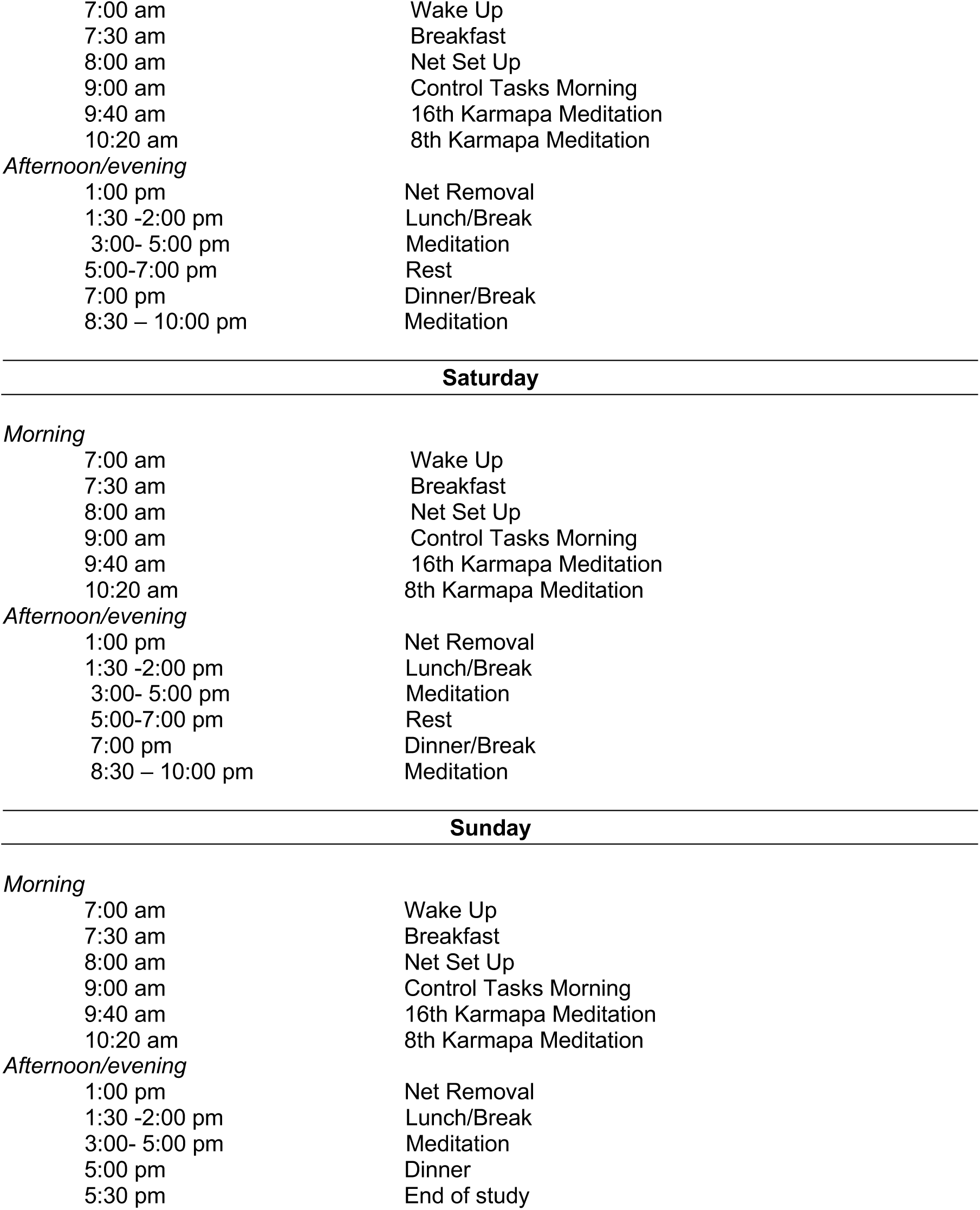

| Tuesday |  |
| --- | --- |
| *Afternoon/evening* |  |
| 3:00-6:00 pm | Travel and arrive at retreat center. Settle in. |
| 6:00 pm | Main explanations |
| 7:00 pm | Paperwork |
| 7:30 pm | Dinner |
| 8:15 pm | Meditation/Familiarization with Mouse responses |
| Wednesday |  |
| *Morning* |  |
| 7:00 am | Wake Up |
| 7:30 am | Breakfast |
| 8:00 am | Net Set Up |
| 9:00 am | Control Tasks Morning |
| 9:40 am | 16th Karmapa Meditation |
| 10:20 am | 8th Karmapa Meditation |
| *Afternoon/evening* |  |
| 1:00 pm | Net Removal |
| 1:30 -2:00 pm | Lunch/Break |
| 3:00- 5:00 pm | Meditation |
| 5:00-7:00 pm | Rest |
| 7:00 pm | Dinner/Break |
| 8:30 – 10:00 pm | Meditation |
| Thursday |  |
| *Morning* |  |
| 7:00 am | Wake Up |
| 7:30 am | Breakfast |
| 8:00 am | Net Set Up |
| 9:00 am | Control Tasks Morning |
| 9:40 am | 16th Karmapa Meditation |
| 10:20 am | 8th Karmapa Meditation |
| *Afternoon/evening* |  |
| 1:00 pm | Net Removal |
| 1:30 -2:00 pm | Lunch/Break |
| 3:00- 5:00 pm | Meditation |
| 5:00-7:00 pm | Rest |
| 7:00 pm | Dinner/Break |
| 8:30 – 10:00 pm | Meditation |
| Friday |  |
| *Morning* |  |
| 7:00 am | Wake Up |
| 7:30 am | Breakfast |
| 8:00 am | Net Set Up |
| 9:00 am | Control Tasks Morning |
| 9:40 am | 16th Karmapa Meditation |
| 10:20 am | 8th Karmapa Meditation |
| *Afternoon/evening* |  |
| 1:00 pm | Net Removal |
| 1:30 -2:00 pm | Lunch/Break |
| 3:00- 5:00 pm | Meditation |
| 5:00-7:00 pm | Rest |
| 7:00 pm | Dinner/Break |
| 8:30 – 10:00 pm | Meditation |
| Saturday |  |
| *Morning* |  |
| 7:00 am | Wake Up |
| 7:30 am | Breakfast |
| 8:00 am | Net Set Up |
| 9:00 am | Control Tasks Morning |
| 9:40 am | 16th Karmapa Meditation |
| 10:20 am | 8th Karmapa Meditation |
| *Afternoon/evening* |  |
| 1:00 pm | Net Removal |
| 1:30 -2:00 pm | Lunch/Break |
| 3:00- 5:00 pm | Meditation |
| 5:00-7:00 pm | Rest |
| 7:00 pm | Dinner/Break |
| 8:30 – 10:00 pm | Meditation |
| Sunday |  |
| *Morning* |  |
| 7:00 am | Wake Up |
| 7:30 am | Breakfast |
| 8:00 am | Net Set Up |
| 9:00 am | Control Tasks Morning |
| 9:40 am | 16th Karmapa Meditation |
| 10:20 am | 8th Karmapa Meditation |
| *Afternoon/evening* |  |
| 1:00 pm | Net Removal |
| 1:30 -2:00 pm | Lunch/Break |
| 3:00- 5:00 pm | Meditation |
| 5:00 pm | Dinner |
| 5:30 pm | End of study |

### 3. Questionnaires administered to Zen and Vajrayana (Diamond Way Buddhism, DWB) LTMs

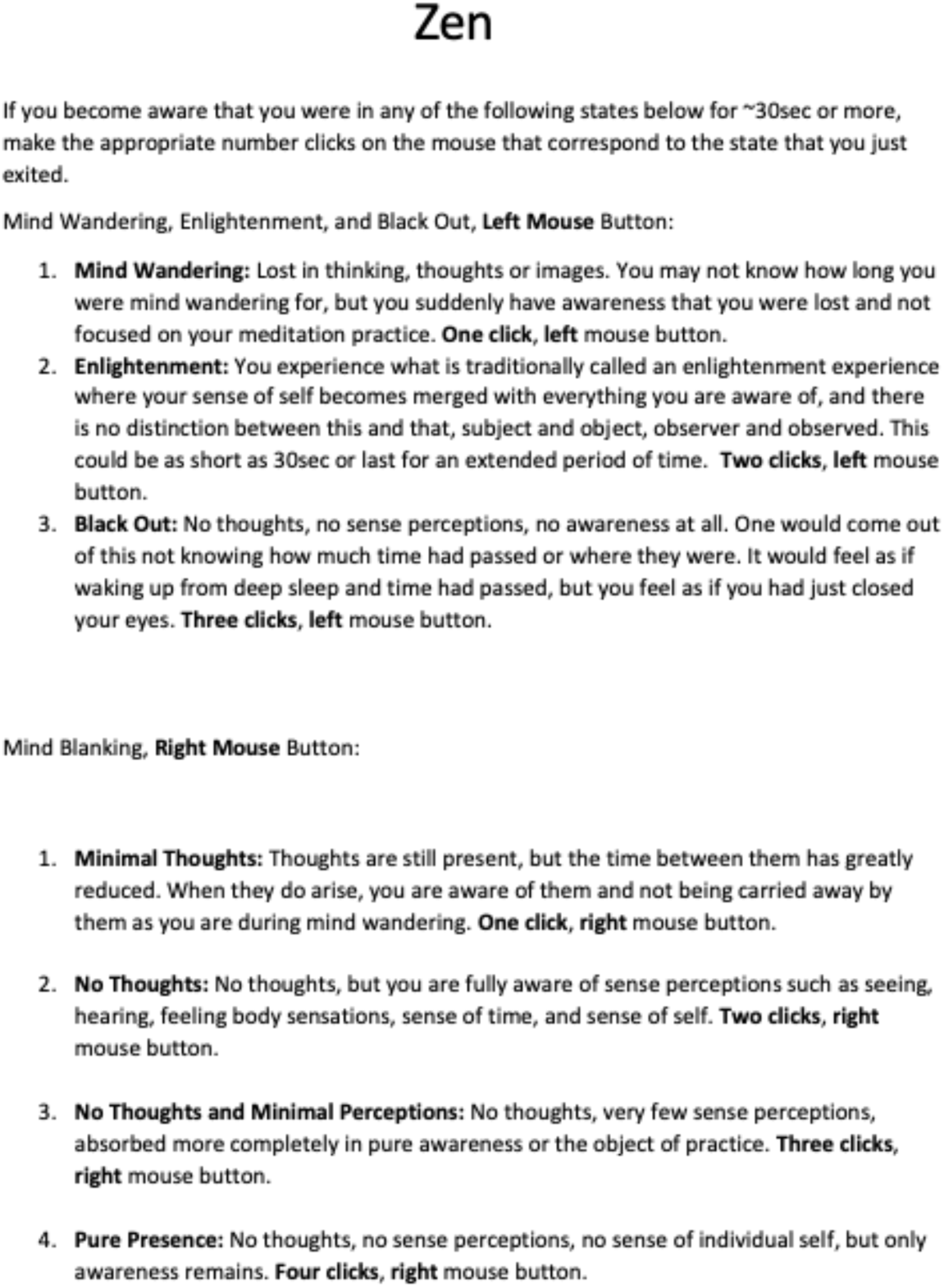

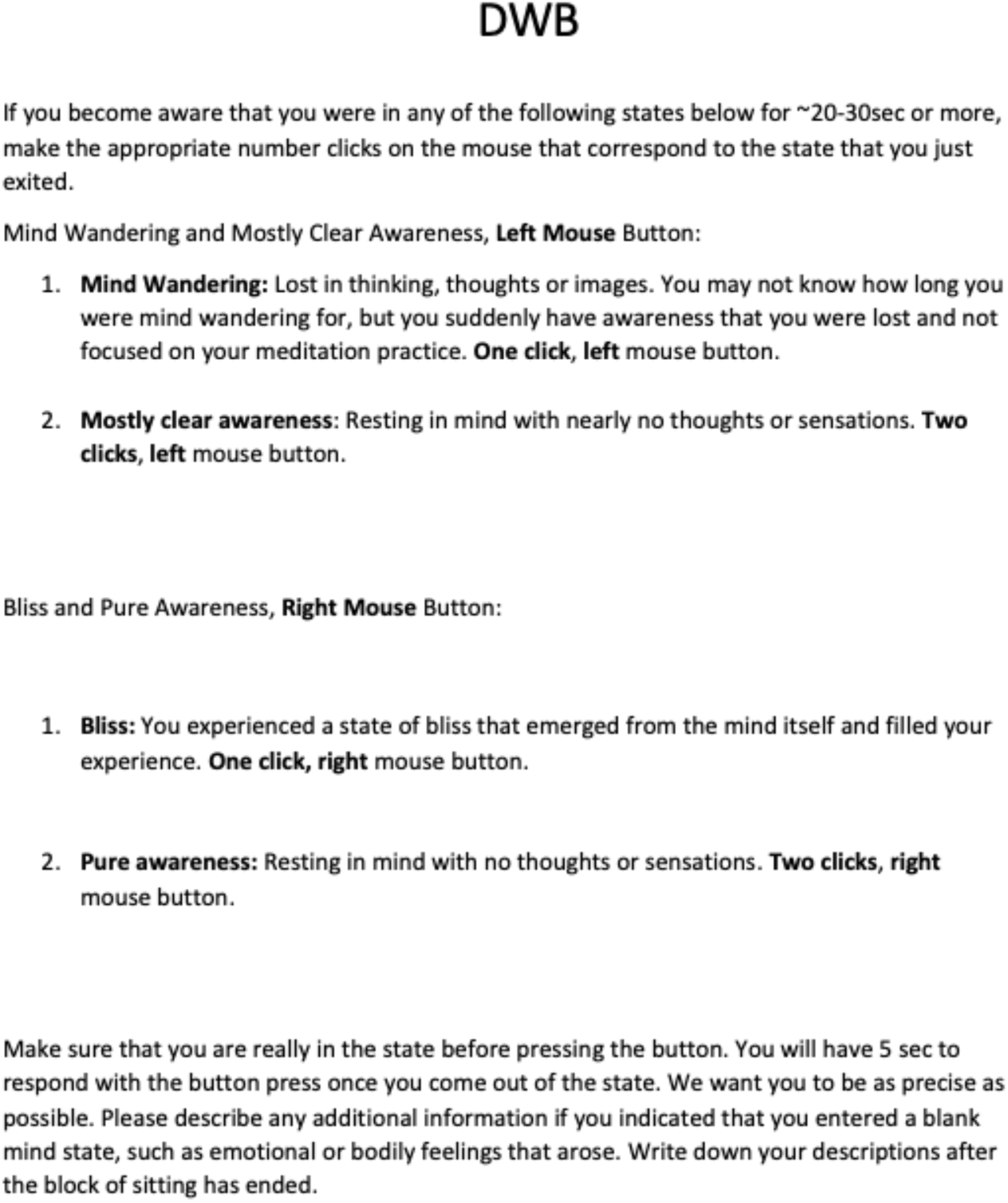

## Supplementary results: Phenomenological Interviews

The interviews followed the guidelines of the micro-phenomenological interview method (Petitmengin, 2006), which employs open-ended and iterative questioning to elicit rich, detailed accounts of specific experiential moments, including prereflective aspects, while minimizing subjective bias. They were performed by a researcher trained in microphenomenological interviews through an official one-week training course (MB). The Zen interview was conducted in a quiet and private room, and lasted 25 minutes. The Diamond Way interview was conducted by Zoom and lasted 40 minutes. Following an initial open description, each moment surrounding PP as well as PP experiences themselves were revisited in a chronological order. All interviews were audio-recorded and transcribed verbatim below.

## II. Phenomenological interview transcripts

### Phenomenological interview – DIAMOND WAY PRACTITIONER

(Practitioner was instructed to meditate just before the interview meeting, and mentioned that the dissolving phase he reached corresponded to a pure presence experience).

*[OVERVIEW OF THE SEQUENCE OF EXPERIENCES AROUND DISSOLVING PHASE – descriptions to dissolving phase/PP per se are highlighted in bold]*

<u>Interviewer:</u> I’m going to ask you to go back to the moment where you think the kind a dissolving state that was like ‘empty space’ started, basically, you know, when you want to define that start. We’re going to go back there and then we’re going to do a first pass to kind of see first maybe if you want to describe a little bit how it evolved, you know, it doesn’t need to be very long. Again, you kind of decide how long that moment is but kind of see what happened then after that in a second pass we would kind of go more a little bit in each moment if there were different moments, if that makes sense.

<u>Practitioner:</u> Yeah

<u>Interviewer:</u> Okay, so… So we’re going to start. So… Now I’m going to ask you to… go back to this moment. And, you know, when you were meditating, kind of maybe what you would consider is the start of that dissolving phase. Do you recall… basically where it was where it was, what you were doing? So where was it? Where were you located when that happened?

<u>Practitioner:</u> Sitting in my cushion meditation cushion in front of my altar.

<u>Interviewer:</u> Okay. So it was just like a few minutes ago, right?

<u>Practitioner</u>: Yes.

<u>Interviewer:</u> Okay. Okay. And so… you were sitting on your meditation cushion and – um - were you… eyes open, eyes closed, you Remember?

<u>Practitioner:</u> Yeah, I do it normally with eyes closed. If you want, I can… *[Note: this eyes-closed phase was unlike when meditating for the actual hdEEG study, where all practitioners were instructed to keep eyes open]*.

<u>Interviewer:</u> Awesome, no no. So that time we were eyes closed, okay. And tell me… What happened? What would you say happened kind of first?

<u>Practitioner:</u> So basically how the process goes is that you know – um - we call to mind an image of the 16th Karmapa and then after we finish the reciting of the mantra. I let a little moment between I finished mantra and the dissolving phase, just to keep the connection with Karmapa and then… When Karmapa melts into rainbow light and it melts into me it feels like, and then it dissolves everything around me, it feels like the process goes together with the breathing. So… I do a little breathing. When **the light comes into me, and** when **the light goes out and dissolves everything into space**, it’s a little breath out. And with this, breathing out, **this is like… letting go of some tension in the body, and through this letting go. I access into like a relaxed state of mind, but it’s not like a relaxation, it’s more like being in itself, or being itself. Mm-hmm. And then everything just settled for a little moment.**

<u>Interviewer:</u> Okay.

<u>Practitioner:</u> **There’s nothing to hold, nothing to… focus on. Nothing to consider.** Mm-hmm.

<u>Interviewer:</u> Right. And so you were on your meditation cushion and after finishing the mantra -i’m just saying it to make sure I understood that, after finishing.

<u>Practitioner:</u> Mm-hmm.

<u>Interviewer:</u> Giving it a little moment, feeling Karmapa melting to rainbow light and – um - you did a little breathe in with the light coming in, then going to breathe out when everything dissolves into space. And then after this you said there was a moment where, there was a moment everything settled. For a little moment for a little moment you were just being there or just being. And – um - and then what happened next?

<u>Practitioner:</u> Yes. I just feel like there’s nothing to hold on, nothing to be worried about, nothing to think, sometimes it’s just images. Or sounds. Actually, it’s just like… It feels like… a little cloud of smoke you know like you can see it but you can see it you cannot grasp it it’s just appears and disappears at the same time. So… Basically, it’s just being there. In this playful.. and, just…

Manifesting scenes but they really need to grasp them or anything, it’s just there.

<u>Interviewer:</u> Okay. And then… Anything more happened during that phase of the meditation? or you think then you moved on to another phase? anything more than you can recall of what happened next in that phase, or you think, transition.

<u>Practitioner:</u> So this is one moment, then I read this next: Whatever may appear as the free play of space and again.

<u>Interviewer:</u> Okay.

<u>Practitioner:</u> Like a little breathe in and breathe out, letting everything settle. It’s not forceful breathing it’s just ‘Okay, Okay’ of breathing out. Again, leaving everything. Letting everything go. Just trying to paint that relaxed state of mind of ‘being in itself’. Mm-hmm.

*[GOING DEEPER INTO EACH PHASE]*

<u>Interviewer:</u> Great. So now if you’re okay with that, we’re going to go back to kind of each kind of micro moment of what you described to me. Yeah. And kind of just… trying to recall how they felt. You know like - um - so you said that the moment you finish the mantra, and then you’d you give it a little moment - is what you said.

<u>Practitioner:</u> Mm-hmm.

<u>Interviewer:</u> And at that time, so you were sitting on your meditation cushion.. What did you feel? Can you explain to me or like describe to me this moment - very kind of beginning.

*[A. Start of dissolving phase]*

<u>Practitioner:</u> Mm-hmm. So… When I finish the mantra i just stay a little bit receiving the three lights. Basically, it’s… enjoying or… sitting in joy and receiving the three lights enjoying the connection with the Karmapa. Being infinitely.. Being thankful for.. It’s… It’s like being thankful for the blessing.. of being able to recognize my own nature, or It’s a feeling like that. It’s not easy to describe but it’s just like a feeling of thankfulness plus joy of being connected in that moment.

<u>Interviewer:</u> What did you see? Did you see anything then?

<u>Practitioner:</u> Just trying to keep the image of Karmapa in front of me and the three lights getting into my three centers. Mm-hmm. And basically. More than the image I try to focus or rest on the feeling; you know it’s more.. it’s not focusing on the feeling it’s more resting in the feeling of the joy and thankfulness for the connection.

<u>Interviewer:</u> And… Where was that feeling? Was it somewhere?

<u>Practitioner:</u> In general because.. in general, as I am receiving the three lights. I mean… You feel the three lights in the three centers. Mm-hmm. But it’s a general feeling of yeah, resting that joy. Just like…

[B. Dissolving phase per se]

<u>Interviewer:</u> Okay. And then after that, you said after that moment it was the next thing you described was… the 16 Karmapa melting into rainbow light, too.

<u>Practitioner:</u> Mm-hmm.

<u>Interviewer:</u> Describe to me be more like what how it felt.

<u>Practitioner:</u> So basically. First, I stopped feeling the three lights, so i just keep the joy and thankfulness, this feeling of joy and thankfulness is still there that is not related to the lights anymore because the lights disappear, it’s just basically being in that thankful and joyful feeling and then, the image of Kampa in front of me just starts to dissolve like if it was a… Yeah, more or less like a smoke, you know. Rainbow smoke. Rainbow light smoke like moving you know like the aNuma more or less image. And then… Travels. It goes… like on onto the top of my head so it falls into me like from top goes into the central channel and then it starts to go everywhere. In my body. And then this is when I breathe in small breathing. When everything comes in. Mm-hmm. And **when I breathe out just like the light goes out from me. Yeah. Touches immediately everything and everything just disappears.**

<u>Interviewer:</u> And at that moment when everything disappears, **do you see something?**

<u>Practitioner:</u> **No, it’s just… a moment.**

<u>Interviewer:</u> How would you describe the breath?

<u>Practitioner:</u> Breathing out, all the tension in my body like the feeling of a stiff back and hands. Arms. Just relax, like completely ghost and I feel my body settling in, and with this feeling of the body settling in also like the mind. And **for a moment, there’s nothing. And the feeling of space and this feeling of joy and thankfulness is still there, you know? Then for a moment, it’s just being there… but… relax. And relax as you know like there’s nothing to hold on, nothing to keep, everything is true. Nothing to do. Nothing to feel it’s just… Everything just goes. Yeah, you’re like just settled in the place.** And even though you start breathing again this is the moment when the brain goes out under breath if you’re out of breath. And then slowly you start to breathe again and breathe normally. This feeling doesn’t, feeling doesn’t disturb the feeling of being there. It’s just… part of everything.

<u>Interviewer:</u> So you said at some point you would… like you had the breathing out, we’re kind of being out of breath and then breathing we start again

<u>Practitioner:</u> Yeah, but it’s not like forced, you know, I don’t try to keep the breath out as much as long as I can hold, it’s just… Without… **certainly in the feeling of space, and mainly it’s just feeling of space.**

<u>Interviewer:</u> Okay.

<u>Practitioner:</u> And then naturally you start to breathe But… doesn’t disturb this feeling of being in itself being in this space.

<u>Interviewer:</u> Okay. And **anything… else does that come to your mind to describe?**

<u>Practitioner:</u> **Space**.

<u>Interviewer:</u> Right.

<u>Practitioner:</u> Mm-hmm.

<u>Interviewer:</u> Um… And so… After that moment, when you said you were just being there.

That your breath was coming back but not effortfully and you were feeling it, but it was not disturbing it. It was still part of it.. And you had that feeling of that space, right?

<u>Practitioner:</u> Yes. Yes.

[C. End of dissolving phase]

<u>Interviewer:</u> Is that a good description of it? So now, you were on your meditation cushion, and then you had, after your breathing comes back and you had that feeling of space, and just being there. What happens next? Just after that, what did you feel?

<u>Practitioner:</u> It’s not like… I don’t hear anything. It’s not like i don’t hear anything I don’t have… the sensation of the body. Or that I don’t have any visual sensation. It is more that all these sensations are like.. Like bubbles maybe you know like you know they are there but there are no essence. No.. No need fix on the task. You see them like a bubble in the sky and then it just pops up and then they.. they disappear… sounds and feelings. Then these start to become more and more present.. Necessarily in the moment. The mind is like cannot.. cannot resist and starts to relate to those experiences. And that’s when I know that when moment is gone. Mm-hmm.

<u>Interviewer:</u> And when you say the mind cannot resist and starts to relate to these experiences what do you feel? Can you describe to me, how.. how is it?

<u>Practitioner:</u> It’s like… You know, to be in that state of openness and space requires.. It’s not effortful. But it requires some kind of balance. But the mind can just be open, and when the breath goes out, it’s a moment like everything is cleaned, you know, all the information, sensation content of the mind is clear for a brief moment. These sensations and ideas and things and images and everything starts to come back again slowly like i told you, yeah, like kind of bubbles.. Like soap bubbles and then they pop, and then they pop, when they are few. It’s easier just to… let them pop and not get involved. But the more time passes, more of these bubbles start to come up. And the more there are the more difficult is to keep just completely open because the mind open somehow has the need, or has this drive.. to relate to them.

<u>Interviewer:</u> And how do you know that your mind starts to relate to them or get involved? How’s the difference in feeling, let’s say.. How do you know that your mind gets involved or starts to relate to things?

<u>Practitioner:</u> Because… The moment the mind relates to one of those, that becomes the dominant thing in the mind. You know, it’s like there are so many sounds, there are many sounds and feelings like the body and images popping in, but in the moment the mind relates to one of them, that is the dominant sensation that starts to.. develop.

<u>Interviewer</u>: Okay.

<u>Practitioner:</u> Strain.. i don’t know if it’s the word is strain or train.. like the train of thoughts and concepts that comes afterwards.. yeah it’s like you have several like this popping up like.. and it’s like your attention is completely diffused or completely open, and it’s not focused on anything. It’s just and then all this starts to become more and more sensations.. And there was one moment in which when the mind relates to one of those is.. is like the attention is fixed on that. And when the attention is fixed on that it becomes the dominant thing in your mind and then all the concepts and dominant and inner narrative and you know inner dialogue starts to go after that.. And that’s the way when I realized okay, the freshness of the moment is gone. And then… as the training in the 8^th^ Karmapa, the idea is that you can train to go back to the place through the breath. The dissolving phase moment. And you can go back to that moment while you’re in the meditation. And the whole process starts again.

<u>Interviewer</u>: So you said to me so you feel the difference when your mind starts to relate because it’s like now one sensation becomes dominant and start like a train of thoughts and concepts.

<u>Practitioner</u>: Mm-hmm.

<u>Interviewer:</u> Do you… feel anything different, thing or sensation in your body, do you feel something different when that happened, when that just happened to you, when you meditated, there on the cushion then you had this kind of sensations, starting to bubble or pop one by one and then.. And then when there’s that kind of moment where your mind starts to relate and starts to have that dominant sensation in a train of thoughts. How does it feel like if you had to kind of think, like, do you feel anything different in your body or like or like your vision, like any senses, anything else to describe that sensation?

<u>Practitioner:</u> Somehow that, you mentioned, it’s like .. Before everything is there you know like sensations of the body, the sounds everything is just present. It’s part of the experience, it’s not, it’s not like it’s a black hole, yeah, it’s like in the moment when everything dissolves. It’s like all everything is there. But the moment that the moment something becomes dominant, everything else is filtered out. Like for example if I, if I hear a sound and that sound remind me something then that’s on, my mind relates to that sound, I start to think about the sound and the feelings of the body disappear or become less, become filtered, they are filtered, uh-huh..

Filtered out. And then the dominant thing just of course the dominant thing for some moment is… It’s that which captures the mind. But then… all the feelings start or other things start to pop up and then the mind jumps to those other things. And that’s when you know, that’s, that’s… Because it might be that.. it might be sometime, you know, when, when the meditation is not that fresh or that it has not helped me go to the dissolving state, these moments of mind getting related to things are more frequent. But somehow it’s easier to let them go if you capture it in the in the beginning, yeah. You just breathe out and it goes away again.

<u>Interviewer:</u> Okay.

<u>Practitioner:</u> The moment where the freshness is gone is like the mind becomes related to one thing and then jump to the next thing and then jump to the next thing and it can be, you know, like starts with the sound and then go to the feeling of the body that the knee is hurting and then goes to an image that just pop in.. And then it’s just… Yeah, it’s just gone. And that’s when… I, like, I go to ending part of the meditation with Lama describes how the world appears., perfect and pure. Everything … every sound is mantra and every thought is wisdom.

<u>Interviewer:</u> Yeah.

<u>Practitioner:</u> You know the description that helps me to, helped me.. Or I use it as a guideline to, guideline to give direction to, direction to the sensations and, the sensations and experiences that are popping into mind. So it helped me to give direction to that kind train of… conceptual thinking or thinking narrative that is starting to happen, that it goes through that idea that everything is perfect, everything is.. to everybody’s Buddha, whether they know it or not and Yeah, it goes there. Mm-hmm.

<u>Interviewer:</u> Okay. So if I… rephrase this, you said that… In 8th Kamapa training sometimes when you notice, sometimes when you notice sensation popping, then you can actually go back to it.

<u>Practitioner:</u> Mm-hmm.

<u>Interviewer:</u> ..to go back to the place. Kind of breathing out again and but it depends on times, yeah, but um, but otherwise if when… you are kind of done with the moment when the moment ends, and your mind starts to relate to do things, then you actually, that’s kind of when you decide when the phase of the meditation is ended and then you go to the next phase, one where the world apprears. That’s what you said, right?

<u>Practitioner:</u> Mm-hmm. Mm-hmm.

<u>Interviewer:</u> Yeah. And there you said that the… streams of sensations continues but then you actually use the meditation instructions as a guide.. ‘I had to give them direction’. That’s what you said, yeah?

<u>Practitioner</u>: Mm-hmm.

[D. World reappears]

<u>Interviewer:</u> And so in that phase, just to kind of give you the contrast, when you start that phase of meditation. So just try to remember, when you were this, on your cushion like a few minutes ago, when you did that part. This meditation phase with the instructions then the word reappears perfect and pure. Can you describe maybe more like, what you.. Did you see anything? Did you feel anything in your body?

<u>Practitioner:</u> Um… Let me see.

<u>Interviewer:</u> What happened when you just you did this instruction the world reappears perfect and pure. So what did you feel, or what did you see, what happened then?

<u>Practitioner:</u> Yeah, the moment, the sensations of the body becomes more solid. It’s not just something that it’s present there, but it’s something that has become something, a body. You know, it’s like when the sensations are there and there is no point of reference point, even though I know it’s my body. It’s just the sensation. There is no relations created after the sensation. But when the world reappear, when there’s no point of reference, that’s when you were in the previous phase like.. And now then the road appears. Yes, exactly. When the world appears, then the sensations have a point of reference that is the body, and then it becomes solid. And then when it becomes solid I start to see again, view the pain or the feeling that I have there in my shoulders and my back. And then… the cushion, the feeling of being somewhere like sitting somewhere Um.

<u>Interviewer:</u> Do you see images in your mind? Do you see anything?

<u>Practitioner:</u> Even though… I think not really. It’s just more the first thing is the body. It starts to get more solid. And then… So when the… instruction says like, um, Every sounds are mantras and every thoughts are wisdom, whether you know it or not. Then sounds become solid also like, you know, that something is there because someone is outside talking. You know, like the reference of those sensations become solid somehow, and then, um, but with the instruction that we get from this last part of the meditation when it solidifies the point the reference of the of the, the reference source of that of the sensation. When they start with this reference source it starts to become solid then you can solidify them with this pure view. Concept you know like okay, these mantras. The sounds something pure. And also because, you know, when I meditate in my house there’s a meditation also there’s like a little park, a park, there are always people there and.. The first thing I hear, and I say to you the sound is there before and it doesn’t have any disturbance because there is no reference, so the sounds just disappear, the sensation, it doesn’t have any meaning. It doesn’t have any value. But when the world starts to appear then the sensation starts to get the value, and then the value solidifies into someone you know like uh, there are some people. But then these people with instruction sounds much more how you see the instructions, now beings manifest near and far they are female or male buddhas whether they know it or not. When… when this sensation becomes solidified in a person then in a person you can attach to that concept the understanding that they have Buddha nature. And therefore they are Buddhas whether they know it or not. So it’s like when it’s defined. If you attach the instruction to this process then… the becoming of those sensations instead of become something normal it becomes the pure view experience.

<u>Interviewer:</u> Thank you. Anything else you want to… in general?

<u>Practitioner</u>: No.

<u>Interviewer</u>: Wonderful.

<u>Practitioner:</u> Yep.

## II. Phenomenological interview – Zen practitioner

*[This interview refers to the PP experience during hdEEG recording. Descriptions to PP per se are highlighted in bold]*.

<u>Interviewer:</u> I’m going to invite you to think about a moment of your choice - maybe in a recent meditation retreat or whenever you think is appropriate - that you would like to describe as what would correspond to like a good example of pure presence. You can think a moment and let me know when I think you found one.

<u>Practitioner</u>: Okay.

<u>Interviewer</u>: So… that moment that we’re going back together, where was it?

<u>Practitioner:</u> the Madison Zen Center. In the Zendo. Sitting on a cushion.

<u>Interviewer</u>: Okay.

<u>Practitioner:</u> Um.. in the corner opposite the altar.

<u>Interviewer:</u> Okay. And… So when you start at this moment, what did you have in front of you? it was the altar?

<u>Practitioner</u>: it would have been… kind of a screen, just like a black screen.

<u>Interviewer:</u> And… Anything else you remember? That day of time. Just before it started. You were meditating on a cushion then?

<u>Practitioner</u>: Yeah. Yeah.

<u>Interviewer:</u> Okay. And so… The first thing you remember of that moment: what would you say?

<u>Practitioner:</u> So right before the moment… just thinking like the transition a few seconds before, a certain intensity, very awake. Very absorbed and concentrated. Not much. There not really seeing much. Maybe right before, maybe they’re still a little small sensation in the body, but not a whole lot. Everything feels very quiet. My spine feels very straight. I said, very awake.

<u>Interviewer:</u> And then what happened? First, we’re going to unroll it a bit and then come back.

<u>Practitioner:</u> Yeah. Then my breathing is rather slow. Almost like longer pauses in between my inhalation and exhalations. And then… It’s like.. what I usually would feel is my body being in space. It’s like my body starts to fill all of the space. So there’s not the image of the body, but just the awareness itself. Kind of fills the space of the space - I guess the room - or at that point it’s not really the room, it’s just It’s just like everything is just like everything. Vast spaciousness. But without necessarily a center.

<u>Interviewer</u>: Okay.

<u>Practitioner:</u> And… Definitely there’s a little, I wouldn’t say it’s a feeling, but there’s definitely like a - um - a sense of intimacy and I wouldn’t say bliss, but certainly, lack of desire. Definitely lack of grasping. Pleasant would be the way to describe it, but it’s it’s more the - um - not the normal pleasant of like being in the body and feeling pleasant it’s more sense of freedom, I guess, maybe. You can call it. Being completely unencumbered by anything. Complete ease and rest at the same time, but also having a lot of energy. But very peaceful, calm energy even though intense. I guess intense maybe for like - I don’t know how to describe it but – um - okay, almost like a - like you could hear a pin drop like a like a cavern - i don’t know - cavernous spaciousness. There’s… There’s no colors or anything. Like the absence of all images so it’s just there’s no forward or back. Not really an up or down. I guess a lot, there’s a lot of vibrant - I don’t know, pulsing might not be the word but – um - vibrant energy - that’s not separate from anything else. It’s just kind of is separate vibrant energy being - i suppose that’s aware and awake in sensing. I’m definitely aware. Without location, without… a sense of time. Without really a sense of place. And it kind of feels like maybe it stretches forever.

<u>Interviewer:</u> Okay. And… how do you know the… like what do you feel next, when the moment kind of stops and this state kind of changes.

<u>Practitioner:</u> Let’s say maybe the first thing I become aware of is my breathing. Coming out of it.

Yeah, probably my breathing. And then… maybe sense of body a little bit. At this point, I remember… maybe feeling the mouse in my hand. That I was using to indicate.. different states.

And then at some point just… Not so much thoughts, but… Certainly concepts like, you know, the concept of the body and the mouse. But at this point, not really thoughts yet. It’s more - um - more general concepts of things. Maybe hearing. Sense of hearing. And then a certain remembering of: okay, I need to click. On the mouse. Feeling that and then probably sometime after that thoughts just start to you know like reflection, self-reflection or reflection on – um - the state itself. Still feeling - um - very energized and clear. And… Very, I don’t know… Like definitely lack of wanting.

<u>Interviewer:</u> Okay. Yeah. And at that moment, you kind of know that the experience ended.

<u>Practitioner:</u> Yeah, but it’s still, it ended, but there’s still like that after-effect, afterglow. I don’t know what to call it.. Like the taste of it that’s still there.

<u>Interviewer:</u> So right so.. Let me just rephrase, kind of a little summary of what you said and we might go back to one moment. So you said that just before the experience you had was a few seconds where you were very awake.

<u>Practitioner:</u> Yeah. And but also like absorbed, concentrated.

<u>Interviewer:</u> And now breathing slow.

<u>Practitioner:</u> Yeah. And then…

<u>Interviewer:</u> So that’s kind of just before it starts before it starts. You wanted to say more about?

<u>Practitioner:</u> Yeah, I would say the um my vision kind of goes black. So eyes are slightly open, but I don’t see anything. Not really hearing a whole lot, I still could hear if I needed to, but it’s like everything’s there’s like a silence that kind of starts to be more present like just because i’m so absorbed in the awareness itself, other things just start to fall away.

<u>Interviewer:</u> And that awareness itself

<u>Practitioner:</u> Yeah

<u>Interviewer:</u> like where is it, or - um - does they have a shape or anything else to describe?

<u>Practitioner:</u> It’s more, you know, my self-conscious awareness that I would use to say place my awareness on an object. But instead of having objects, it’s just kind of like um taking the awareness that’s internal and placing it on the awareness that’s external. That’s just and at that point, my body tends to feel very empty. So it’s not really located anywhere. Yeah, it’s more of a merging of kind of the inner and outer awareness, I guess.

<u>Interviewer:</u> And then you said your breathing slows?

<u>Practitioner:</u> Yeah, slows down and my pauses between inhalation and exhalation start to maybe increase a little bit. So there’s moments where I just have a little bit of breath suspension. Just naturally without, you know, I’m not trying to control my breathing at this point. It’s just noticing that my breathing is getting slower and - um - probably deeper. In the transition between inhalation and exhalation, especially after I inhale the exhale sometimes takes, there’s like a pause between the inhalation and the exhalation. And at that point and where that pauses, my concentration feels even heightened. It’s like at that moment, the concentration gets more acute or more… focused, which seems to be the thing that happens right before I would get into this state of everything kind of falling away.

<u>Interviewer:</u> And then when you say the feeling of everything falling away from you..

<u>Practitioner:</u> Yeah, it’s – um - kind of the body, like not feeling my body in space. Kind of like a just kind of like everything merges with this just broad spacious awareness. It’s like there’s no markers between inside and outside. Those disappear and it’s just all one basically one, just one. Awareness with no nothing in between. And time, you know, I know at that point, I had no idea how much time is passing.

<u>Interviewer:</u> Do you think time.. do you feel time passing?

<u>Practitioner:</u> Not really. I mean, when I come out of it, I might be able to guess for how long. I think I could retrospectively look back and maybe have some idea but – um - in the moment I typically don’t.

<u>Interviewer</u>: And when you feel like this, with your body there is no inside, no outside.

<u>Practitioner:</u> Yeah.

<u>Interviewer:</u> Do you see anything or hear anything?

<u>Practitioner:</u> Don’t really see anything. I guess hearing be it’s hard to describe. It’s like hearing silence.

<u>Interviewer</u>: You had said something about cavernous spaciousness.

<u>Practitioner:</u> Yeah, I mean, it would be like seeing emptiness. And hearing silence. That’s probably… Maybe the best way I could describe it.

<u>Interviewer:</u> Okay. Do you feel anything else?

<u>Practitioner:</u> Not in my body, but it’s just a feeling of intimacy. So like deep connection, intimacy, non-separation - which, you know, feels good - like - it’s, it’s like a… I mean, you know, I mean, it would be very close to like love, I guess, in some ways. Like not you know that type of non-separation, so it’s very… close, like it’s very close. And then afterwards, so say there’d be like a sensation of say looking at an object and looking at an object still then feeling some separation, but feeling so much closer. Everything feels – in Zen, we call it a term called suchness or thusness. Like a… close, intimate, almost things are more alive, like more alive or a certain sparkle, for lack of a better word. Very close. Everything feels very, very…

<u>Interviewer</u>: That’s like after when you…

<u>Practitioner</u>: Yeah, that’s after.

<u>Interviewer</u>: And what would you say so you said the first thing when you come out of it, Is there breathing?

<u>Practitioner:</u> Breathing. Comes back. Breathing comes back and then maybe feeling my body on the mat would be second. Sense of time, probably, to a certain degree - there’s a, you know, not thinking about time but I would say with the coming back of the body and the breathing, more awareness of time.

<u>Interviewer</u>: And then…

<u>Practitioner:</u> Kind of awareness of, of the reporting or just being aware that I’m in the zendo. Meditating.

<u>Interviewer</u>: Okay.

<u>Practitioner:</u> Then maybe… Hearing. At some point I think hearing; I’m not sure if hearing kind of comes back after before the body, but I’m thinking after. And it’s a subtle transition. It’s not dramatic. So it’s not like feeling like I’m suddenly dropped into the body it’s more - um - much more gradual, it’s much more soft transition. Right. And there’s a feeling of awe. A little bit of feeling of awe. I would say…Like hard… on the sense that it’s a bit hard to describe that state. If I start to think about it. Being – um - always harder to describe, yeah. Yeah, I think because it’s kind of hard to describe, there’s a feeling kind of awe or wonder, I would say one, like a feeling of wonder, like.

<u>Interviewer</u>: Is there a location to that?

<u>Practitioner:</u> No, not really. Not so much. It’s more of a conception. It’s more of a Okay.

<u>Interviewer</u>: You had like this gesture (mimicking practitioner opening both arms and hands)

<u>Practitioner:</u> Yeah, yeah, I mean, when I… At some point the transition is like this and everything’s just like empty space. Right. And then at some point everything kind of starts to come back. But I do feel more transparent after this. So my body feels less concrete. And… Probably my heart area. This whole area feels a lot more open, expansive. And so there’s, so there’s, here’s an after-effect. It’s still kind of silent. There’s still a silence within hearing noise like there’s a quiet that continues afterwards. And it’s not desire, like I have no desire to like, go back to that necessarily at all. It’s just…

<u>Interviewer:</u> Yeah. Anything else you wanted to share about this experience of this moment?

<u>Practitioner:</u> To a certain degree, when it happens is not always under my control, meaning I’ll be meditating and I just I just am focusing on, you know. We call Shikantaza, awareness itself. And I’m not trying to grasp onto anything or reach a certain state. When I’m really concentrated, just kind of will happen, just something, something kind of switches.

<u>Interviewer:</u> So.. Thank you very much for sharing.

<u>Practitioner</u>: Yeah, it really feels sharing the experience brings a lot of the feelings back.

<u>Interviewer</u>: Great for everyone then.

<u>Practitioner</u>: Yeah, exactly.

<u>Interviewer:</u> Perfect. Well, thank you.

